# Monomer binding modes of small molecules that modulate the kinetics of hIAPP amyloid formation

**DOI:** 10.1101/2025.09.22.677832

**Authors:** Michelle Garcia, Korey M. Reid, Paul Robustelli

## Abstract

Human islet amyloid polypeptide (hIAPP) forms amyloid fibrils that accumulate in pancreatic β-cells of Type II Diabetes (T2D) patients. Recently discovered small molecules that modulate the kinetics of hIAPP amyloid formation could serve as starting points for developing T2D therapeutics, but no structural or mechanistic rationale exists to explain their binding mechanisms or effects on hIAPP aggregation pathways. Here, we utilize all-atom molecular dynamics computer simulations to elucidate the binding mechanisms of an hIAPP aggregation inhibitor (YX-I-1) and an aggregation accelerator (YX-A-1) to disordered monomers of wild-type hIAPP and the naturally occurring pathogenic S20G hIAPP variant associated with early-onset T2D. We observe that the inhibitor exhibits substantially higher affinity for monomeric wild-type hIAPP than the accelerator, consistent with previously reported biophysical experiments. We dissect the interactions that stabilize binding of each molecule to wild-type and S20G hIAPP and characterize conformational changes that occur upon ligand binding. In all ligand-bound ensembles, distinct fragments of YX-I-1 and YX-A-1 are sequestered from solvent upon binding while other fragments remain solvent-exposed. Based on our simulations we hypothesize that buried ligand moieties confer hIAPP monomer binding affinity while solvent-exposed ligand moieties modulate the kinetics of intermolecular association of bound hIAPP into higher-order oligomeric intermediates on amyloid aggregation pathways.

## Introduction

Amyloid diseases are characterized by the misfolding of proteins into toxic oligomeric aggregates and highly structured β-sheet rich fibrils^1^. Over 50 diseases of this class exist including Parkinson’s, Alzheimer’s Disease (AD), and Type II Diabetes (T2D)^1–3^. While many amyloid diseases are age-related, T2D is not. T2D has reached epidemic levels as the most common metabolic disease^4^. Patient symptoms such as insulin resistance, pancreatic β-cell dysfunction and β-cell death are associated with the formation of amyloid fibrils of human islet amyloid polypeptide (hIAPP or “amylin”), a 37-residue glucose-regulating hormone co-secreted with insulin^5–11^. The self-assembly of soluble cytotoxic oligomeric intermediates, rather than insoluble amyloid plaques, is hypothesized to be the primary driver of β-cell dysfunction and T2D pathogenesis^12–16^. Through receptor-mediated and membrane interactions, these oligomeric species may increase levels of reactive oxygen species, induce apoptosis and overload the protein quality control machinery^13^. In some Chinese and Japanese populations, patients with a Ser20-to-Gly (S20G) mutation in hIAPP−the only known naturally occurring hIAPP variant−have an increased predisposition for developing early-onset T2D^10,11,17,18^. The S20G hIAPP variant exhibits accelerated aggregation kinetics^19,20^ and increased intracellular cytotoxicity^21^ compared to wild-type (WT) hIAPP.

Numerous fibrillation studies using hIAPP peptide variants have provided insight into its aggregation pathways of hIAPP^22,23^, but the molecular mechanisms of hIAPP aggregation remain poorly understood at atomic resolution. In solution, hIAPP is an intrinsically disordered protein (IDP) in its monomeric form, adopting a heterogeneous ensemble of interconverting conformations under physiological conditions^1,14,24,25^. Due to the dynamic nature of hIAPP monomers, it is challenging to identify specific conformational states associated with aggregation pathways^1,14^. In general, the kinetics of amyloid formation follows a two-stage nucleation-growth mechanism consisting of a lag phase followed by an elongation/growth phase^3,14^. In the lag-phase, monomers aggregate into higher order oligomers via primary nucleation. The conformational properties of monomeric and oligomeric hIAPP states that drive primary nucleation remain poorly understood, and multiple hypotheses have been proposed. One hypothesis suggests that the transition states of primary nucleation may involve disordered assemblies in which random coil regions of multiple hIAPP monomers coalesce to form protofilaments^14,26,27^. An alternative hypothesis posits that nucleation intermediates are ordered oligomers in which hairpin motifs of multiple monomers convert to the fibrilization-prone β-sheets^28–31^. Fibril growth kinetics experiments have shown that once the concentration and length of fibrils reaches a critical threshold, secondary nucleation on the surface of existing fibrils becomes the dominant pathway for fibril formation in the growth phase of hIAPP^14,15,20^.

Therapeutic interventions have been proposed at all stages of fibril-forming pathways. Recent advances in cryo-EM techniques have produced structures of numerous polymorphs of mature hIAPP fibrils^32–36^, providing insight into fibril-forming regions and protofilament interfaces. These structures and additional *in vitro* studies frequently distinguish three regions the hIAPP sequence: the N-terminus (residues 1-19), which is involved in membrane interactions, the amyloid core (residues 20-29), and the C-terminus (residues 30-37), which are primarily involved in protein-protein interactions^37,38^. Some inhibitor design strategies have targeted the amyloid core, aiming to disrupt fibril elongation by targeting buried hydrophobic residues or by sterically capping fibril ends^38,39^. A wide range of peptide and peptidomimetic inhibitors have been developed using diverse strategies including incorporating amyloid core-derived fragments, proline or charged residue substitutions, N-methylated backbones, β-sheet breakers, cyclic or stapled motifs and non-natural amino acids^38,40^. Peptide segments derived from the biologically potent native inhibitor insulin have also been shown to inhibit monomeric forms of hIAPP^38,40,41^. Small molecule screens^42,43^ and in vitro studies of polyphenols have identified several weak inhibitors of aggregation with diverse chemical structures^38,42,44–47^. MD simulations of hIAPP with several polyphenol inhibitors suggest that binding modes are largely stabilized by hydrophobic and π–stacking interactions with aromatic residues (F15, H18, F23, Y37), which are hypothesized to prevent self-association^40,47–49^. The chemical properties of polyphenol and peptide hIAPP inhibitors have generally been found to exhibit poor stability^50^, limited selectivity^46^, and variable bioavailability^46,50^. This has generated interest in the development of more drug-like small molecule inhibitors of hIAPP as potential therapeutics for T2D.

In recent work by Radford and coworkers, a combination of thioflavin T (ThT) kinetic assays and native electrospray ionization mass-spectrometry (nESI-MS) was used to screen a library of 1500 small molecules to identify ligands that alter the aggregation kinetics of hIAPP^20^. This screen identified the small molecule YX-I-1, which delays the aggregation of wild-type (WT) hIAPP but has no effect on the aggregation kinetics of the S20G variant, and the small molecule YX-A-1, which accelerates the rate of aggregation of both WT and S20G hIAPP at substoichiometric concentrations. Kinetic analysis, native mass-spectrometry, surface plasmon resonance and nuclear magnetic resonance (NMR) experiments revealed that YX-I-1 directly binds WT hIAPP monomers and delays primary nucleation, secondary nucleation and elongation of WT hIAPP. YX-A-1 displayed minimal interaction with monomeric WT and S20G hIAPP, suggesting that it likely interacts with minorly populated higher-order species formed by both species in the lag phase of amyloid misfolding pathways. Both compounds were shown to be specific in their interactions with hIAPP, showing no interaction or effects on the aggregation kinetics of Aβ42, which has approximately 50% sequence similarity to hIAPP^20^.

Subsequent work by Taylor, Radford and coworkers^36^ used YX-I-1 as a lead compound to identify structurally related, regulator-approved drugs with similar structures and chemical scaffolds. Experimental screens of these compounds identified additional ligands that bind monomeric hIAPP, inhibit its aggregation, and induce substantial changes in hIAPP fibril morphology^36^. These results demonstrate that YX-I-1 and related compounds may represent promising leads for the development of T2D drugs. The hIAPP binding mechanisms of YX-I-1, YX-A-1 and related compounds, however, remain poorly understood in atomic detail.

Identifying the molecular basis of the interactions of YX-I-1 and YX-A-1 with WT and S20G hIAPP could provide insight into the aggregation pathways of hIAPP and guide the development of hIAPP inhibitors with therapeutic potential for T2D patients. To this end, we performed enhanced sampling all-atom explicit-solvent molecular dynamics (MD) computer simulations of WT and S20G hIAPP in the presence and absence of YX-I-1 and YX-A-1 and analyzed these simulations using a recently developed circuit topology approach^51^. We compared the apo WT hIAPP ensemble to previously reported NMR chemical shifts^25^ and found excellent agreement, demonstrating that our simulation approach provides an accurate description of the conformational properties of this system. In simulations of apo S20G hIAPP, we observed that the S20G mutation introduces a central hinge that increases the populations of intramolecular interactions and β-sheets formed by residues ^7^CATQRLANFLV^17^ and ^23^NNFGAIL^27^ (which are located within the hIAPP amyloid core), relative to WT hIAPP.

Enhanced sampling MD simulations of monomeric WT and S20G hIAPP in the presence of YX-I-1 and YX-A-1 revealed that each molecule samples a highly heterogeneous ensemble of distinct binding modes. We characterized changes in the conformational ensembles of WT and S20G hIAPP that occur upon ligand binding and dissected the intermolecular interactions that stabilize the binding modes of each compound. We observed that WT hIAPP has a substantially higher affinity for YX-I-1 than YX-A-1 and that YX-I-1 produces larger changes in the conformational ensemble of WT hIAPP upon binding than YX-A-1, in agreement with previously reported NMR chemical shift perturbations^20^. We found that both ligands interact with monomeric hIAPP via hydrophobic and aromatic interactions, but YX-I-1 has a substantially larger population of hydrogen bonding interactions with hIAPP relative to YX-A-1. We found that the simulated binding affinity of YX-I-1 to S20G hIAPP was comparable to the simulated affinity of YX-I-1 to WT hIAPP, in contrast to NMR chemical shift perturbation data and other biophysical measurements which indicate weaker binding of YX-I-1 to S20G hIAPP. However, we observed that YX-I-1 binding induced larger changes in the populations of secondary structure elements of WT hIAPP relative to S20G hIAPP, and that YX-I-1 binding modes to WT hIAPP more frequently engaged multiple distinct regions of hIAPP.

Intriguingly, we observed that in ligand-binding simulations distinct fragments of YX-I-1 and YX-A-1 are consistently buried and sequestered from solvent upon binding hIAPP while specific chemical moieties of each molecule consistently remain solvent exposed across binding modes. Based on our simulations, we hypothesize that distinct molecular moieties confer binding affinity to monomeric hIAPP, while solvent-exposed moieties may predominantly affect rates of intermolecular association into higher order species. Taken together, the atomically detailed binding mechanisms of YX-I-1 and YX-A-1 reported in this work suggest strategies for designing additional ligands to modulate the aggregation propensity of hIAPP and may facilitate the rational development of improved hIAPP aggregation inhibitors and potential T2D therapeutics.

## Results

### Comparing conformational ensembles of wild-type and S20G hIAPP

We performed unbiased, all-atom, explicit solvent simulations of apo wild-type (WT) hIAPP and S20G hIAPP with the replica exchange solute tempering (REST2) enhanced sampling algorithm^52–54^. REST2 simulations were run with 20 replicas using a solute temperature ladder spanning 300-500 K. hIAPP and water molecules were parameterized with the a99SB*-disp* protein force field and a99SB*-disp* water model^55^. Apo simulations of WT and S20G hIAPP monomers were run for 4.3 µs/replica and 3.8 µs/replica, with aggregate simulation times of 86 µs and 76 µs, respectively. We evaluated the convergence of REST2 simulations by assessing the time evolution and statistical error estimates of secondary structure propensities, intramolecular contact probabilities, and the α-helical order parameter Sα^56^ in each temperature replica and independent demultiplexed replica (Table 1, Figures S1-S3). We observed a slow decay of initial Sα values in the 300 K replicas of both simulations and therefore discarded the first 1μs of sampling in both trajectories as an equilibration period (Figure S3). The following analyses are reported for the final 3.3 and 2.8 µs of the 300 K temperature replica of the apo WT hIAPP and S20G hIAPP simulations, respectively.

**Table 1.** Conformational properties of apo and holo hIAPP ensembles from REST2 MD simulations. Average values and statistical error estimates of the Cα radius of gyration (R_g_), helix fraction, β-sheet fraction, and the Sα helical order parameter from apo and ligand-binding simulations of WT and S20G hIAPP with YX-I-1 and YX-A-1. Values are reported for the 300K replica of REST2 MD simulations. Statistical error estimates were computed using a blocking analysis.

| System | $R_g$ (nm) | Helix<br>Fraction (%) | β-Sheet<br>Fraction (%) | $S\alpha$ |
| --- | --- | --- | --- | --- |
| WT hIAPP (apo) | $1.30 \pm 0.01$ | $12.4 \pm 0.4$ | $4.9 \pm 0.4$ | $1.69 \pm 0.09$ |
| WT hIAPP (apo)<br>NMR reweighted | $1.35 \pm 0.01$ | $11.0 \pm 0.3$ | $5.8 \pm 0.2$ | $1.68 \pm 0.09$ |
| S20G hIAPP (apo) | $1.25 \pm 0.01$ | $11.4 \pm 0.6$ | $8.7 \pm 0.6$ | $1.43 \pm 0.09$ |
| WT hIAPP + YX-I-1 | $1.25 \pm 0.01$ | $16.1 \pm 0.8$ | $5.4 \pm 0.8$ | $2.81 \pm 0.17$ |
| S20G hIAPP + YX-I-1 | $1.26 \pm 0.01$ | $12.3 \pm 0.5$ | $7.5 \pm 0.5$ | $1.52 \pm 0.08$ |
| WT hIAPP + YX-A-1 | $1.26 \pm 0.01$ | $12.8 \pm 0.9$ | $6.7 \pm 0.9$ | $1.67 \pm 0.14$ |
| S20G hIAPP + YX-A-1 | $1.27 \pm 0.01$ | $12.5 \pm 1.1$ | $4.9 \pm 1.1$ | $1.81 \pm 0.28$ |

We assessed the accuracy of the WT hIAPP conformational ensemble obtained from MD by computing backbone NMR chemical shifts of the 300 K REST2 replica with the chemical shift prediction software SPARTA+^57^ and comparing them with previously reported NMR chemical shift assignments (BMRB 34069)^25^ (Table 2, Figure S4). We also compared simulated helical propensities to those predicted from experimental backbone NMR chemical shifts with the CheSPI program^58^ (Figure S4). We observed good overall agreement with experimental chemical shifts, with root-mean-square deviations (RMSDs) between predicted and experimental shifts that are comparable to values obtained from the best performing force fields in benchmarks of MD simulations of IDPs^55^. Ca shifts calculated from the 300K REST2 ensemble were largely within the estimated 0.92 ppm SPARTA+ Ca shift prediction error for all residues, apart from larger deviations in residues ^3^NTA^5^, which are located in the disulfide bond constrained ^2^CNTATC^7^ region where we observed a ∼10-15% underestimation of the helical propensities relative to CheSPI predictions (Figure S4).

**Table 2.** Agreement between calculated and experimental backbone NMR chemical shifts from WT hIAPP MD ensembles. . The root-mean-square-deviation (RMSD) between predicted and experimental backbone NMR chemical shifts from unbiased MD and maximum-entropy reweighted^59,60^ WT hIAPP ensembles. Chemical shifts were calculated from the 300 K replica of an unbiased REST2 MD simulation performed with the a99SB-*disp* force field with SPARTA+^57^. Only Cα shifts were used as restraints for reweighting. Experimental chemical shifts were obtained from BMRB entry 34069.

| | Cα | HN | N | Cβ |
| --- | --- | --- | --- | --- |
|  | Unbiased WT hIAPP MD Ensemble |  |  |  |
| RMSD | **0.76** | 0.21 | 1.77 | 0.57 |
| | Cα-Reweighted WT hIAPP Ensemble | | | |
| RMSD | **0.47** | 0.14 | 1.33 | 0.35 |

To further quantify deviations between the unbiased REST2 MD ensemble and experimental NMR chemical shifts we performed maximum entropy reweighting^59,60^ using backbone Cα chemical shifts as restraints. We have deposited the reweighted ensemble of WT hIAPP in the Protein Ensemble Database^61^ (accession code pending). We observed a reduction in the RMSD of restrained Cα chemical shifts from 0.76 ppm to 0.47 ppm upon reweighting and observed similar improvements in unrestrained HN, N and Cβ chemical shifts, demonstrating that the reweighted ensemble is not overfit to the Cα chemical shift data (Table 2). The reweighted hIAPP ensemble is in closer agreement with experimental estimates of helical propensities (Figure S4). The reweighting results demonstrate that with the exception of a ∼10-15% underestimation of helical propensity within the disulfide constrained ^2^CNTATC^7^ region and a ∼10% overestimation of helical propensities in residues ^18^HSSNNFG^24^ and residues ^28^LLSSTNVGS^34^, the unbiased REST2 WT hIAPP MD ensemble is in good agreement with NMR chemical shift data and predicted secondary structure propensities. This demonstrates that the a99SB-*disp* force field and water model provide an accurate description of the conformational ensemble of hIAPP. Based on the similarity of the unbiased and reweighted ensembles of WT hIAPP (Table 1) and given that no backbone NMR chemical shift assignments have been reported for apo S20G hIAPP nor WT or S20G hIAPP in the presence of ligands, in the remainder of this manuscript we present direct comparisons of unbiased MD ensembles from the 300 K replica of each REST2 simulation performed. This enables direct comparisons between conformational ensembles generated using the same force field and modeling approach for all apo and ligand-binding simulations.

We determined the effects of the S20G mutation on the conformational ensemble of hIAPP by comparing the secondary structure propensities, populations of intramolecular contacts, radii of gyration (R_g_) and values of the α-helical order parameter Sα of the WT and S20G hIAPP ensembles (Figure 1, Table 1). We observed that the S20G mutation introduces a flexible central hinge that increases the populations of intramolecular interactions and β-sheets formed by residues ^7^CATQRLANFLV^17^ and ^23^NNFGAIL^27^ (which are located within the hIAPP amyloid core^32,33,39^) relative to WT hIAPP. We observed an average per-residue β-sheet population of 8.7 ± 0.6 % in the S20G hIAPP ensemble compared to 4.9 ± 0.4 % in the unbiased WT hIAPP ensemble. Correspondingly, the S20G mutation substantially reduces the helical propensity in residues ^19^SGNNFG^24^ relative to WT. The S20G ensemble is moderately more compact (R_g_=1.25 ± 0.01 nm) relative to WT (R_g_=1.30 ± 0.01 nm), due to increased populations of β-hairpins and intramolecular contacts.

**Figure 1.**
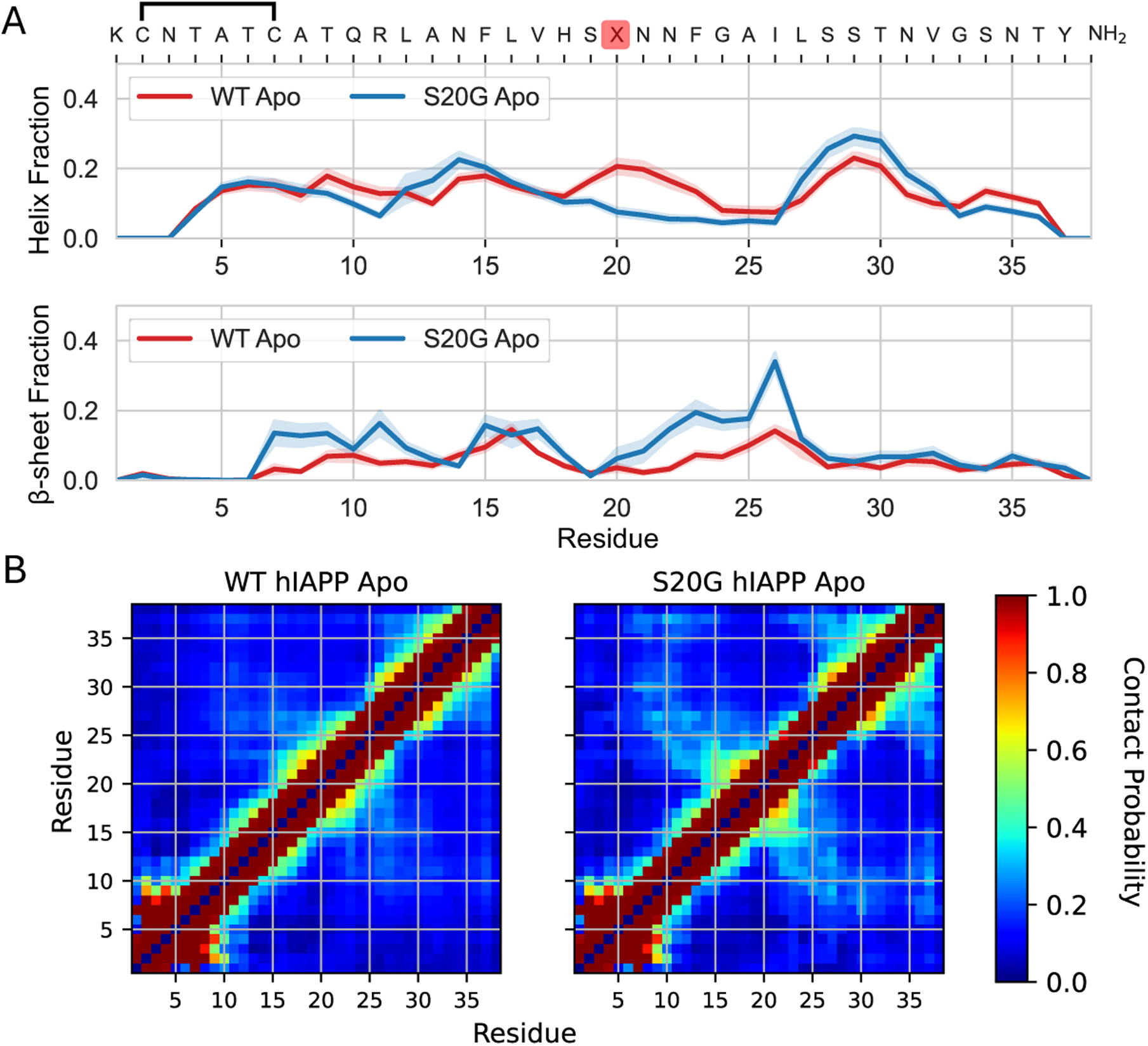
Comparison of conformational ensembles of apo wild-type (WT) and S20G hIAPP from enhanced sampling MD simulations. (A) Populations of α-helices and β-sheets observed in the 300 K replica of explicit solvent REST2 MD simulations of WT (red) and S20G (blue) hIAPP in their apo state. Secondary structure propensities were calculated with the DSSP algorithm. Shaded regions indicate statistical error estimates from blocking. The simulated hIAPP sequence is shown, with residue X corresponding to Ser in WT and Gly in S20G hIAPP. (B) Populations of intramolecular contacts in the apo WT and S20G hIAPP ensembles. Contacts between residues were defined using an 8 Å cutoff between the closest pair of heavy (non-hydrogen) atoms.

We applied a recently developed circuit topology approach^51^ to identify common conformational substates in the WT and S20G hIAPP ensembles (See “Circuit Topology Analysis” in Methods, Figure S5). Briefly, this framework assigns all intramolecular contact pairs occurring in a protein conformation as occurring in series, concerted series, parallel, parallel^-1^, concerted parallel, concerted parallel^-1^ or crossing based on their relative positions in the protein sequence to identify topological patterns in conformational space that may not be discernable from information on intramolecular contacts alone. We assigned elementary circuit topology classifications to all contact pairs in each frame of the 300 K replicas of all systems simulated in this work (including ligand-binding simulations, described below) and performed principal component analysis (PCA) on the resulting circuit topology assignment matrices to obtain a single two-dimensional (2D) latent space that represents the topological similarity of all hIAPP conformations sampled in apo and ligand binding simulations (Figure S6). We performed k-means clustering of the data points on these 2D projections (Figure S7, details in “Determination of k-means Clusters” in Supplementary Information) and identified six clusters as a good compromise between producing a tractable number of distinct conformational states for further analyses and various clustering validation metrics (Figure 2, Figures S7-S9). We observed that cluster 6 is only substantially populated in the simulation of S20G hIAPP in the presence of YX-A-1 (Table S2).

**Figure 2.**
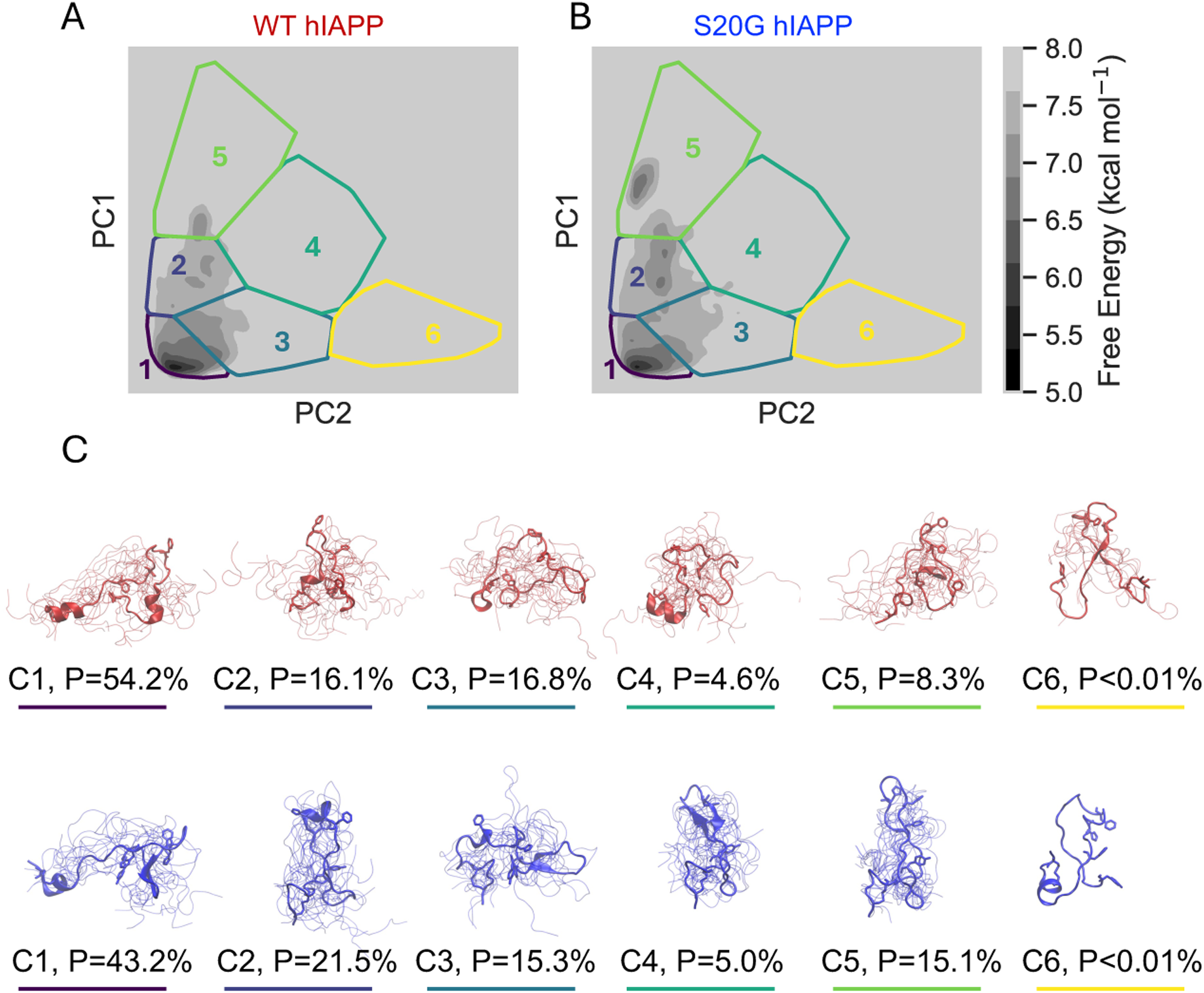
Conformational substates of WT and S20G hIAPP identified by clustering in a circuit topology latent space. (A, B) Free energy surfaces of apo WT (A) and S20G (B) hIAPP projected onto a latent space derived from circuit topology assignments. Principal component analysis (PCA) was performed on matrices of circuit topology assignments to produce a two-dimensional latent space that reflects the topological similarity of sampled conformations. k-means clustering was applied to all data points in this latent space to identify six conformational substates (cluster boundaries shown). (C) Representative structural ensembles of each cluster for WT (red) and S20G (blue) hIAPP. The structure closest to the average coordinates of each cluster is shown as a cartoon, with additional conformations are depicted as backbone traces. Cluster populations

Projections of the WT and S20G hIAPP apo ensembles on the circuit topology latent space and representative conformational ensembles of each k-means cluster are shown in Figure 2. We compared the populations of each cluster in each ensemble (Figure 2, Table S2) and the populations of intramolecular contacts and secondary structure elements within each cluster in the WT and S20G hIAPP ensembles (Figure S8, Figure S9). The WT and S20G hIAPP apo ensembles display similar patterns of intramolecular contacts in corresponding clusters, with higher contact populations in the S20G ensemble. We observed relatively small deviations in the secondary structure populations of each cluster in the apo WT hIAPP ensemble (Figure S8), and larger differences in the secondary structure populations among clusters in the apo S20G ensemble (Figure S9). We observed that cluster 1, which has the largest population (P) in the WT (P=54.2%) and S20G (P=43.2%) ensembles, is the most disordered and has the lowest populations of intramolecular contacts and secondary structure elements. This cluster is more populated in the WT hIAPP ensemble, while the S20G ensemble has higher populations of clusters 2 and 5. These S20G clusters have higher populations of intramolecular contacts and β-sheets involving residues ^23^NNFGAIL^27^, which are found in the amyloid core of hIAPP fibrils.

### Comparing binding modes of YX-I-1 and YX-A-1 to WT and S20G hIAPP

We utilized the REST2 MD simulation protocol described above to perform simulations of WT and S20G hIAPP in the presence of the small molecule ligands YX-I-1 and YX-A-1 (chemical structures shown in Figure 3). Each ligand was parameterized using the GAFF1 forcefield^62^. All four hIAPP:ligand systems were simulated for at least 2.7 µs per replica for a minimum aggregate simulation time of 54 µs (Table S1). To maintain consistency with the analysis of apo simulations, we discarded the first 1µs of ligand binding simulations as equilibration (Figure S3). We evaluated convergence in ligand binding REST2 simulations by comparing the secondary structure populations observed in each temperature replica and each independent demultiplexed (Figure S10-S13), assessing the time evolution and statistical error estimates of the ligand bound fraction (Figure S14) and additional simulated conformational properties (Table 1, Figure S15). Circuit topology classifications of the 300 K replica of all ligand-binding simulations were included in the previously described PCA and clustering analysis, and each frame of the ligand-binding simulations was assigned to one of the six previously described clusters (Table S2, Figure S16-S19).

**Figure 3.**
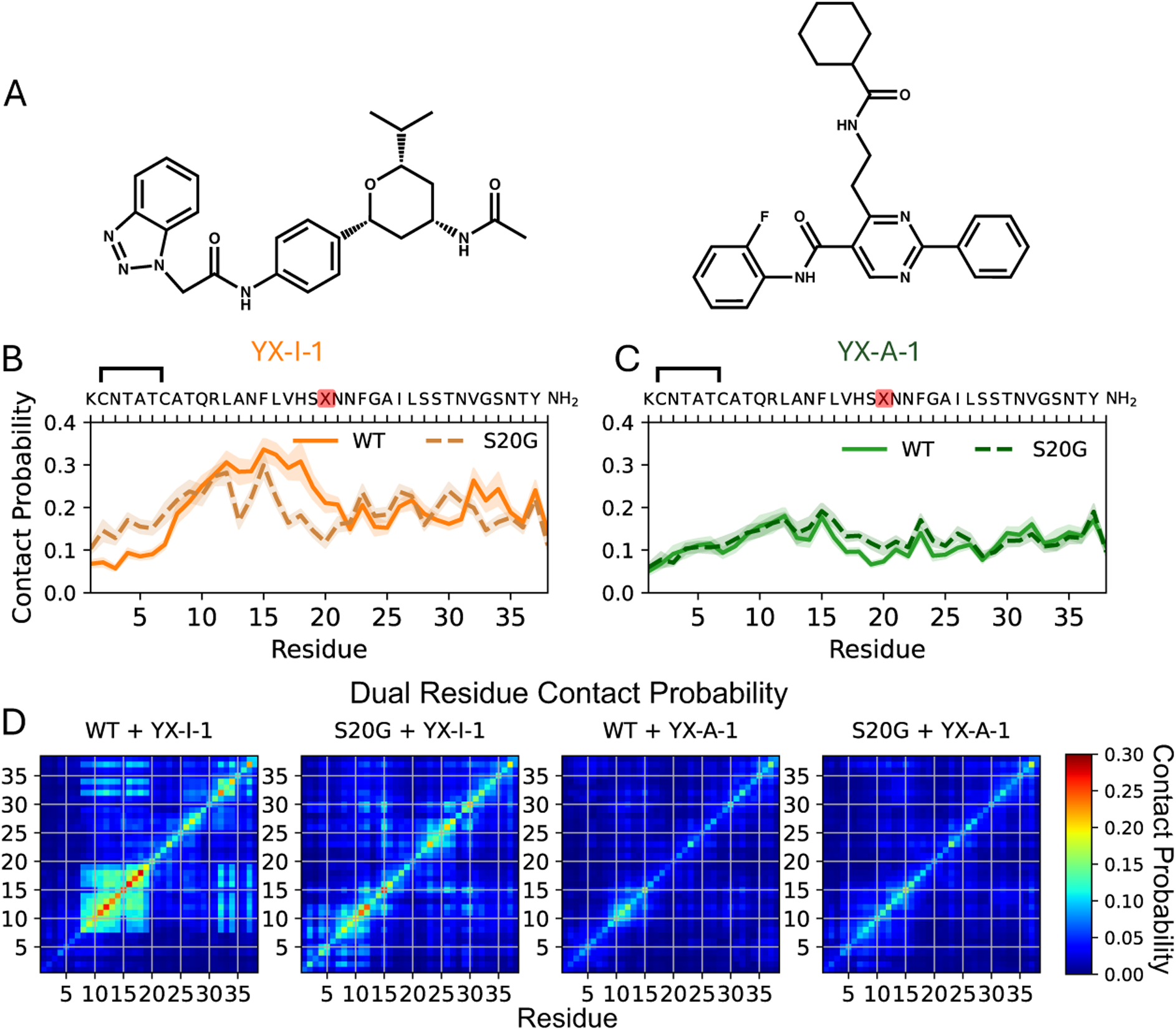
Binding modes of YX-I-1 and YX-A-1 to WT and S20G hIAPP. (A) Chemical structures of YX-I-1 and YX-A-1. (B,C) Populations of intermolecular contacts between ligands and each residue of hIAPP in the 300 K replica of REST2 MD simulations. Solid lines indicate WT hIAPP and dashed lines indicate S20G hIAPP. Shaded regions indicate statistical error estimates from blocking. The simulated hIAPP sequence is shown, where residue X corresponds to Ser in WT and Gly in S20G hIAPP. Protein-ligand contacts were defined using a 6 Å cutoff between heavy atoms. (D) The probability that each pair of hIAPP residues simultaneously forms a contact with a ligand in WT and S20G binding simulations.

We assigned all frames where at least one heavy (non-hydrogen) ligand atom is within 6 Å of a protein heavy atom as “bound frames” (See “Analysis of Protein-Ligand Intermolecular Interactions” in Methods). We assessed the convergence of the bound fraction of the 300 K temperature replica of each ligand-binding simulation, observing converged estimates of bound-fractions in the final 1μs of these simulations (Figure S14). The secondary structure populations and the populations of intramolecular contacts observed in the 300 K temperature replicas of ligand binding simulations of WT and S20G hIAPP are displayed in Figure S15. The populations of secondary structure elements and protein ligand contacts observed in each cluster of ligand-binding simulations are displayed in Figures S16-S19.

The simulated bound fractions and corresponding simulated K_D_ values of each ligand to WT and S20G hIAPP are shown in Table 3. These values are compared to the change in the helical fraction of ligand-binding simulations relative to apo simulations, previously reported *in vitro* effects on hIAPP fibril formation kinetics, the average values of previously reported ^1^H-^15^N NMR chemical shift perturbations (CSPs) observed in ligand titrations, and relative monomer binding affinities assessed by a combination native electrospray ionization mass spectrometry (nESI-MS), fluorescence emission quenching and surface plasmon resonance (SPR)^20^. NMR CSPs were measured in 20μM samples of hIAPP in the presence of 100μM YX-I-1 or 20μM YX-A-1, due to the limited solubility of YX-A-1^20^. The relative magnitudes of CSPs from YX-I-1 and YX-A-1 may therefore not be directly quantitively comparable. The weaker monomer binding of YX-A-1 to WT and S20G hIAPP suggested by smaller NMR CSPs is, however, supported by additional biophysical measurements^20^. Collectively, these measurements indicate that YX-I-1 directly binds monomeric WT hIAPP, while binding of YX-I-1 to monomeric S20G and of YX-A-1 to monomeric WT or S20G hIAPP is substantially weaker and below the detection limits of several assays. Kinetic analyses of ThT fluorescence assays suggest that YX-A-1 predominantly engages with sparsely populated oligomeric species formed via primary nucleation, accelerating the formation of higher-order species during the lag phase of aggregation of both WT and S20G hIAPP^20^.

**Table 3.**
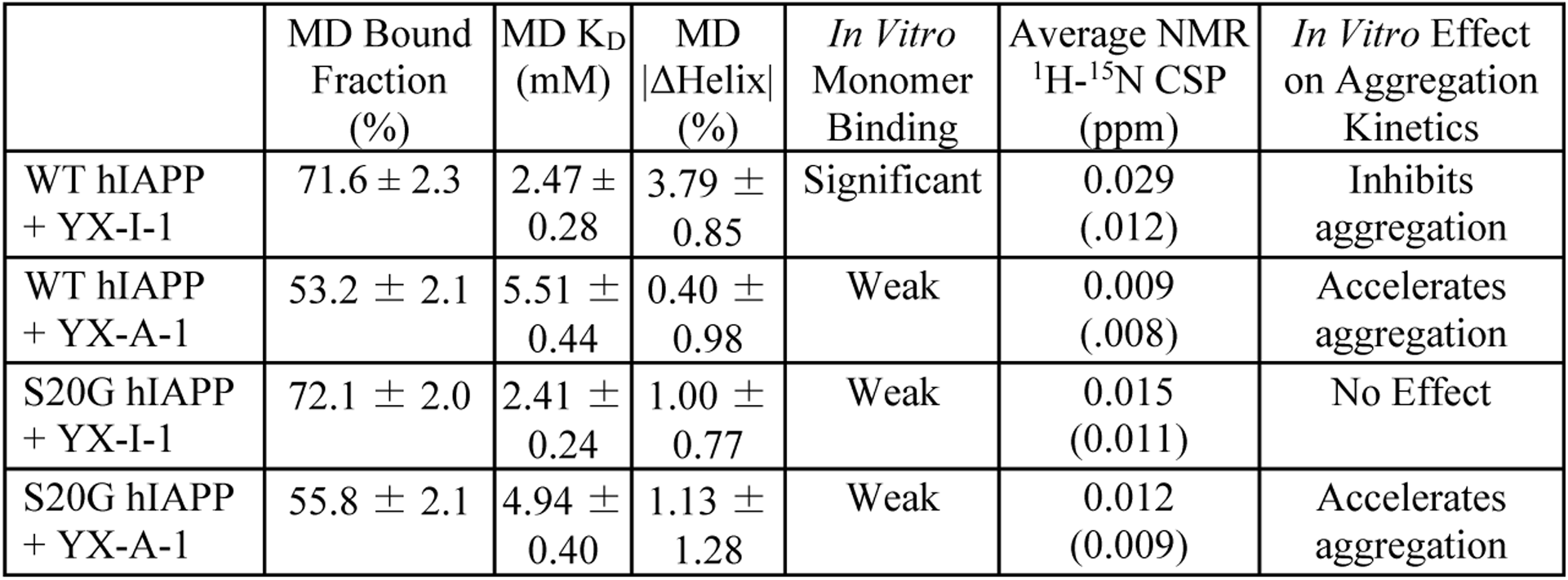
Bound Fractions of ligand binding REST2 MD simulations compared to previously reported experimental measurements. The strength of *in vitro* monomer binding was previously assessed by NMR, nESI-MS, fluorescence quenching and SPR measurements^20^. The average value of previously reported ^1^H-^15^N NMR chemical shift perturbations (CSPs) are shown along with their standard deviations (in parentheses). Experimental NMR ^1^H-^15^N CSPs of each residue were calculated according to 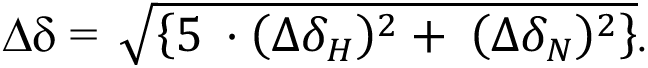 NMR CSPs were measured in 20μM samples of hIAPP in the presence of 100μM YX-I-1 or 20μM YX-A-1 (due to the limited solubility of YX-A-1). *In vitro* effects on the kinetics of hIAPP fibril formation were determined by kinetic analysis of thioflavin T (ThT) fluorescence assays. |ΔHelix| is the magnitude of the change in the helical fraction of ligand-binding simulations relative to apo simulations.

|  | MD Bound Fraction (%) | MD K<sub>D</sub> (mM) | MD \|ΔHelix\| (%) | *In Vitro* Monomer Binding | Average NMR <sup>1</sup>H-<sup>15</sup>N CSP (ppm) | *In Vitro* Effect on Aggregation Kinetics |
| --- | --- | --- | --- | --- | --- | --- |
| WT hIAPP + YX-I-1 | 71.6 ± 2.3 | 2.47 ± 0.28 | 3.79 ± 0.85 | Significant | 0.029 (.012) | Inhibits aggregation |
| WT hIAPP + YX-A-1 | 53.2 ± 2.1 | 5.51 ± 0.44 | 0.40 ± 0.98 | Weak | 0.009 (.008) | Accelerates aggregation |
| S20G hIAPP + YX-I-1 | 72.1 ± 2.0 | 2.41 ± 0.24 | 1.00 ± 0.77 | Weak | 0.015 (0.011) | No Effect |
| S20G hIAPP + YX-A-1 | 55.8 ± 2.1 | 4.94 ± 0.40 | 1.13 ± 1.28 | Weak | 0.012 (0.009) | Accelerates aggregation |

In ligand binding REST2 simulations of WT hIAPP, YX-I-1 and YX-A-1 exhibited bound fractions of 71.6 ± 2.3 % and 53.2 ± 2.1 %, respectively. Given the concentrations of ligand and protein in the simulation box (6.25 mM each), these bound fractions correspond to simulated K_D_ values of 2.35 ± 0.21 mM and 5.00 ± 0.32 mM, respectively, which are consistent with the small magnitudes of NMR CSPs observed in ligand titrations^20^. We observed an increase in helical fraction of the WT hIAPP ensemble from 12.4 ± 0.4 % in its apo state to 16.1 ± 0.8 % in the YX-I-1 binding simulation. The helical fraction of WT hIAPP does not appreciably change in the YX-A-1 binding simulation (12.8 ± 0.9 %). The stronger simulated binding affinity of YX-I-1 and larger hIAPP conformational changes observed upon YX-I-1 binding are consistent with larger ^1^H-^15^N NMR CSPs and additional biophysical experiments suggesting stronger binding of YX-I-1 to monomeric WT hIAPP compared to YX-A-1^20^.

In ligand binding simulations of S20G hIAPP, YX-I-1 and YX-A-1 exhibited bound fractions of 72.1 ± 2.0 % and 55.8 ± 2.1 %, respectively. The simulated helical propensities of S20G hIAPP in its apo state and in the presence of each ligand are within the statistical error estimates of each simulation (Table 1). NMR measurements in the presence of YX-I-1 revealed substantially smaller NMR chemical shift perturbations (CSPs) in S20G hIAPP compared to WT hIAPP (Table 3, Figure S20-S22). nESI-MS and fluorescence quenching experiments indicated markedly weaker binding of YX-I-1 to S20G, and no detectable binding was observed by surface plasmon resonance (SPR). We observed that the bound fraction of YX-I-1 to S20G hIAPP was within statistical error estimates to the bound fraction of YX-I-1 to WT hIAPP in REST2 simulations. This suggests that the simulated affinity of this interaction may be overestimated in the force fields employed here. The secondary structure populations of S20G hIAPP exhibited smaller changes than WT hIAPP in the presence of YX-I-1 (Table 1, Figure S15) which is consistent with smaller experimental NMR CSPs (Figures S20-S22). This does not, however, resolve discrepancies with weaker binding observed by nESI-MS, fluorescence quenching, and SPR. It is possible that YX-I-1 weakly interacts with monomeric S20G hIAPP in solution, but with more diffuse binding modes relative to WT hIAPP that do not produce detectable signals in these assays.

We compared residue-level populations of intermolecular protein-ligand contacts in hIAPP in each ligand binding simulation (Figure 3, Figures S16-S19). We observed that protein ligand contacts are dispersed along the protein sequence, consistent with a heterogeneous ensemble of interconverting binding modes suggested by the small magnitudes of experimental NMR CSPs. The lowest populations of ligand contacts where consistently observed in the disulfide bond constrained ^2^CNTATC^7^ region (Figure 3), consistent with a lack of CSPs in residues ^2^CNT^5^ in experiments (Figure S20-S21). To identify multisite binding interactions, we measured the probability that each pair of hIAPP residues simultaneously form contacts with a ligand in the same frame in each binding simulation (Figure 3D, Figures S16-S19). We observed that the ligand-bound ensemble of YX-I-1 to WT hIAPP has a higher population of binding modes that simultaneously engage residues ^8^ATQRLANFLVHS^19^ and residues ^32^VGSNTY^37^.

We compared the previously reported NMR CSPs of each residue to the per-residue populations of ligand contacts and changes in helical propensity relative to apo simulations in ligand-binding simulations in Figure S20 (YX-I-1) and Figure S21 (YX-A-1). Comparisons between experimental NMR CSPs and ligand-bound ensembles are, however, not straightforward. Experimental NMR CSPs from IDP-ligand binding will have contributions from both direct interactions with ligands and conformational changes in IDP ensembles induced by ligand binding. These effects are challenging to disentangle, as the directions of CSPs from different contributions can either reinforce or oppose one another. Additionally, through-space interactions from ligand binding can have competing effects on protein chemical shifts; for example, upfield CSPs in a nucleus produced by ring currents in a subset of binding poses can be effectively offset by downfield CSPs from ring currents in another subset of binding poses^63^. Simulated intermolecular contact probabilities and local protein conformational changes may therefore not always serve as direct proxies for the magnitude of experimental NMR CSPs. Acknowledging these caveats, we compared the average magnitude of ^1^H-^15^N NMR CSPs observed in ligand titrations with the simulated bound fractions, simulated changes in helical fraction, and average per-residue contact probabilities of each ligand in Figure S22. We observed that experimental NMR CSPs are most highly correlated with simulated changes in helical fractions in each protein (r^2^=0.97).

NMR measurements of 20μM WT hIAPP in the presence of 100μM YX-I-1 showed small but significant ^1^H-^15^N CSPs centered at residues ^10^QRLA^13^, ^17^VHSS^20^, ^23^FGAILS^28^ and the C-terminal residue Y37 (Figure S20). We did not observe a quantitative correlation between residue-level populations of ligand contacts or fluctuations in helical propensity and the magnitude of NMR CSPs but observed some qualitative similarities. In WT hIAPP, residue pairs in the ^8^ATQRLANFLHS^19^ region frequently formed simultaneous binding interactions with YX-I-1 (Figure 3D), which is qualitatively consistent with the large NMR CSPs observed in residues ^10^QRLAN^14^ and ^17^VHSS^20^. Additional simultaneous interactions with YX-I-1 were formed by residues 8-19 and residues ^32^VGSNTY^37^. In WT hIAPP, YX-I-1 binding produced the largest increase in helix fraction within residues A5 to S19, consistent with elevated experimental CSPs in this region, although CSPs were notably absent at residues ^14^NFL^16^. In WT hIAPP, residues ^23^FGAILS^28^ showed experimental NMR CSPs that do not correspond to either high ligand contacts populations or increased helical propensities in YX-I-1 binding simulations.

NMR measurements of 20μM S20G hIAPP in the presence of 100μM YX-I-1 produced substantially smaller ^1^H–^15^N CSPs compared to WT, with relatively small perturbations (<0.04 ppm) observed at residues L16, S19, N21, G24, I26, and V32. In the S20G hIAPP YX-I-1 binding simulation, we did not observe clear correlations between residue-level ligand contact probabilities or changes in helical populations with experimental ^1^H–^15^N CSPs. NMR measurements of 20μM WT hIAPP in the presence of 20μM YX-A-1 produced almost no detectable CSPs in WT hIAPP, with the exception of a small CSP (∼0.04ppm) at S19 (SI Figure S21). We observed that S19 had the largest increase in helix population relative to apo hIAPP in the YX-A-1 binding simulation. 20uM YX-A-1 produced extremely minor CSPs in S20G hIAPP, with largest CSPs observed at residues L16 and S19 (∼0.03ppm and ∼0.04ppm, respectively). These CSPs do not correlate with notable residue-level ligand contact probabilities or changes in helical populations observed in YX-A-1 binding simulations.

### YX-I-1 and YX-A-1 binding modes leave specific ligand moieties exposed to the solvent

In each ligand binding simulation, we computed the fraction of frames that each ligand heavy-atom is within 5 Å of a protein heavy-atom (Figure 4A). In YX-I-1 binding simulations, we observed that the benzotriazole group, the amide group connecting the benzotriazole and phenyl group (which we refer to as the amide linker), and phenyl groups of YX-I-1 had the highest protein contact probabilities with WT and S20G hIAPP (Figure 4, Figure S23). In the WT hIAPP:YX-I-1 binding simulation at least one heavy-atom of the benzotriazole group, the amide linker, and the phenyl group formed protein contacts in 65.2 ± 2.5%, 65.6 ± 2.5%, and 64.0 ± 2.5% of simulation frames, respectively. In contrast, the tetrahydropyran group, the isopropyl group, and the terminal amide group of YX-I-1 formed protein contacts in 60.5 ± 2.4%, 45.8 ± 1.7% and 48.5 ± 1.8% of simulation frames, respectively. In the S20G hIAPP:YX-I-1 binding simulation similar contact probabilities were observed for the benzotriazole group (65.1 ± 2.2%), the amide linker (65.2 ± 2.2%), and the phenyl group (63.3 ± 2.3%), while lower contact probabilities were observed for the tetrahydropyran group (57.0 ± 2.1%), the isopropyl group (43.2 ± 1.4%) and the terminal amide group (43.2 ± 2.0%). Based on these observations we hypothesize that the monomer affinity of YX-I-1 is predominantly conferred by a binding motif consisting of the benzotriazole group, the amide linker, and the phenyl group and that these groups are largely buried and sequestered from solvent upon binding. We hypothesize that the more solvent exposed tetrahydropyran, isopropyl group and terminal amide groups may be more important for inhibiting interactions between hIAPP monomers and higher order hIAPP species in amyloid misfolding pathways.

**Figure 4.**
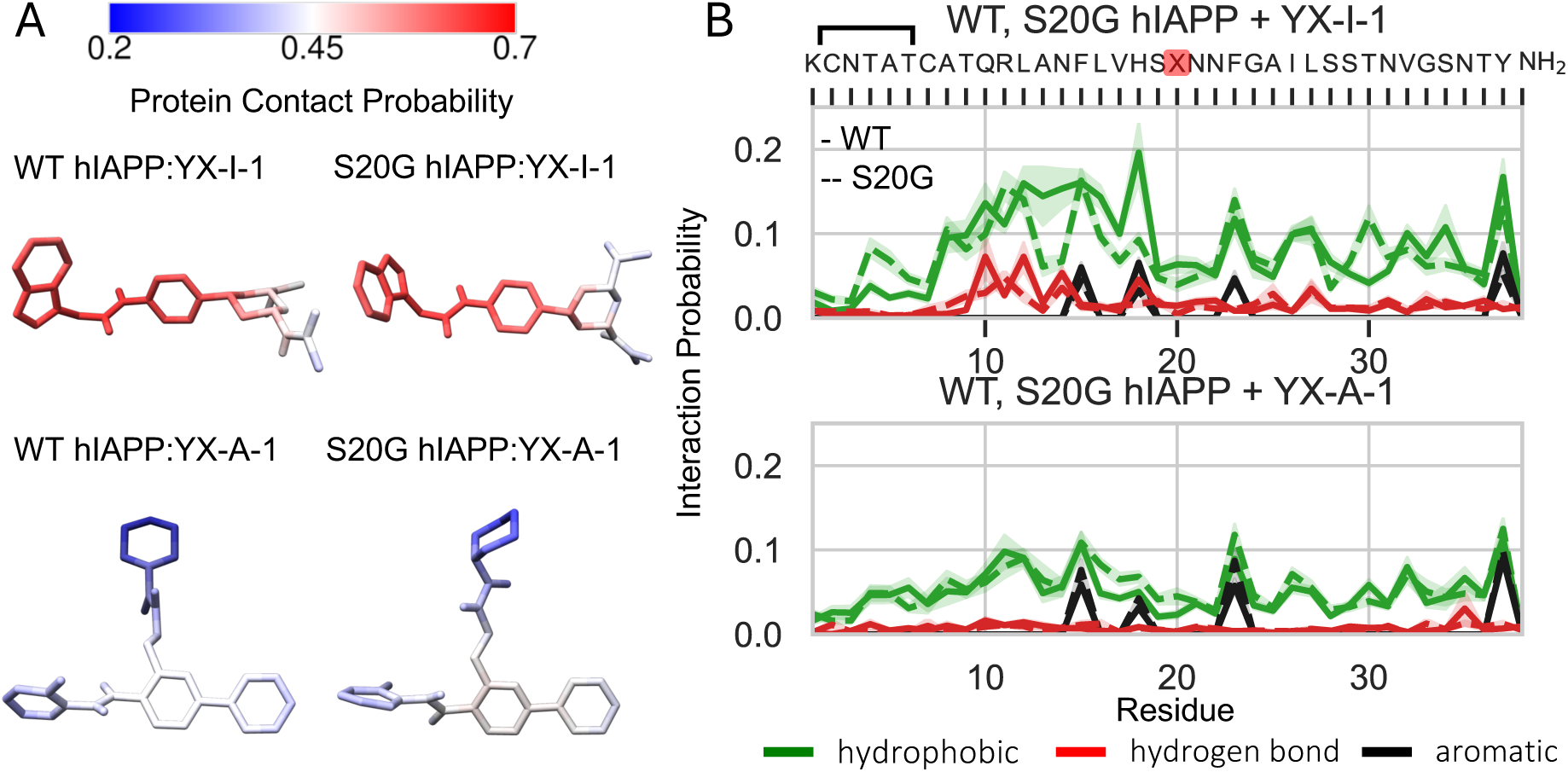
Protein-ligand interactions in YX-I-1 and YX-A-1 hIAPP binding simulations. (A) The fraction of simulation frames where each ligand atom is within 5 Å of a protein atom. (B) Populations of intermolecular interactions observed in hIAPP binding simulations. Populations of hydrophobic (green), aromatic stacking (black), and hydrogen bonding (red) interactions are shown for WT (solid lines) and S20G (dashed lines) hIAPP.

In the WT hIAPP:YX-A-1 binding simulation we observed that all chemical groups (phenyl, pyrimidine, amide, fluorobenzyl, alkyl amide, and cyclohexane) – had protein contact probabilities less than 46.0% (Figure S23). The lowest protein contact probabilities were observed for the cyclohexane group (34.3 ± 1.3%) and the fluorobenzyl group (41.6 ± 1.5%) (Figure S23). We observed similar protein contact probabilities (within statistical error estimates) for all chemical groups of YX-A-1 in S20G hIAPP binding simulations. We note that the most solvent exposed cyclohexane and fluorobenzyl moieties of YX-A-1 are more hydrophobic than the solvent exposed amide and isopropyl groups of YX-I-1. We hypothesize that the presence of substantial solvent-exposed hydrophobic surface area of YX-A-1 when it is weakly bound to hIAPP monomers or oligomers could accelerate association of hIAPP into higher order species in amyloid misfolding pathways.

### Comparing populations of intermolecular interactions between YX-I-1 and YX-A-1 and hIAPP

To understand the molecular interactions that confer affinity of each ligand to hIAPP we calculated the per-residue populations of intermolecular hydrophobic interactions, hydrogen bonds, and aromatic stacking interactions (See “Analysis of Protein-Ligand Intermolecular Interactions” in methods) formed between YX-I-1 and YX-A-1 and each residue of WT or S20G hIAPP in each ligand binding simulation (Figure 4B). YX-I-1 generally exhibited elevated populations of hydrophobic interactions and hydrogen bonds relative to YX-A-1 throughout the hIAPP sequence, with substantially elevated populations of hydrophobic contacts and hydrogen bonds in residues ^8^ATQRLANFLHS^19^.

We assessed the populations of all hydrogen bond donor-acceptor pairs in WT hIAPP:YX-I-1 binding simulations and observed a highly populated network of hydrogen bonds formed between the nitrogen and oxygen atoms of the benzotriazole group and amide linker of YX-I-1 and hIAPP residues Q10, L12, N14, H18 and L27. We observed that the most populated hydrogen bond pairs were formed between the backbone amide nitrogen of Q10 and the carbonyl oxygen of the amide linker, the backbone carbonyl oxygen of L12 and the amide linker nitrogen, the backbone amide nitrogen of L27 and the benzotriazole nitrogens, and the backbone amide nitrogen of N14 and the benzotriazole nitrogens. We also observed substantial populations of hydrogen bonds donated by the backbone amide nitrogen and protonated imidazole nitrogen of H18 and accepted by the benzotriazole nitrogens and the amide linker carbonyl oxygen. While the network of hydrogen bonds and hydrophobic contacts with residues ^8^ATQRLANFLHS^19^ is partially populated in each cluster of the WT hIAPP:YX-I-1 binding simulation, the populations of these interactions are particularly pronounced in cluster 4 (Figure S16), where they are coupled with hydrophobic and stacking interactions with residue Y37. Representative snapshots of this binding pose are shown in Cluster 4A in Figure 5.

**Figure 5.**
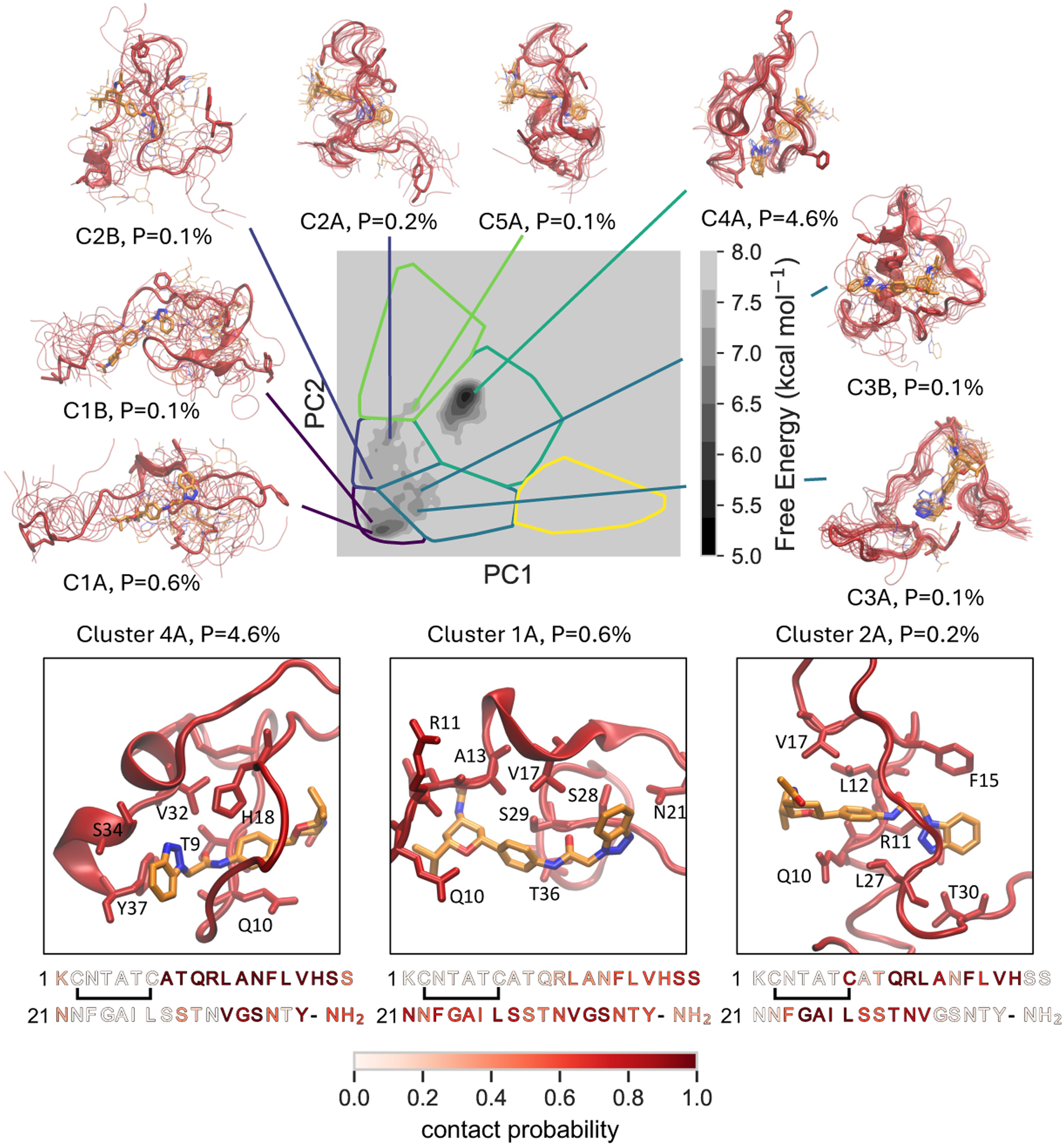
Multisite binding modes of YX-I-1 to WT hIAPP. Free energy surface of WT hIAPP conformations in which at least 15 residues form contacts with YX-I-1, projected onto the circuit topology latent space with cluster boundaries shown. Structures with ≥ 15 protein–ligand contacts represent 12.8% of all simulated conformations. Representative structural ensembles of the most populated free energy minima are shown, with populations (P) indicated. Panels display representative structures of the most populated multisite binding modes. Amino acid sequences below each panel are colored by per-residue protein–ligand contact probabilities of the corresponding substates.

We compared the overall populations of specific intermolecular interactions in binding simulations of YX-I-1 and YX-A-1 to WT hIAPP. The probability of observing at least one hydrogen bonding interaction was substantially larger for YX-I-1 (33.7 ± 1.7%) than YX-A-1 (12.1 ± 0.9%) in the WT hIAPP binding simulations. We observed that the probability of observing at least one protein-ligand stacking interaction in conformations of binding simulations was slightly larger for YX-I-1 (21.7 ± 1.4 %) than YX-A-1 (16.1 ± 1.0%). We observed more cooperative formation of stacking interactions and hydrogen bonding interactions in YX-I-1 binding simulations; the probability of simultaneously observing at least one aromatic-stacking interaction and one intermolecular hydrogen bonding interaction in the WT hIAPP binding simulation of YX-I-1 was 11.6 ± 1.1% compared to 4.2 ± 0.5% in the YX-A-1 binding simulation.

In previous investigations of the ligand binding to the IDPs α-synuclein^64^ and the androgen receptor activation domain^65–67^, it was observed that higher affinity ligands had substantially increased populations of aromatic stacking interactions in the ligand-bound ensembles, suggesting that increased propensities to form stacking interactions were a dominant factor for conferring stronger binding affinity. In contrast, we observed relatively similar per-residue populations of stacking interactions in the binding simulations of YX-I-1 and YX-A-1 to WT hIAPP (Figure S24). The average populations of stacking interactions of aromatic residues of WT hIAPP were 6.2 ± 0.9 % and 5.9 ± 0.7% for YX-I-1 and YX-A-1, respectively. This suggests that increased affinity of YX-I-1 to hIAPP is primarily driven by the network of cooperative hydrophobic contacts and hydrogen bonding interactions formed by residues ^8^ATQRLANFLHS^19^, rather than an increased propensity of aromatic groups of YX-I-1 to form stacking interactions.

### Visualizing Multisite Binding Modes of YX-I-1 and YX-A-1

Ligand binding simulations of YX-I-1 and YX-A-1 reveal heterogeneous ensembles of binding modes, rather than a small subset of stable protein-ligand complexes. These binding modes are consistent with the previously proposed dynamic shuttling mechanism of IDP ligands^64^, where specific intermolecular interactions stochastically break and reform as a ligand samples a large number of distinct binding poses distributed across the sequence of an IDP. Within each bound ensemble, however, we observe a smaller population of bound poses where ligands interact more globally with a larger number of hIAPP residues. To obtain more insight into these binding modes, we analyzed bound conformations where YX-I-1 and YX-A-1 form contacts with at least 15 residues of WT hIAPP. These conformations represent 12.8% and 3.9% of bound conformations observed in simulations YX-I-1 and YX-A-1 binding WT hIAPP, respectively. We projected these subensembles onto the circuit topology latent space of our simulations in Figure 5 and Figure 6. We identified minima from the projected free-energy surfaces and display representative structural ensembles of each minimum along with snapshots of representative binding poses of the most populated states.

**Figure 6.**
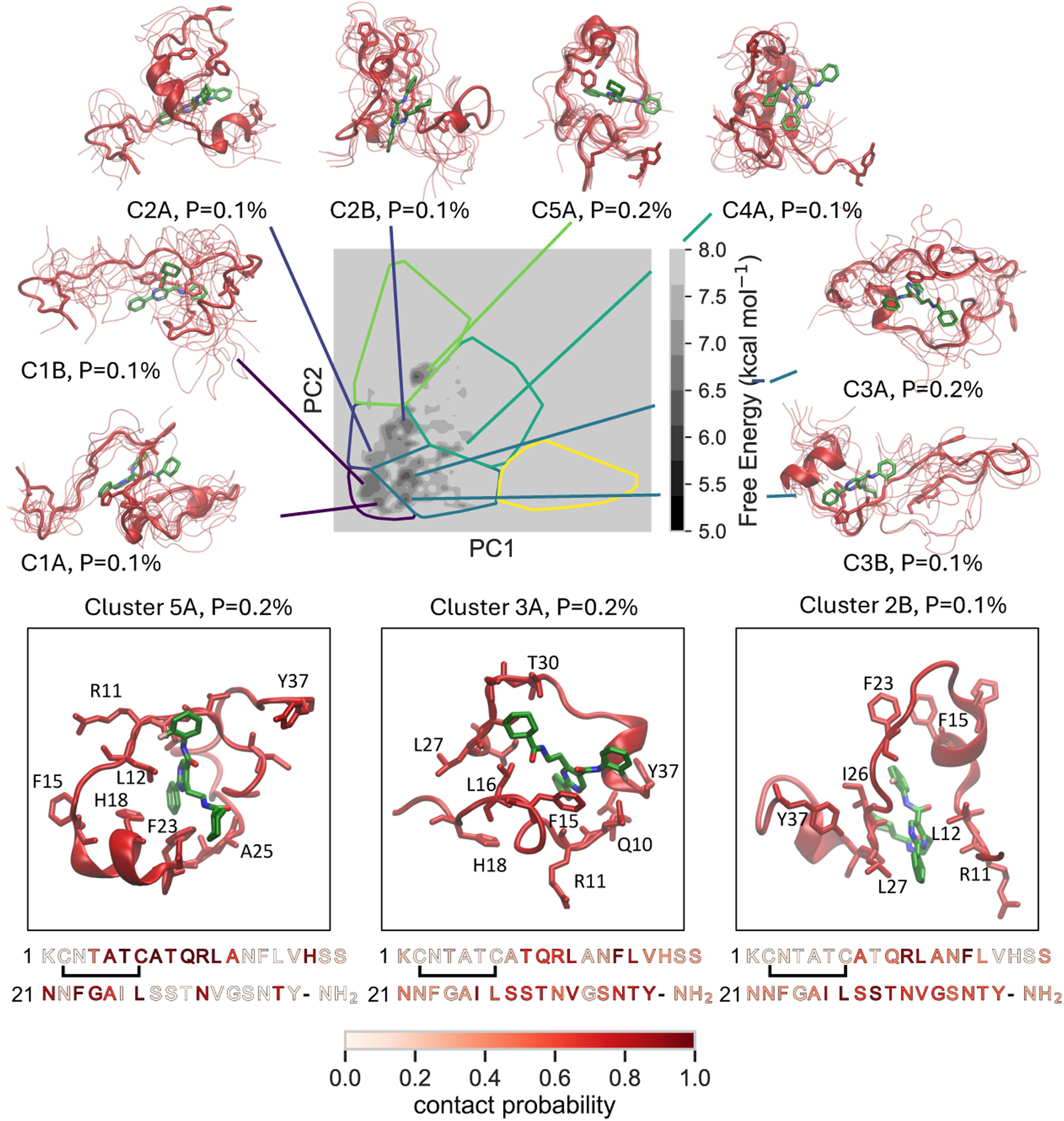
Multisite binding modes of YX-A-1 to WT hIAPP. Free energy surface of WT hIAPP conformations in which at least 15 residues form contacts with YX-A-1, projected onto the circuit topology latent space with cluster boundaries shown. Structures with ≥ 15 protein–ligand contacts represent 3.9% of all simulated conformations. Representative structural ensembles of free energy minima are shown, with populations (P) indicated. Panels display representative structures of the most populated multisite binding modes. Amino acid sequences below each panel are colored by per-residue protein–ligand contact probabilities of the corresponding substates.

We observed that several of the most populated multisite binding modes contain substantial conformational heterogeneity, with the backbone of hIAPP fluctuating about an average topology. The multisite binding modes of YX-I-1 contain several binding poses where the benzotriazole, amide linker and phenyl groups are held by two distinct regions of hIAPP while the tetrahydropyran, isopropyl and terminal amide groups protrude into solvent. It is notable that the most structurally homogenous multisite binding modes identified have very low populations, representing only 0.1-0.2% of simulation frames. A notable exception is state C4A (Figure 5), which has a population of 4.6% and contains high populations of hydrogen bonds formed between benzotriazole and amide linker groups of YX-I-1 with hIAPP residues Q10, L12, N14 and H18. Taken together, these results demonstrate that only a small subset of the bound ensembles of YX-I-1 and YX-A-1 contain well defined multisite binding poses, and that the ligand binding simulations do not sample appreciable populations of homogenous states where the entire ligand is enveloped in a stable binding pocket and fully sequestered from solvent.

## Discussion

We have employed all-atom, explicit solvent enhanced sampling MD simulations to provide an atomic-level characterization of the conformational ensembles of WT and S20G hIAPP in their apo state and in the presence of two ligands previously shown to modulate the kinetics of amyloid formation^20^. Our apo simulation of WT hIAPP was validated against backbone NMR chemical shifts, demonstrating that a99SB-*disp* force field and water model employed here provide an accurate description of this system. We observed that the S20G mutant introduces a flexible central hinge in hIAPP, increasing populations of intramolecular contacts and β-sheets formed by residues frequently found in hIAPP amyloid cores^32,34,35^. The S20G mutant has been shown to accelerate rates of both primary nucleation, amyloid surface-catalyzed secondary nucleation, and fibril elongation in hIAPP aggregation pathways^18–21^. Elevated populations of intramolecular contacts and β-sheets in S20G hIAPP could potentially increase rates of self-association into β-sheet rich oligomeric prefibrillar species and/or increase rates of conversions of disordered oligomeric assemblies into β-sheet rich fibrillar species.

Ligand binding simulations revealed that YX-I-1 has a substantially stronger binding affinity to disordered monomers of WT hIAPP than YX-A-1, which is consistent with previously reported biophysical experiments^20^. We observed that the stronger binding affinity of YX-I-1 to WT hIAPP is conferred by the formation of a network of hydrophobic interactions and hydrogen bonds with residues ^8^ATQRLANFLHS^19^. The binding modes of YX-I-1 have substantially elevated populations of protein-ligand hydrogen bonds (∼2.75-fold larger) compared to YX-A-1, and only a marginally larger population of aromatic stacking interactions (∼1.3-fold larger). This result contrasts with previous integrated experimental and computational investigations of molecular mechanisms of ligand binding to the disordered activation domain of the androgen receptor^65–67^, where the increased affinity of tighter binders was most correlated with higher populations of aromatic stacking interactions in higher affinity ligands. This suggests that rationally inserting networks of hydrogen bond donors and acceptors into ligand moieties that simultaneously interact with both polar and hydrophobic sidechains and polar backbone moieties may be a possible strategy for increasing the affinity of IDP ligands. We observed substantially higher populations of binding modes where YX-I-1 simultaneously forms interactions with multiple regions of WT hIAPP compared to YX-A-1. The observation that higher affinity IDP ligands form higher populations of binding modes that engage multiple regions of an IDP target is consistent with previous investigations into the binding mechanisms of androgen receptor activation domain and α-synuclein ligands^64–67^.

We observed specific moieties of YX-I-1 and YX-A-1 are consistently exposed to solvent in simulated binding modes. In binding simulations of YX-I-1, we observed the benzotriazole group, the amide linker, and the phenyl group are largely buried and sequestered from solvent upon binding, while the tetrahydropyran, isopropyl group and terminal amide groups consistently remain more exposed to solvent in bound conformations. Based on these observations we hypothesize that the monomer affinity of YX-I-1 may be predominantly conferred by a binding motif consisting of the benzotriazole group, the amide linker, and that the more solvent exposed tetrahydropyran, isopropyl and terminal amide groups may be more important for interactions between hIAPP monomers and higher order hIAPP species in amyloid misfolding pathways. In a recent study by Taylor, Radford and coworkers^36^ an analogue of YX-I-1 lacking the tetrahydropyran, isopropyl and terminal amide groups was found to have no effect on the kinetics of hIAPP amyloid formation, demonstrating the importance of these moieties for inhibiting aggregation.

In binding simulations of YX-A-1, we observed that the phenyl, amide, and alkyl amide groups have a higher propensity to be buried and sequestered from solvent, while the cyclohexane and fluorobenzyl groups are largely solvent exposed. We note that the solvent exposed cyclohexane and fluorobenzyl moieties of YX-A-1 are substantially more hydrophobic than the solvent exposed tetrahydropyran, amide and isopropyl groups of YX-I-1. We hypothesize that these solvent-exposed hydrophobic groups may be responsible for accelerating the formation of higher order oligomeric species of hIAPP when YX-A-1 is weakly bound to monomers and oligomers. It is, however, also possible that YX-A-1 may only engage and stabilize distinct conformations of oligomeric species on hIAPP amyloid formation pathways not sampled in monomer simulations. Simulations of the assembly of hIAPP into higher-order oligomeric species, and the effects of YX-I-1 and YX-A-1 on oligomer assembly will be the subject of future investigations.

We observed that YX-I-1 had a similar bound fraction to WT and S20G hIAPP in ligand-binding simulations. This result is inconsistent with previously experimental results demonstrating that YX-I-1 has a substantially lower affinity to monomeric S20G hIAPP than WT hIAPP, and that YX-I-1 has no effect on the aggregation rate of S20G hIAPP. This suggests that our simulations may overestimate the affinity of YX-I-1 to S20G hIAPP. We do, however, observe a larger population of YX-I-1 binding modes that simultaneously engage multiple regions of hIAPP in the WT hIAPP bound ensemble compared to the S20G bound ensemble. It is possible that these more cooperative binding modes are more effective for inhibiting oligomerization pathways and produce stronger signals in biophysical experiments. Failure of YX-I-1 to inhibit S20G aggregation could also potentially arise from a reduced affinity to higher order S20G oligomers relative to WT oligomers hIAPP.

In summary, we have modeled the conformational ensembles of monomers of WT and S20G hIAPP in their apo state and in the presence of ligands that modulate the rates of hIAPP amyloid formation using long-timescale, enhanced sampling REST2 MD simulations. We have provided a detailed dissection of the conformational ensembles of WT and S20G hIAPP in their apo and holo states and have elucidated ensembles of ligand binding modes that rationalize the higher affinity of the aggregation inhibitor YX-I-1 to monomeric hIAPP compared the aggregation accelerator YX-A-1. Our simulations suggest that distinct ligand moieties of YX-I-1 and YX-A-1 confer affinity to monomeric hIAPP, while solvent-exposed ligand moieties may modulate the kinetics of intermolecular association of ligand-bound hIAPP into higher-order oligomeric intermediates on amyloid aggregation pathways. This study provides an atomically detailed mechanistic framework for understanding the molecular driving forces of ligand binding to monomeric hIAPP that can inform the design of novel ligands to test in future structure-activity-relationship studies of potential hIAPP aggregation inhibitors. In future work, we plan to use molecular simulations to study how YX-I-1 and YX-A-1, and the recently discovered hIAPP aggregation inhibitors Canagliflozin and Doxazosin^36^, affect the formation of oligomeric species of hIAPP. This work, together with future investigations on the effects of ligands on the assembly of hIAPP into oligomeric species, will provide atomic resolution insights into the molecular interactions that drive the primary nucleation of hIAPP into oligomeric states on amyloid formation pathways. Characterizing these states and their interactions with small molecule ligands could inform the design of more potent T2D therapeutics.

## Methods

### Simulation Methods

We performed unbiased, all-atom, explicit solvent simulations of wild-type (WT) and S20G hIAPP with the replica exchange solute tempering (REST2) enhanced sampling algorithm^52–54^ in the presence and absence of the ligands YX-I-1 and YX-A-1. REST2 simulations were run with 20 replicas using a solute temperature ladder spanning 300-500 K using GROMACS v2022.3^68,69^ patched with PLUMED v2.8.0^70^. All protein and ligand atoms were selected as solute for solute tempering. hIAPP and water molecules were parameterized with the a99SB*-disp* protein force field and a99SB*-disp* water model^55^, respectively. Ligands were parametrized using the GAFF1 forcefield^62^ and Na^+^ and Cl^-^ ions were parameterized with Åqvist parameters^20^. Apo simulations of WT and S20G hIAPP monomers were run for 4.3 µs/replica and 3.8 µs/replica, with aggregate simulation times of 86 µs and 76 µs, respectively. Ligand-binding simulations of WT hIAPP with YX-I-1, S20G hIAPP with YX-I-1, WT hIAPP with YX-A-1, and S20G hIAPP with YX-A-1 were simulated for 3.4, 2.8, 2.7, 2.8 μs per replica with aggerate total simulation times of 68.0, 56.0, 54.0, and 56.0 μs, respectively. The combined simulation time in this investigation totals 396.0 μs.

Simulations were performed using amidated and oxidized human hIAPP constructs containing a disulfide bond between residues C2-C7 and a NH_2_ cap at the C-terminus. The first conformer of the previously determined solution NMR structure of amidated, C2-C7 oxidized WT human hIAPP (PDB ID: 2L86)^71^ was used as an initial conformation to generate simulation starting structures. This conformation was energy minimized and used a starting structure for a short 100 ps, 600 K high-temperature NVT unfolding simulation in vacuo. Twenty structures with varying levels of helical-content were selected from the unfolding trajectory. The PyMol command-line interface was used to generate an S20G mutant of each selected conformation. Each structure was solvated with ∼8600 water molecules in a cubic box of length 6.4 nm and neutralized with 4 Na^+^ ions and 7 Cl^-^ ions for a total concentration of 25 mM NaCl to match the salt concentrations of experimental data^20^. After solvation, each structure was energy minimized with a maximum force cutoff of 1000 kJ mol^-1^ nm^-1^. Following the energy minimization, the structures were subject to a 1 ns, 300 K NVT simulation performed with the velocity-rescale^72^ thermostat, a 1 fs timestep and bond lengths and angles constrained with the LINCS^73^ algorithm. Next, 1 ns NPT simulations with a 2 fs timestep and a target pressure of 1 bar were performed using Parinello-Rahman pressure coupling^74,75^ and a 300 K temperature maintained with the velocity rescaling thermostat^72^. Finally, structures were equilibrated for at least 20 ns at 300 K in the NPT ensemble with the velocity-rescale thermostat and Parrinello-Rahman pressure coupling to obtain 20 equilibrated starting structures for REST2 simulations.

Initial ligand conformations of YX-I-1 and YX-A-1 were generated using openbabel^76^ command-line interface to generate 3D conformers from smile strings. The ligands were parameterized with GAFF1^62^ force field parameters utilizing the ACPYPE^77^ server. Ligand conformations were added to each of 20 starting structures selected for used for apo simulations, and each system was solvated with ∼8600 water molecules in a cubic box of length 6.4 nm and neutralized with 4 Na^+^ ions and 7 Cl^-^ ions for a total concentration of 25 mM NaCl. The equilibration protocol described above was applied to each hIAPP and ligand system.

Production REST2 simulations were performed with the velocity-rescale thermostat, a 2 fs timestep and bond lengths and angles constrained with the LINCS^73^ algorithm. REST2 simulations were run with 20 replicas using a solute temperature ladder spanning 300-500 K and all protein and ligand atoms selected as solute. Coordinated exchanges between replicas were attempted every 1.6 ps. Simulation frames were saved every 80 ps. All GROMACS simulation input files are freely in accompanying GitHub repository.

The first microsecond of all simulations was discarded due to the slow equilibration of α-helical order parameter Sα^56^ (SI Figure S3). Sα measures the similarity of all 7-residue segments to an ideal helical structure (𝜙=-57,𝜓=-47) calculated according 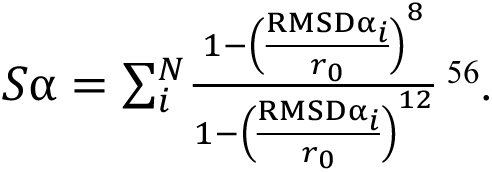

### Analysis of Protein-Ligand Intermolecular Interactions

Analyses were run utilizing MDTraj^78^, NumPy, Scikit-Learn^79^, and SciPy python packages. Bound frames were defined as any frame where at least one protein and ligand heavy (non-hydrogen) atom are within 6 Å. This cutoff was selected to reflect the distance that non-bonded interactions have been shown to have measurable effects on the NMR chemical shift predictions in proteins^57,63,80^. We note that changing this distance threshold does not change the ratio of bound fractions of each ligand. Intermolecular hydrophobic contacts were defined using a distance cutoff of 4 Å between protein and ligand carbon atoms. The Baker-Hubbard algorithm was used to identify hydrogen bonds with a distance cutoff of 3.5 Å between donated hydrogens and acceptor atoms and a donor-hydrogen-acceptor angle cutoff of >150°. Aromatic stacking interactions were calculated following Marsili et al. with modified distance and angle cutoffs^76^. All statistical error estimates were calculated using a blocking analysis^81,82^ with the pyblock python package. *K*_D_ values were calculated according to *K*_D_ = P_u_/P_b_(*vc*°N_A_)^−1^ where (P_b_) is the fraction of frames with at least one protein-ligand contact, (P_u_) is the fraction of frames with no ligand contacts, *v* is the volume of the simulation box, *N*_A_ is Avogadro’s number, *c*° is a standard state concentration (1 mol L^−1^), and (*vc*°N_A_)^−1^ is the simulated concentration. The simulated concentration of one copy of hIAPP (or ligand) in the 6.4 nm water boxes used here is 6.25 mM. Contact probabilities of each ligand heavy atom (Figure 4) were determined using a 6 Å distance cutoff to protein heavy atoms.

### Circuit Topology Analysis

For each conformation sampled in the 300 K solute temperature replicas of each simulated system, we identified intramolecular contacts using a 10 Å distance cutoff considering all atoms. After identifying contacts, all pairs of contacts in each frame were enumerated and assigned an elementary circuit topology relation following the matrix assignment scheme in Figure S5. Each contact pair (consisting of four contact sites) was assigned one of seven topological relations: series=1, concerted series=2, parallel=3, parallel^-1^=4, concerted parallel=5, concerted parallel^-1^ =6, and crossing=7 where numbers increase with increasing complexity as in Scalvini et al. 2023^51^ (Figure S6, Supplementary Information section “Analysis of Circuit Topology Assignments”). For each 300 K temperature replica, we obtain a circuit topology matrix containing assignments for all possible contact pairs in each frame. To obtain a single low-dimensional latent-space for all simulations performed in this work – we merged all circuit topology matrices from the 300 K temperature replica ensembles of each simulated system and performed principal component analyses (PCA) on the merged set of circuit topology assignments. For dimensionality reduction with PCA, we considered only the top half of the matrix of circuit topology assignments and chose to project the reduced latent space onto two principal components. We display the resulting projections of each 300 K temperature replica onto the shared circuit topology latent space in Figure S6. We then performed k-means clustering of circuit topology assignments in principal component space to identify conformational states across all simulated ensembles (see “Determination of k-means Clusters” in the Supplementary Information). We computed free energy surfaces of the 300 K solute temperature replica projected onto the first two PCs. Free energies were computed according to Δ𝐺 = −𝐾𝑇 ln(𝑃) where *P* is the population obtained using a 50 x 50 2D histogram grid defined by the two principal components (PCs) of the circuit topology latent space.

### Maximum-Entropy Ensemble Reweighting

We utilized the maximum entropy reweighting algorithm of Borthakur et. al. 2024^59,60^ with Cα NMR chemical shifts as restraints. The kish ratio of the resulting ensemble was 10.4, meaning the reweighted ensemble retains ∼10.4% of the frames from the unbiased MD simulation.

### Data and Code Availability

All MD trajectories, GROMACS simulation input files and code used for analyses are freely available from the accompanying GitHub repository (https://github.com/paulrobustelli/Garcia_hIAPP_monomer_binders_2025). The backbone NMR chemical shifts of amidated and oxidized WT hIAPP for trajectory reweighting were obtained from Biological Magnetic Resonance Bank entry 34069.

## Acknowledgements

P.R. acknowledges the support of NIH award R35GM142750 and grant CS-CSA-2024-080 from the Research Corporation for Science Advancement. M. G. acknowledges the support of Department of Energy award DE-SC0024386. K.R.M. acknowledges the support of the Dartmouth Magnuson Center for Entrepreneurship and grant CS-CSA-2024-080 from the Research Corporation for Science Advancement. The authors thank Prof. Sheena Radford and Dr. Alexander Taylor for valuable discussions and feedback. The authors thank Jiaqi Zhu for valuable discussions and technical assistance with MD simulations and Kaushik Borthakur for assistance with maximum entropy reweighting calculations.

## Author Contributions Statement

P.R. conceived, designed and supervised the research. M.G. performed research with technical assistance from K.M.R. M.G. performed MD simulations, designed and implemented simulation analyses and wrote code accompanying the manuscript. M.G. and P.R. wrote and edited the manuscript with feedback from K.M.R. All authors reviewed and approved the final manuscript.

## References

(1) Knowles, T. P. J.; Vendruscolo, M.; Dobson, C. M. The Amyloid State and Its Association with Protein Misfolding Diseases. Nat. Rev. Mol. Cell Biol. 2014, 15 (6), 384–396. 10.1038/nrm3810.

(2) Iadanza, M. G.; Jackson, M. P.; Hewitt, E. W.; Ranson, N. A.; Radford, S. E. A New Era for Understanding Amyloid Structures and Disease. Nat. Rev. Mol. Cell Biol. 2018, 19 (12), 755– 773. 10.1038/s41580-018-0060-8.

(3) Chiti, F.; Dobson, C. M. Protein Misfolding, Amyloid Formation, and Human Disease: A Summary of Progress Over the Last Decade. Annu. Rev. Biochem. 2017, 86 (Volume 86, 2017), 27–68. 10.1146/annurev-biochem-061516-045115.

(4) Zimmet, P. Z. Diabetes and Its Drivers: The Largest Epidemic in Human History? Clin. Diabetes Endocrinol. 2017, 3 (1), 1. 10.1186/s40842-016-0039-3.

(5) Hiddinga, H. J.; Eberhardt, N. L. Intracellular Amyloidogenesis by Human Islet Amyloid Polypeptide Induces Apoptosis in COS-1 Cells. Am. J. Pathol. 1999, 154 (4), 1077–1088. 10.1016/S0002-9440(10)65360-6.

(6) Press, M.; Jung, T.; König, J.; Grune, T.; Höhn, A. Protein Aggregates and Proteostasis in Aging: Amylin and β-Cell Function. Mech. Ageing Dev. 2019, 177, 46–54. 10.1016/j.mad.2018.03.010.

(7) Milardi, D.; Gazit, E.; Radford, S. E.; Xu, Y.; Gallardo, R. U.; Caflisch, A.; Westermark, G. T.; Westermark, P.; Rosa, C. L.; Ramamoorthy, A. Proteostasis of Islet Amyloid Polypeptide: A Molecular Perspective of Risk Factors and Protective Strategies for Type II Diabetes. Chem. Rev. 2021, 121 (3), 1845–1893. 10.1021/acs.chemrev.0c00981.

(8) Westermark, P.; Wernstedt, C.; Wilander, E.; Hayden, D. W.; O’Brien, T. D.; Johnson, K. H. Amyloid Fibrils in Human Insulinoma and Islets of Langerhans of the Diabetic Cat Are Derived from a Neuropeptide-like Protein Also Present in Normal Islet Cells. Proc. Natl. Acad. Sci. 1987, 84 (11), 3881–3885. 10.1073/pnas.84.11.3881.

(9) Roberts, A. N.; Leighton, B.; Todd, J. A.; Cockburn, D.; Schofield, P. N.; Sutton, R.; Holt, S.; Boyd, Y.; Day, A. J.; Foot, E. A. Molecular and Functional Characterization of Amylin, a Peptide Associated with Type 2 Diabetes Mellitus. Proc. Natl. Acad. Sci. 1989, 86 (24), 9662–9666. 10.1073/pnas.86.24.9662.

(10) Jackson, K.; Barisone, G. A.; Diaz, E.; Jin, L.; DeCarli, C.; Despa, F. Amylin Deposition in the Brain: A Second Amyloid in Alzheimer Disease? Ann. Neurol. 2013, 74 (4), 517–526. 10.1002/ana.23956.

(11) Miklossy, J.; Qing, H.; Radenovic, A.; Kis, A.; Vileno, B.; Làszló, F.; Miller, L.; Martins, R. N.; Waeber, G.; Mooser, V.; Bosman, F.; Khalili, K.; Darbinian, N.; McGeer, P. L. Beta Amyloid and Hyperphosphorylated Tau Deposits in the Pancreas in Type 2 Diabetes. Neurobiol. Aging 2010, 31 (9), 1503–1515. 10.1016/j.neurobiolaging.2008.08.019.

(12) Yagihashi, S.; Inaba, W.; Mizukami, H. Dynamic Pathology of Islet Endocrine Cells in Type 2 Diabetes: β-Cell Growth, Death, Regeneration and Their Clinical Implications. J. Diabetes Investig. 2016, 7 (2), 155–165. 10.1111/jdi.12424.

(13) Westermark, P.; Andersson, A.; Westermark, G. T. Islet Amyloid Polypeptide, Islet Amyloid, and Diabetes Mellitus. Physiol. Rev. 2011, 91 (3), 795–826. 10.1152/physrev.00042.2009.

(14) Dai, Z.; Ben-Younis, A.; Vlachaki, A.; Raleigh, D.; Thalassinos, K. Understanding the Structural Dynamics of Human Islet Amyloid Polypeptide: Advancements in and Applications of Ion-Mobility Mass Spectrometry. Biophys. Chem. 2024, 312, 107285. 10.1016/j.bpc.2024.107285.

(15) Rodriguez Camargo, D. C.; Chia, S.; Menzies, J.; Mannini, B.; Meisl, G.; Lundqvist, M.; Pohl, C.; Bernfur, K.; Lattanzi, V.; Habchi, J.; Cohen, S. I.; Knowles, T. P. J.; Vendruscolo, M.; Linse, S. Surface-Catalyzed Secondary Nucleation Dominates the Generation of Toxic IAPP Aggregates. Front. Mol. Biosci. 2021, 8. 10.3389/fmolb.2021.757425.

(16) Abedini, A.; Cao, P.; Plesner, A.; Zhang, J.; He, M.; Derk, J.; Patil, S. A.; Rosario, R.; Lonier, J.; Song, F.; Koh, H.; Li, H.; Raleigh, D. P.; Schmidt, A. M. RAGE Binds Preamyloid IAPP Intermediates and Mediates Pancreatic **β** Cell Proteotoxicity. J. Clin. Invest. 2018, 128 (2), 682–698. 10.1172/JCI85210.

(17) Sakagashira, S.; Sanke, T.; Hanabusa, T.; Shimomura, H.; Ohagi, S.; Kumagaye, K. Y.; Nakajima, K.; Nanjo, K. Missense Mutation of Amylin Gene (S20G) in Japanese NIDDM Patients. Diabetes 1996, 45 (9), 1279–1281. 10.2337/diab.45.9.1279.

(18) Hayakawa, T.; Nagai, Y.; Ando, H.; Yamashita, H.; Takamura, T.; Abe, T.; Nomura, G.; Kobayashi, K. S20G Mutation of the Amylin Gene in Japanese Patients with Type 2 Diabetes. Diabetes Res. Clin. Pract. 2000, 49 (2), 195–197. 10.1016/S0168-8227(00)00142-X.

(19) Sakagashira, S.; Hiddinga, H. J.; Tateishi, K.; Sanke, T.; Hanabusa, T.; Nanjo, K.; Eberhardt, N. L. S20G Mutant Amylin Exhibits Increased *in Vitro* Amyloidogenicity and Increased Intracellular Cytotoxicity Compared to Wild-Type Amylin. Am. J. Pathol. 2000, 157 (6), 2101–2109. 10.1016/S0002-9440(10)64848-1.

(20) Xu, Y.; Maya-Martinez, R.; Guthertz, N.; Heath, G. R.; Manfield, I. W.; Breeze, A. L.; Sobott, F.; Foster, R.; Radford, S. E. Tuning the Rate of Aggregation of hIAPP into Amyloid Using Small-Molecule Modulators of Assembly. Nat. Commun. 2022, 13 (1), 1040. 10.1038/s41467-022-28660-7.

(21) Meier, D. T.; Entrup, L.; Templin, A. T.; Hogan, M. F.; Mellati, M.; Zraika, S.; Hull, R. L.; Kahn, S. E. The S20G Substitution in hIAPP Is More Amyloidogenic and Cytotoxic than Wild-Type hIAPP in Mouse Islets. Diabetologia 2016, 59 (10), 2166–2171. 10.1007/s00125-016-4045-x.

(22) Hassan, S.; White, K.; Terry, C. Linking hIAPP Misfolding and Aggregation with Type 2 Diabetes Mellitus: A Structural Perspective. Biosci. Rep. 2022, 42 (5), BSR20211297. 10.1042/BSR20211297.

(23) Li, X.; Lao, Z.; Zou, Y.; Dong, X.; Li, L.; Wei, G. Mechanistic Insights into the Co-Aggregation of Aβ and hIAPP: An All-Atom Molecular Dynamic Study. J. Phys. Chem. B 2021, 125 (8), 2050–2060. 10.1021/acs.jpcb.0c11132.

(24) Williamson, J. A.; Miranker, A. D. Direct Detection of Transient α-Helical States in Islet Amyloid Polypeptide. Protein Sci. 2007, 16 (1), 110–117. 10.1110/ps.062486907.

(25) Rodriguez Camargo, D. C.; Tripsianes, K.; Buday, K.; Franko, A.; Göbl, C.; Hartlmüller, C.; Sarkar, R.; Aichler, M.; Mettenleiter, G.; Schulz, M.; Böddrich, A.; Erck, C.; Martens, H.; Walch, A. K.; Madl, T.; Wanker, E. E.; Conrad, M.; de Angelis, M. H.; Reif, B. The Redox Environment Triggers Conformational Changes and Aggregation of hIAPP in Type II Diabetes. Sci. Rep. 2017, 7 (1), 44041. 10.1038/srep44041.

(26) Bleiholder, C.; Dupuis, N. F.; Wyttenbach, T.; Bowers, M. T. Ion Mobility–Mass Spectrometry Reveals a Conformational Conversion from Random Assembly to β-Sheet in Amyloid Fibril Formation. Nat. Chem. 2011, 3 (2), 172–177. 10.1038/nchem.945.

(27) Goldsbury, C.; Goldie, K.; Pellaud, J.; Seelig, J.; Frey, P.; Müller, S. A.; Kistler, J.; Cooper, G. J. S.; Aebi, U. Amyloid Fibril Formation from Full-Length and Fragments of Amylin. J. Struct. Biol. 2000, 130 (2), 352–362. 10.1006/jsbi.2000.4268.

(28) Dupuis, N. F.; Wu, C.; Shea, J.-E.; Bowers, M. T. Human Islet Amyloid Polypeptide Monomers Form Ordered β-Hairpins: A Possible Direct Amyloidogenic Precursor. J. Am. Chem. Soc. 2009, 131 (51), 18283–18292. 10.1021/ja903814q.

(29) Sun, Y.; Kakinen, A.; Xing, Y.; Pilkington, E. H.; Davis, T. P.; Ke, P. C.; Ding, F. Nucleation of β-Rich Oligomers and β-Barrels in the Early Aggregation of Human Islet Amyloid Polypeptide. Biochim. Biophys. Acta BBA - Mol. Basis Dis. 2019, 1865 (2), 434–444. 10.1016/j.bbadis.2018.11.021.

(30) Zanjani, A. A. H.; Reynolds, N. P.; Zhang, A.; Schilling, T.; Mezzenga, R.; Berryman, J. T. Amyloid Evolution: Antiparallel Replaced by Parallel. Biophys. J. 2020, 118 (10), 2526–2536. 10.1016/j.bpj.2020.03.023.

(31) Serrano, A. L.; Lomont, J. P.; Tu, L.-H.; Raleigh, D. P.; Zanni, M. T. A Free Energy Barrier Caused by the Refolding of an Oligomeric Intermediate Controls the Lag Time of Amyloid Formation by hIAPP. J. Am. Chem. Soc. 2017, 139 (46), 16748–16758. 10.1021/jacs.7b08830.

(32) Cao, Q.; Boyer, D. R.; Sawaya, M. R.; Abskharon, R.; Saelices, L.; Nguyen, B. A.; Lu, J.; Murray, K. A.; Kandeel, F.; Eisenberg, D. S. Cryo-EM Structures of hIAPP Fibrils Seeded by Patient-Extracted Fibrils Reveal New Polymorphs and Conserved Fibril Cores. Nat. Struct. Mol. Biol. 2021, 28 (9), 724–730. 10.1038/s41594-021-00646-x.

(33) Röder, C.; Kupreichyk, T.; Gremer, L.; Schäfer, L. U.; Pothula, K. R.; Ravelli, R. B. G.; Willbold, D.; Hoyer, W.; Schröder, G. F. Cryo-EM Structure of Islet Amyloid Polypeptide Fibrils Reveals Similarities with Amyloid-β Fibrils. Nat. Struct. Mol. Biol. 2020, 27 (7), 660– 667. 10.1038/s41594-020-0442-4.

(34) Heerde, T.; Rennegarbe, M.; Biedermann, A.; Savran, D.; Pfeiffer, P. B.; Hitzenberger, M.; Baur, J.; Puscalau-Girtu, I.; Zacharias, M.; Schwierz, N.; Haupt, C.; Schmidt, M.; Fändrich, M. Cryo-EM Demonstrates the in Vitro Proliferation of an Ex Vivo Amyloid Fibril Morphology by Seeding. Nat. Commun. 2022, 13 (1), 85. 10.1038/s41467-021-27688-5.

(35) Cao, Q.; Boyer, D. R.; Sawaya, M. R.; Ge, P.; Eisenberg, D. S. Cryo-EM Structure and Inhibitor Design of Human IAPP (Amylin) Fibrils. Nat. Struct. Mol. Biol. 2020, 27 (7), 653– 659. 10.1038/s41594-020-0435-3.

(36) Taylor, A. I. P.; Xu, Y.; Wilkinson, M.; Chakraborty, P.; Brinkworth, A.; Willis, L. F.; Zhuravleva, A.; Ranson, N. A.; Foster, R.; Radford, S. E. Kinetic Steering of Amyloid Formation and Polymorphism by Canagliflozin, a Type-2 Diabetes Drug. J. Am. Chem. Soc. 2025, 147 (14), 11859–11878. 10.1021/jacs.4c16743.

(37) Westermark, P.; Engström, U.; Johnson, K. H.; Westermark, G. T.; Betsholtz, C. Islet Amyloid Polypeptide: Pinpointing Amino Acid Residues Linked to Amyloid Fibril Formation. Proc. Natl. Acad. Sci. 1990, 87 (13), 5036–5040. 10.1073/pnas.87.13.5036.

(38) Saini, R. K.; Goyal, D.; Goyal, B. Targeting Human Islet Amyloid Polypeptide Aggregation and Toxicity in Type 2 Diabetes: An Overview of Peptide-Based Inhibitors. Chem. Res. Toxicol. 2020, 33 (11), 2719–2738. 10.1021/acs.chemrestox.0c00416.

(39) Cao, Q.; Boyer, D. R.; Sawaya, M. R.; Ge, P.; Eisenberg, D. S. Cryo-EM Structure and Inhibitor Design of Human IAPP (Amylin) Fibrils. Nat. Struct. Mol. Biol. 2020, 27 (7), 653– 659. 10.1038/s41594-020-0435-3.

(40) Tang, Y.; Zhang, D.; Zhang, Y.; Liu, Y.; Gong, X.; Chang, Y.; Ren, B.; Zheng, J. Introduction and Fundamentals of Human Islet Amyloid Polypeptide Inhibitors. ACS Appl. Bio Mater. 2020, 3 (12), 8286–8308. 10.1021/acsabm.0c01234.

(41) Susa, A. C.; Wu, C.; Bernstein, S. L.; Dupuis, N. F.; Wang, H.; Raleigh, D. P.; Shea, J.-E.; Bowers, M. T. Defining the Molecular Basis of Amyloid Inhibitors: Human Islet Amyloid Polypeptide–Insulin Interactions. J. Am. Chem. Soc. 2014, 136 (37), 12912–12919. 10.1021/ja504031d.

(42) Bolarinwa, O.; Li, C.; Khadka, N.; Li, Q.; Wang, Y.; Pan, J.; Cai, J. γ-AApeptides–Based Small Molecule Ligands That Disaggregate Human Islet Amyloid Polypeptide. Sci. Rep. 2020, 10 (1), 95. 10.1038/s41598-019-56500-0.

(43) Young, L. M.; Saunders, J. C.; Mahood, R. A.; Revill, C. H.; Foster, R. J.; Tu, L.-H.; Raleigh, D. P.; Radford, S. E.; Ashcroft, A. E. Screening and Classifying Small-Molecule Inhibitors of Amyloid Formation Using Ion Mobility Spectrometry–Mass Spectrometry. Nat. Chem. 2015, 7 (1), 73–81. 10.1038/nchem.2129.

(44) Mahboob, A.; Senevirathne, D. K. L.; Paul, P.; Nabi, F.; Khan, R. H.; Chaari, A. An Investigation into the Potential Action of Polyphenols against Human Islet Amyloid Polypeptide Aggregation in Type 2 Diabetes. Int. J. Biol. Macromol. 2023, 225, 318–350. 10.1016/j.ijbiomac.2022.11.038.

(45) Saravanan, M. S.; Ryazanov, S.; Leonov, A.; Nicolai, J.; Praest, P.; Giese, A.; Winter, R.; Khemtemourian, L.; Griesinger, C.; Killian, J. A. The Small Molecule Inhibitor Anle145c Thermodynamically Traps Human Islet Amyloid Peptide in the Form of Non-Cytotoxic Oligomers. Sci. Rep. 2019, 9 (1), 19023. 10.1038/s41598-019-54919-z.

(46) Dhouafli, Z.; Cuanalo-Contreras, K.; Hayouni, E. A.; Mays, C. E.; Soto, C.; Moreno-Gonzalez, I. Inhibition of Protein Misfolding and Aggregation by Natural Phenolic Compounds. Cell. Mol. Life Sci. 2018, 75 (19), 3521–3538. 10.1007/s00018-018-2872-2.

(47) King, K. M.; Bevan, D. R.; Brown, A. M. Molecular Dynamics Simulations Indicate Aromaticity as a Key Factor in the Inhibition of IAPP(20–29) Aggregation. ACS Chem. Neurosci. 2022, 13 (11), 1615–1626. 10.1021/acschemneuro.2c00025.

(48) Pithadia, A.; Brender, J. R.; Fierke, C. A.; Ramamoorthy, A. Inhibition of IAPP Aggregation and Toxicity by Natural Products and Derivatives. J. Diabetes Res. 2016, 2016 (1), 2046327. 10.1155/2016/2046327.

(49) Lao, Z.; Chen, Y.; Tang, Y.; Wei, G. Molecular Dynamics Simulations Reveal the Inhibitory Mechanism of Dopamine against Human Islet Amyloid Polypeptide (hIAPP) Aggregation and Its Destabilization Effect on hIAPP Protofibrils. ACS Chem. Neurosci. 2019, 10 (9), 4151–4159. 10.1021/acschemneuro.9b00393.

(50) Sang, S.; Lee, M.-J.; Hou, Z.; Ho, C.-T.; Yang, C. S. Stability of Tea Polyphenol (−)-Epigallocatechin-3-Gallate and Formation of Dimers and Epimers under Common Experimental Conditions. J. Agric. Food Chem. 2005, 53 (24), 9478–9484. 10.1021/jf0519055.

(51) Scalvini, B.; Sheikhhassani, V.; van de Brug, N.; Heling, L. W. H. J.; Schmit, J. D.; Mashaghi, A. Circuit Topology Approach for the Comparative Analysis of Intrinsically Disordered Proteins. J. Chem. Inf. Model. 2023, 63 (8), 2586–2602. 10.1021/acs.jcim.3c00391.

(52) Wang, L.; Friesner, R. A.; Berne, B. J. Replica Exchange with Solute Scaling: A More Efficient Version of Replica Exchange with Solute Tempering (REST2). J. Phys. Chem. B 2011, 115 (30), 9431–9438. 10.1021/jp204407d.

(53) Bussi, G. Hamiltonian Replica Exchange in GROMACS: A Flexible Implementation. Mol. Phys. 2014, 112 (3–4), 379–384. 10.1080/00268976.2013.824126.

(54) Sugita, Y.; Okamoto, Y. Replica-Exchange Molecular Dynamics Method for Protein Folding. Chem. Phys. Lett. 1999, 314 (1), 141–151. 10.1016/S0009-2614(99)01123-9.

(55) Robustelli, P.; Piana, S.; Shaw, D. E. Developing a Molecular Dynamics Force Field for Both Folded and Disordered Protein States. Proc. Natl. Acad. Sci. U. S. A. 2018, 115 (21), E4758– E4766. 10.1073/pnas.1800690115.

(56) Pietrucci, F.; Laio, A. A Collective Variable for the Efficient Exploration of Protein Beta-Sheet Structures: Application to SH3 and GB1. J. Chem. Theory Comput. 2009, 5 (9), 2197– 2201. 10.1021/ct900202f.

(57) Shen, Y.; Bax, A. SPARTA+: A Modest Improvement in Empirical NMR Chemical Shift Prediction by Means of an Artificial Neural Network. J. Biomol. NMR 2010, 48 (1), 13–22. 10.1007/s10858-010-9433-9.

(58) Nielsen, J. T.; Mulder, F. A. A. CheSPI: Chemical Shift Secondary Structure Population Inference. J. Biomol. NMR 2021, 75 (6), 273–291. 10.1007/s10858-021-00374-w.

(59) Cesari, A.; Reißer, S.; Bussi, G. Using the Maximum Entropy Principle to Combine Simulations and Solution Experiments. Computation 2018, 6 (1), 15. 10.3390/computation6010015.

(60) Borthakur, K.; Sisk, T. R.; Panei, F. P.; Bonomi, M.; Robustelli, P. Determining Accurate Conformational Ensembles of Intrinsically Disordered Proteins at Atomic Resolution. bioRxiv November 26, 2024, p 2024.10.04.616700. 10.1101/2024.10.04.616700.

(61) Ghafouri, H.; Lazar, T.; Del Conte, A.; Tenorio Ku, L. G.; PED Consortium; Tompa, P.; Tosatto, S. C. E.; Monzon, A. M. PED in 2024: Improving the Community Deposition of Structural Ensembles for Intrinsically Disordered Proteins. Nucleic Acids Res. 2024, 52 (D1), D536–D544. 10.1093/nar/gkad947.

(62) Wang, J.; Wolf, R. M.; Caldwell, J. W.; Kollman, P. A.; Case, D. A. Development and Testing of a General Amber Force Field. J. Comput. Chem. 2004, 25 (9), 1157–1174. 10.1002/jcc.20035.

(63) Robustelli, P.; Stafford, K. A.; Palmer, A. G. I. Interpreting Protein Structural Dynamics from NMR Chemical Shifts. J. Am. Chem. Soc. 2012, 134 (14), 6365–6374. 10.1021/ja300265w.

(64) Robustelli, P.; Ibanez-de-Opakua, A.; Campbell-Bezat, C.; Giordanetto, F.; Becker, S.; Zweckstetter, M.; Pan, A. C.; Shaw, D. E. Molecular Basis of Small-Molecule Binding to α-Synuclein. J. Am. Chem. Soc. 2022, 144 (6), 2501–2510. 10.1021/jacs.1c07591.

(65) Zhu, J.; Salvatella, X.; Robustelli, P. Small Molecules Targeting the Disordered Transactivation Domain of the Androgen Receptor Induce the Formation of Collapsed Helical States. Nat. Commun. 2022, 13 (1), 6390. 10.1038/s41467-022-34077-z.

(66) Zhu, J.; Robustelli, P. Covalent Adducts Formed by the Androgen Receptor Transactivation Domain and Small Molecule Drugs Remain Disordered. J. Chem. Inf. Model. 2025, 65 (12), 6221–6237. 10.1021/acs.jcim.5c00833.

(67) Basu, S.; Martínez-Cristóbal, P.; Frigolé-Vivas, M.; Pesarrodona, M.; Lewis, M.; Szulc, E.; Bañuelos, C. A.; Sánchez-Zarzalejo, C.; Bielskutė, S.; Zhu, J.; Pombo-García, K.; Garcia-Cabau, C.; Zodi, L.; Dockx, H.; Smak, J.; Kaur, H.; Batlle, C.; Mateos, B.; Biesaga, M., Escobedo, A.; Bardia, L.; Verdaguer, X.; Ruffoni, A.; Mawji, N. R.; Wang, J.; Obst, J. K.; Tam, T.; Brun-Heath, I.; Ventura, S.; Meierhofer, D.; García, J.; Robustelli, P.; Stracker, T. H.; Sadar, M. D.; Riera, A.; Hnisz, D.; Salvatella, X. Rational Optimization of a Transcription Factor Activation Domain Inhibitor. Nat. Struct. Mol. Biol. 2023, 30 (12), 1958–1969. 10.1038/s41594-023-01159-5.

(68) Páll, S.; Zhmurov, A.; Bauer, P.; Abraham, M.; Lundborg, M.; Gray, A.; Hess, B.; Lindahl, E. Heterogeneous Parallelization and Acceleration of Molecular Dynamics Simulations in GROMACS. J. Chem. Phys. 2020, 153 (13), 134110. 10.1063/5.0018516.

(69) Abraham, M. J.; Murtola, T.; Schulz, R.; Páll, S.; Smith, J. C.; Hess, B.; Lindahl, E. GROMACS: High Performance Molecular Simulations through Multi-Level Parallelism from Laptops to Supercomputers. SoftwareX 2015, 1–2, 19–25. 10.1016/j.softx.2015.06.001.

(70) Tribello, G. A.; Bonomi, M.; Branduardi, D.; Camilloni, C.; Bussi, G. PLUMED 2: New Feathers for an Old Bird. Comput. Phys. Commun. 2014, 185 (2), 604–613. 10.1016/j.cpc.2013.09.018.

(71) Nanga, R. P. R.; Brender, J. R.; Vivekanandan, S.; Ramamoorthy, A. Structure and Membrane Orientation of IAPP in Its Natively Amidated Form at Physiological pH in a Membrane Environment. Biochim. Biophys. Acta BBA - Biomembr. 2011, 1808 (10), 2337–2342. 10.1016/j.bbamem.2011.06.012.

(72) Bussi, G.; Donadio, D.; Parrinello, M. Canonical Sampling through Velocity Rescaling. J. Chem. Phys. 2007, 126 (1), 014101. 10.1063/1.2408420.

(73) Hess, B.; Bekker, H.; Berendsen, H. J. C.; Fraaije, J. G. E. M. LINCS: A Linear Constraint Solver for Molecular Simulations. J. Comput. Chem. 1997, 18 (12), 1463–1472. 10.1002/(SICI)1096-987X(199709)18:12<1463::AID-JCC4>3.0.CO;2-H.

(74) Nosé, S.; Klein, M. L. Constant Pressure Molecular Dynamics for Molecular Systems. Mol. Phys. 1983, 50 (5), 1055–1076. 10.1080/00268978300102851.

(75) Parrinello, M.; Rahman, A. Polymorphic Transitions in Single Crystals: A New Molecular Dynamics Method. J. Appl. Phys. 1981, 52 (12), 7182–7190. 10.1063/1.328693.

76. (76) O’Boyle, N. M.; Banck, M.; James, C. A.; Morley, C.; Vandermeersch, T.; Hutchison, G. R. Open Babel: An Open Chemical Toolbox. J. Cheminformatics 2011, 3 (1), 33. 10.1186/1758-2946-3-33.

(77) Sousa da Silva, A. W.; Vranken, W. F. ACPYPE - AnteChamber PYthon Parser interfacE. BMC Res. Notes 2012, 5 (1), 367. 10.1186/1756-0500-5-367.

(78) McGibbon, R. T.; Beauchamp, K. A.; Harrigan, M. P.; Klein, C.; Swails, J. M.; Hernández, C. X.; Schwantes, C. R.; Wang, L.-P.; Lane, T. J.; Pande, V. S. MDTraj: A Modern Open Library for the Analysis of Molecular Dynamics Trajectories. Biophys. J. 2015, 109 (8), 1528–1532. 10.1016/j.bpj.2015.08.015.

(79) Pedregosa, F.; Varoquaux, G.; Gramfort, A.; Michel, V.; Thirion, B.; Grisel, O.; Blondel, M.; Prettenhofer, P.; Weiss, R.; Dubourg, V.; Vanderplas, J.; Passos, A.; Cournapeau, D.; Brucher, M.; Perrot, M.; Duchesnay, É. Scikit-Learn: Machine Learning in Python. J. Mach. Learn. Res. 2011, 12 (85), 2825–2830.

(80) Kohlhoff, K. J.; Robustelli, P.; Cavalli, A.; Salvatella, X.; Vendruscolo, M. Fast and Accurate Predictions of Protein NMR Chemical Shifts from Interatomic Distances. J. Am. Chem. Soc. 2009, 131 (39), 13894–13895. 10.1021/ja903772t.

(81) Flyvbjerg, H.; Petersen, H. G. Error Estimates on Averages of Correlated Data. J. Chem. Phys. 1989, 91 (1), 461–466. 10.1063/1.457480.

(82) Wolff, U. Monte Carlo Errors with Less Errors. Comput. Phys. Commun. 2004, 156 (2), 143–153. 10.1016/S0010-4655(03)00467-3.

