## Supplementary Information for "Monomer binding modes of small molecules that modulate the kinetics of hIAPP amyloid formation"

Paul Robustelli

### REST2 MD simulation convergence analyses

We ran unbiased all-atom explicit solvent replica exchange solute tempering[1–3] (REST2) MD simulations of apo WT hIAPP, apo S20G hIAPP, WT hIAPP in the presence of small molecule inhibitor YX-I-1, S20G hIAPP in the presence of YX-I-1, WT hIAPP in the presence of small molecule accelerator YX-A-1, and S20G hIAPP in the presence of YX-A-1. REST2 MD simulations were performed with the a99SB-*disp* protein force field and a99SB-*disp* water model using a solute temperature ladder spanning 300-500 K. Small molecules were parametrized using the GAFF1 force field.

We evaluated the time course and ensemble averages of simulated conformational properties in the temperature replicas and demultiplexed replicas, which follow the time evolution of independent replicas through the solute temperature ladder, to assess the convergence of each REST2 simulation (Figure S1-S2, S10-S13). To assess simulation convergence we compared the populations of intramolecular contacts ("contact maps"), secondary structure propensities, distributions of the radius of gyration ($R_g$) and distributions of the value of the α-helical order parameter Sα[4] in each temperature and demultiplexed replica. Sα measures the similarity of all 7-residue segments to an ideal helical structure ($\phi$=-57,$\psi$=-47) calculated according $S\alpha = \sum_i^N \frac{1-\left(\frac{\mathrm{RMSD}\alpha_i}{r_0}\right)^8}{1-\left(\frac{\mathrm{RMSD}\alpha_i}{r_0}\right)^{12}}$. We discarded the first microsecond of REST2 simulations of all systems due to the slow equilibration of Sα (Figure S3). Unless otherwise indicated, all analyses are reported for the truncated 300 K temperature of each REST2 MD simulation. Figures S1-S2 and S10-S13 show that the secondary structure propensities of WT and S20G hIAPP are highly consistent for all 20 replicas in the demultiplexed trajectories, demonstrating that each demultiplexed replica is independently sampling similar distributions of conformational space. Additional convergence analyses of each system can be found in accompanying GitHub repository (https://github.com/paulrobustelli/Garcia_hIAPP_monomer_binders_2025).

### Analysis of circuit topology assignments

Circuit topology assignments were made for all frames of the 300K replicas of each simulated system following the procedure of Scalvini et al. 2023[5]. To obtain circuit topology assignments, intramolecular contacts between residues were defined using 10 Å distance cutoff between any pair of atoms. In each frame, each pair of intramolecular contacts (consisting of 4 contact sites) was assigned a circuit topology relationship based on the location of the contacts within the protein sequence, as defined in Figure S5. We assigned topological relations of contact pairs the following numerical values: series=1, concerted series=2, parallel=3, parallel$^{-1}$=4, concerted parallel=5, concerted parallel$^{-1}$ =6, and crossing=7, and constructed a matrix of circuit topology assignments for each contact pair. Circuit topology matrices reflect the topological relationship of all unique $\binom{n}{2}$ (n choose 2) possible contact pairs where *n* is the number of protein residues. Here *n* = 37 and $\binom{n}{2} = 666$. Circuit topology matrices for an ensemble with *N* frames therefore have a dimension of $\left[\binom{n}{2}, \binom{n}{2}, N\right]$ ,and have a dimension of [666, 666, *N*] in this work. To reduce redundancy, we analyze only the top half of the circuit topology matrices. After assigning the circuit topology relations for the 300 K replicas of all trajectories, we merged all circuit topologies matrices into a single matrix and performed principal component analysis (PCA) to reduce to dimensionality of the merged matrix into two principal components for all simulated systems using the scikit-learn package[6–8]. We subsequently used the k-means algorithm to cluster the reduced topological space.

### Determination of k-means clusters

We performed k-means clustering on the 2D latent space resulting from PCA of the merged circuit topology matrices, scanning from k=2 to k=20 clusters, and compared several clustering metrics to assess the quality of clustering assignments (Figure S7). Specifically, we compared the distance between each point in circuit topology latent space and the cluster centroid (the distortion), silhouette scores, the Calinski-Harbasz index and the Davies-Bouldin index for each number of clusters. Briefly, the silhouette score measures the similarity between datapoints assigned to one cluster versus other clusters, the Calinski-Harbasz index measures the ratio of between-cluster variance to within-cluster variance and the Davies-Bouldin index measures the similarity between each cluster and its most similar cluster. As hIAPP populates a highly heterogeneous conformational ensemble in all ensembles and each cluster consists of broad distribution of conformations, no clearly optimal cluster assignment is consistently identified by all clustering metrics. Each metric identifies locally optimal solutions between 3-10 clusters. For each of these cluster assignments, we compared the conformational properties of each cluster and identified six clusters as a good compromise between identifying a tractable number of states with distinct conformational properties for further analysis (Figures S8-S9, Figures S19-S22). We note that cluster 6, is lowly populated (< 0.1% of conformations) in all systems except for S20G + YX-A-1 (P=9.9%).

### Supplementary Tables

**Supplementary Table 1: Simulation times for apo and holo REST2 MD simulations of WT and S20G hIAPP.** All simulations used 20 solute temperature rungs spanning 300 – 500 K. Total simulation times are reported in parentheses. The first 1μs of each simulation was discarded as equilibration, and all subsequent analyses are reported on the truncated simulations. Aggregate simulation times of truncated trajectories are reported.

| System | μs/ replica | Aggregate simulation time for truncated simulations (μs) |
|---|---|---|
| **WT hIAPP Apo** | 3.3 (4.3) | 66 |
| **S20G hIAPP Apo** | 2.8 (3.8) | 56 |
| **WT hIAPP + YX-I-1** | 2.4 (3.4) | 48 |
| **S20G hIAPP + YX-I-1** | 1.8 (2.8) | 36 |
| **WT hIAPP + YX-A-1** | 1.7 (2.7) | 34 |
| **S20G hIAPP + YX-A-1** | 1.8 (2.7) | 36 |

**Supplementary Table 2: Populations of conformational states identified by k-means clustering of the 2D circuit topology latent space**. The populations of each cluster obtained from k-means clustering with k=6 clusters observed in the 300 K temperature replica of REST2 MD simulations. Cluster assignments were made from a merged trajectory containing all ensembles.

| | WT hIAPP Apo (%) | S20G hIAPP Apo (%) | WT hIAPP + YX-I-1 (%) | S20G hIAPP + YX-I-1 (%) | WT hIAPP + YX-A-1 (%) | S20G hIAPP + YX-A-1 (%) |
|---|---|---|---|---|---|---|
| C1 | 54.2 | 43.2 | 42.8 | 51.2 | 43.0 | 48.7 |
| C2 | 16.1 | 21.5 | 24.0 | 20.7 | 20.7 | 10.5 |
| C3 | 16.8 | 15.3 | 14.7 | 15.1 | 21.4 | 20.3 |
| C4 | 4.6 | 5.0 | 15.2 | 6.1 | 11.4 | 5.4 |
| C5 | 8.3 | 15.1 | 3.3 | 6.9 | 3.6 | 5.3 |
| C6 | 0.01 (6 frames) | 0.003 (1 frame) | 0.0* | 0.0* | 0.01 (3 frames) | 9.9 |

*No conformations assigned to this cluster.

**Supplementary Table 3: Ligand-bound fraction of conformational states identified by k-means clustering on the 2D circuit topology latent space.** The ligand-bound fraction of each cluster obtained from k-means clustering with k=6 clusters observed in the 300 K temperature replica of REST2 MD simulations. Cluster assignments were made from a merged trajectory containing all ensembles. Statistical error estimates were computed using a blocking analysis.

| Cluster | WT + YX-I-1 (%) | S20G + YX-I-1 (%) | WT + YX-A-1 (%) | S20G + YX-A-1 (%) |
|---|---|---|---|---|
| 1 | 75.2 ± 2.4 | 72.52 ± 2.1 | 51.4 ± 2.25 | 53.62 ± 2.95 |
| 2 | 58.3 ± 5.6 | 72.35 ± 6.21 | 52.53 ± 4.42 | 58.12 ± 4.42 |
| 3 | 77.8 ± 3.3 | 75.0 ± 3.18 | 48.01 ± 3.48 | 58.53 ± 4.25 |
| 4 | 80.2 ± 10.0 | 67.31 ± 5.95 | 73.14 ± 20.7 | 74.77 ± 5.73 |
| 5 | 55.1 ± 8.4 | 67.02 ± 6.68 | 46.48 ± 18.8 | 59.63 ± 10.23 |
| 6 | * | * | 66.7 ± 33.33 | 46.68 ± 12.43 |

*No conformations were assigned to this cluster.

### Supplemental Figures

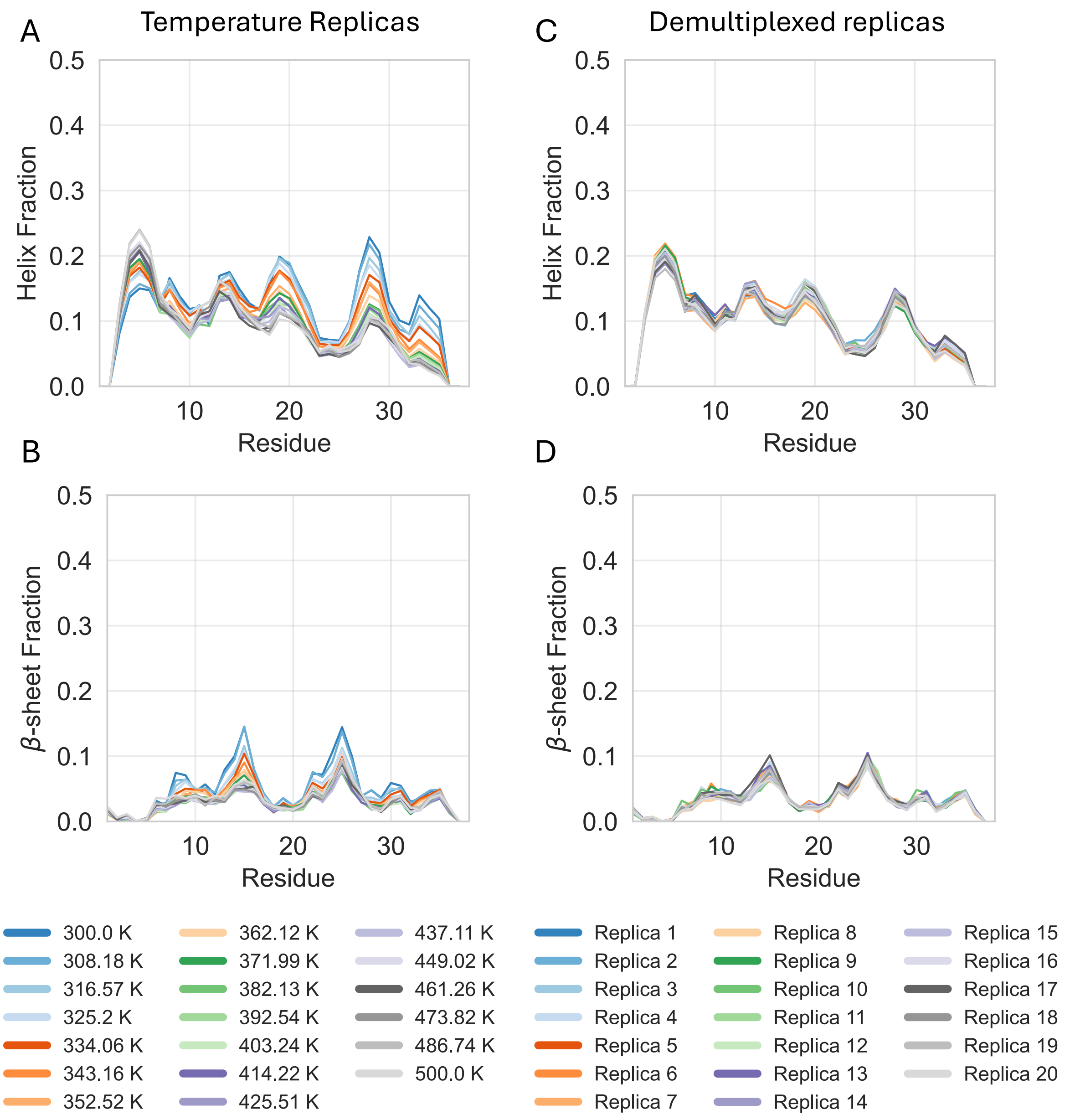


**Supplementary Figure 1. Secondary structure populations observed in a REST2 MD simulation of apo WT hIAPP.** Comparison of α-helical and β-sheet propensities of WT hIAPP apo observed in the 20 solute temperature rungs spanning 300-500 K (A, B) and 20 demultiplexed replicas (C, D). The first microsecond of data was discarded as equilibration, yielding 3.3 μs per replica for analysis. Secondary structure content was calculated with the DSSP algorithm.

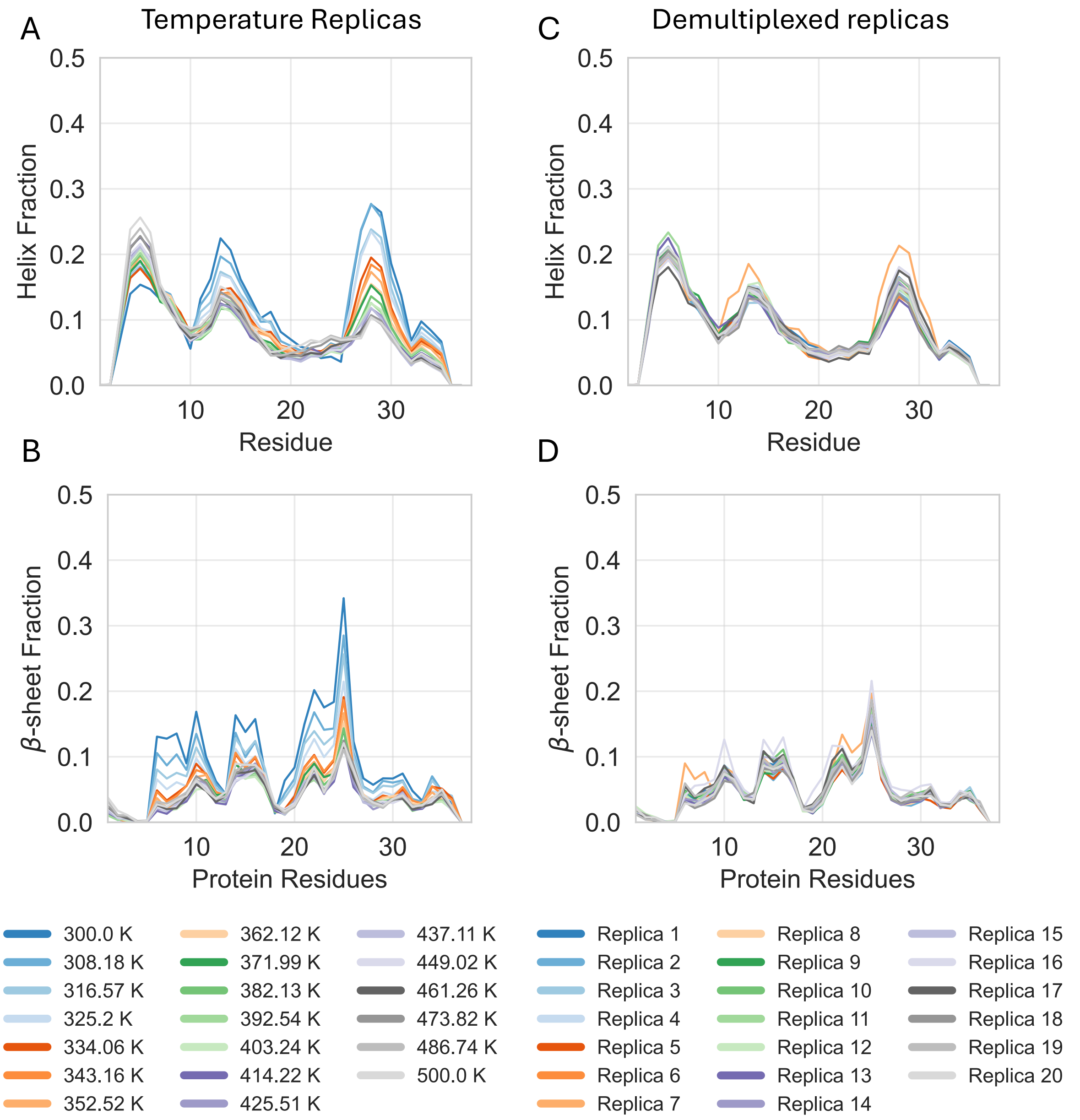


**Supplementary Figure 2. Secondary structure populations observed in a REST2 MD simulation of apo S20G hIAPP.** Comparison of α-helical and β-sheet propensities of S20G hIAPP apo observed in the 20 solute temperature rungs spanning 300-500 K (A, B) and 20 demultiplexed replicas (C, D). The first microsecond of data was discarded as equilibration, yielding 2.8 μs per replica for analysis. Secondary structure content was calculated with the DSSP algorithm.

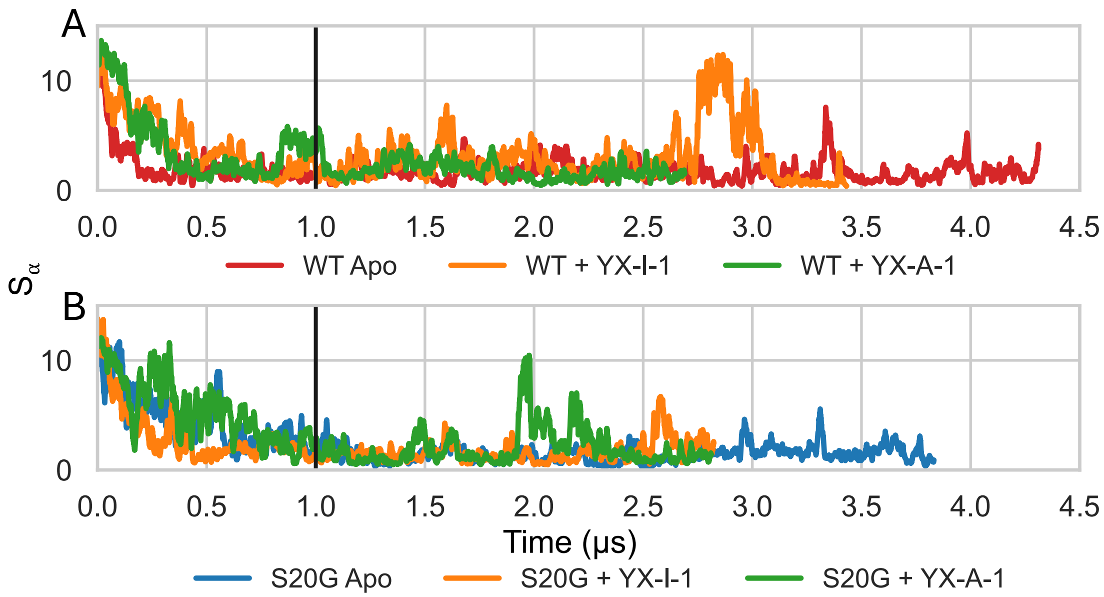


**Supplementary Figure 3. Time evolution of the α-helical order parameter Sα in the 300 K replica of REST2 MD simulations.** Comparison of Sα time series in the 300 K replicas of WT hIAPP (A) and S20G hIAPP (B) before discarding the first microsecond of each simulation. A horizontal black line is drawn at the first microsecond of the simulation. The data is smoothed by using window averaging with a sliding window of size N=100.

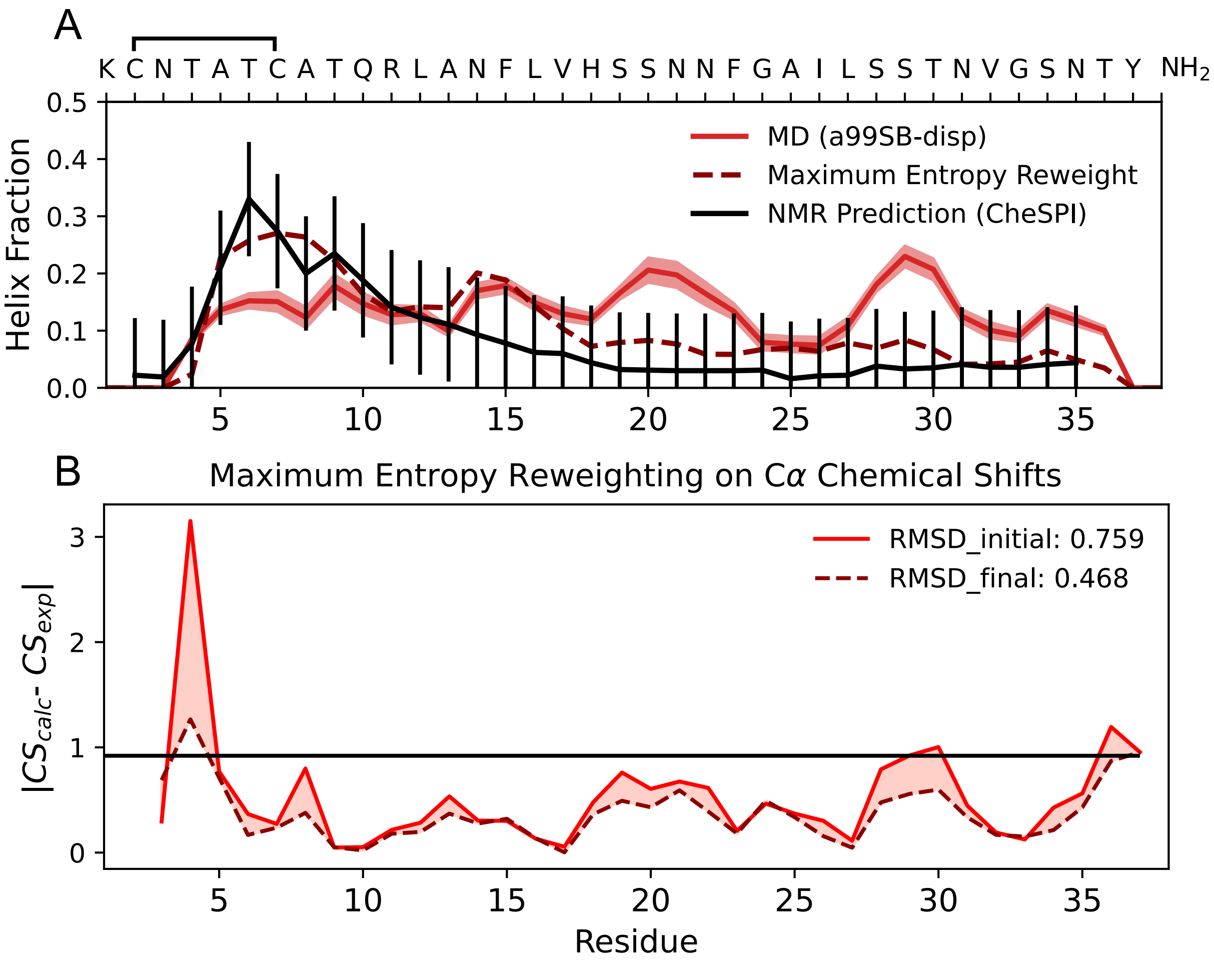


**Supplementary Figure 4. Comparison of experimental and calculated helix populations and NMR Cα chemical shifts of the 300 K replica of the apo WT hIAPP REST2 simulation. (**A) Comparison of the per-residue helix fractions of unbiased MD (solid red line) and maximum entropy reweighted MD ensembles (dashed maroon line) of apo WT hIAPP with per-residue helix fractions predicted from experimental NMR chemical shifts using the CheSPI[9] algorithm (black line). CheSPI values are shown with an estimated ± 10% error. Statistical error estimates of helix fractions calculated from unbiased MD computed by blocking are shown as shaded regions. (B) The per-residue deviation between experimental NMR Cα chemical shifts computed by SPARTA+[10] from unbiased MD (solid red line) and maximum entropy reweighted MD ensembles (dashed maroon line) of apo WT hIAPP. The solid black line indicates the estimated Cα prediction error of SPARTA+ (0.92 ppm).

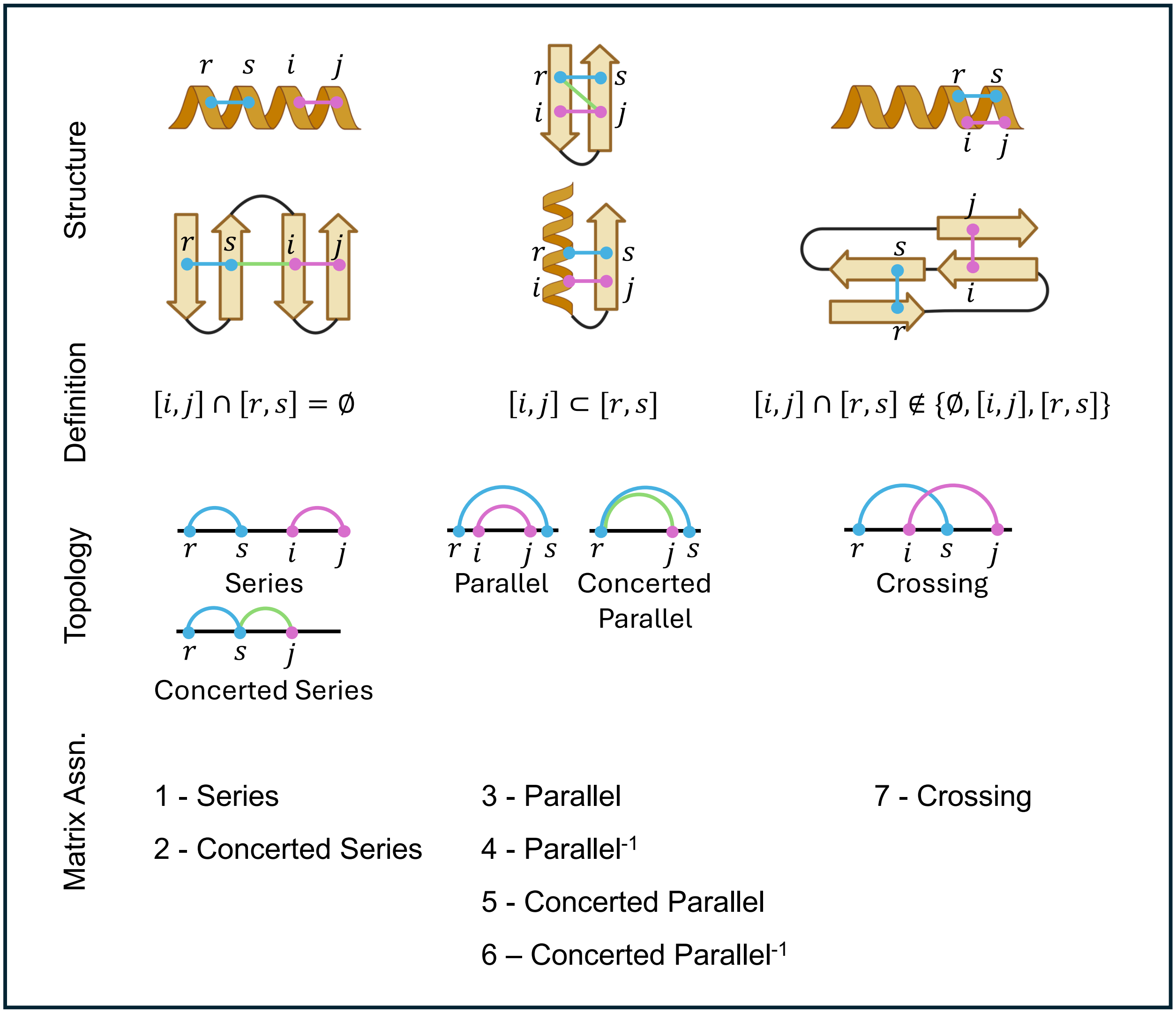


**Supplementary Figure 5. Comparison of elementary circuit topology assignments**. Illustrations of circuit topology relations of intramolecular contacts within protein structures defined in Scalvini et al. 2023[5]. Contact pairs are denoted by blue, green, or pink lines. For each circuit topology relationship of contact pairs observed in hIAPP conformations analyzed here, we indicate the numerical values assigned in circuit topology matrices. The magnitude of the value increases with the complexity of the circuit topology assignments.

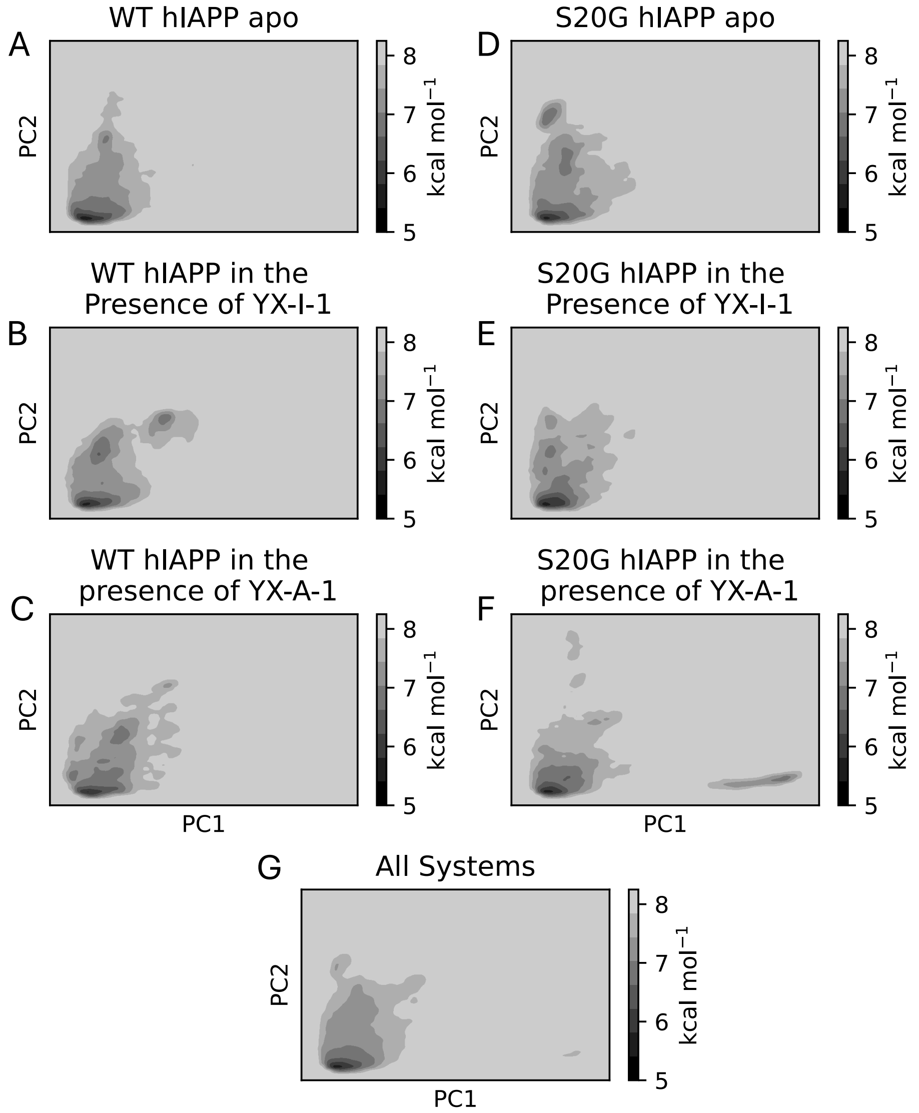


**Supplementary Figure 6. Free energy surfaces of apo and ligand-binding simulations of WT and S20G hIAPP projected onto a latent space derived from circuit topology assignments.** Free energy surfaces of hIAPP conformations sampled by WT hIAPP in its apo state (A), WT hIAPP in the presence of YX-I-1 (B), WT hIAPP in the presence of YX-A-1 (C), S20G hIAPP in its apo state (D), S20G hIAPP in the presence of YX-I-1 (E), S20G hIAPP in the presence of YX-A-1 (F) and in a merged ensemble obtained from all simulations (G) projected onto the 2D latent space derived from circuit topology assignments. Relative free energies were determined according to $\Delta G = -K_B T \ln(P)$ where $P$ is the population obtained using a 50 x 50 2D histogram grid defined by the two principal components (PCs) of the circuit topology latent space.

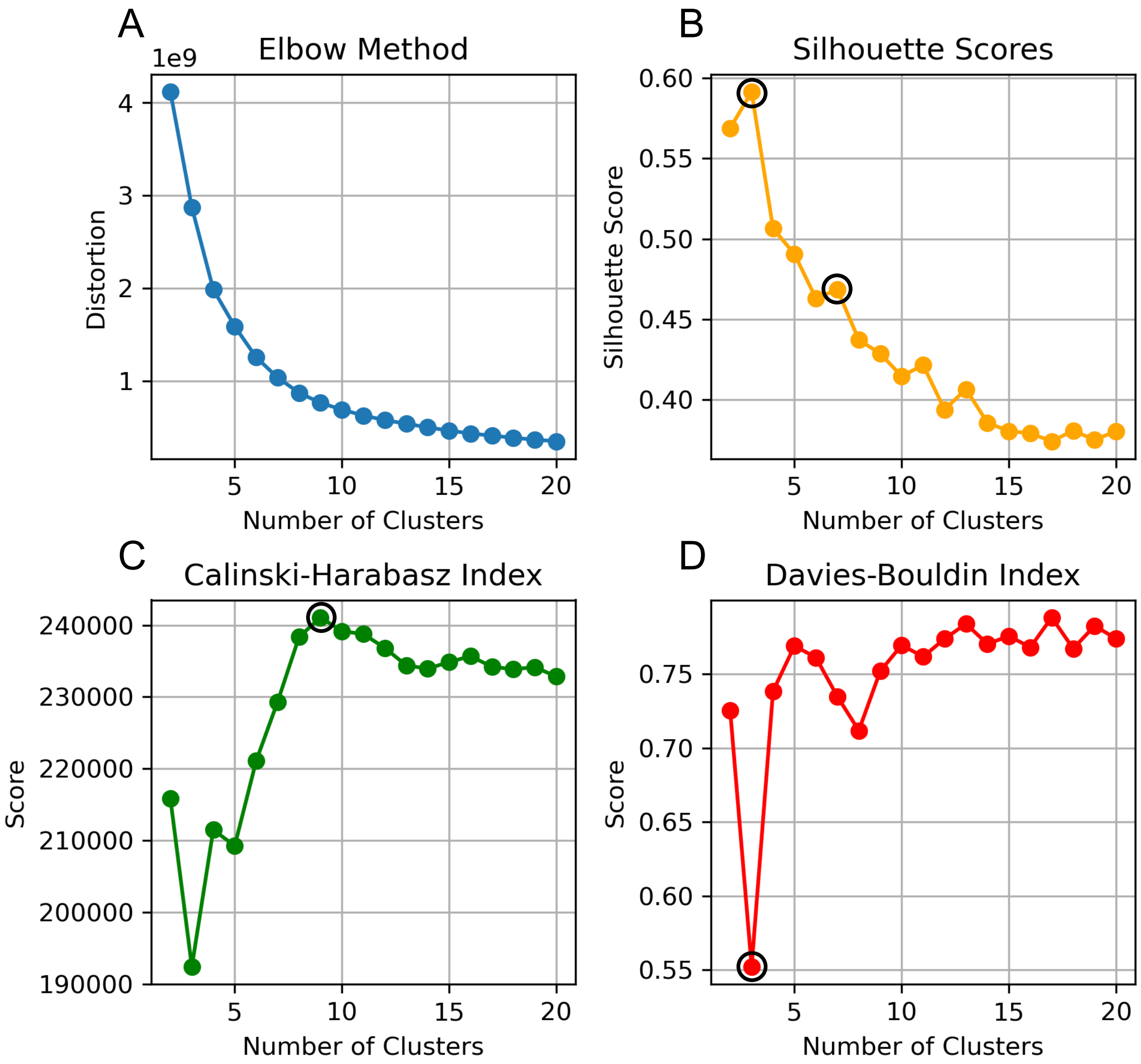


**Supplementary Figure 7. Comparison of clustering metrics for k-means clustering of hIAPP conformations.** Evaluation of k-means clustering assignments of hIAPP conformations projected onto the reduced-dimensionality 2D circuit topology latent space with varying numbers of clusters (k = 2–20). Shown are (A) distortion, quantifying the average distance of points from their cluster centroids, (B) silhouette score, (C) Calinski–Harabasz index, and (D) Davies–Bouldin index. Locally optimal scores for each metric are indicated by circles.

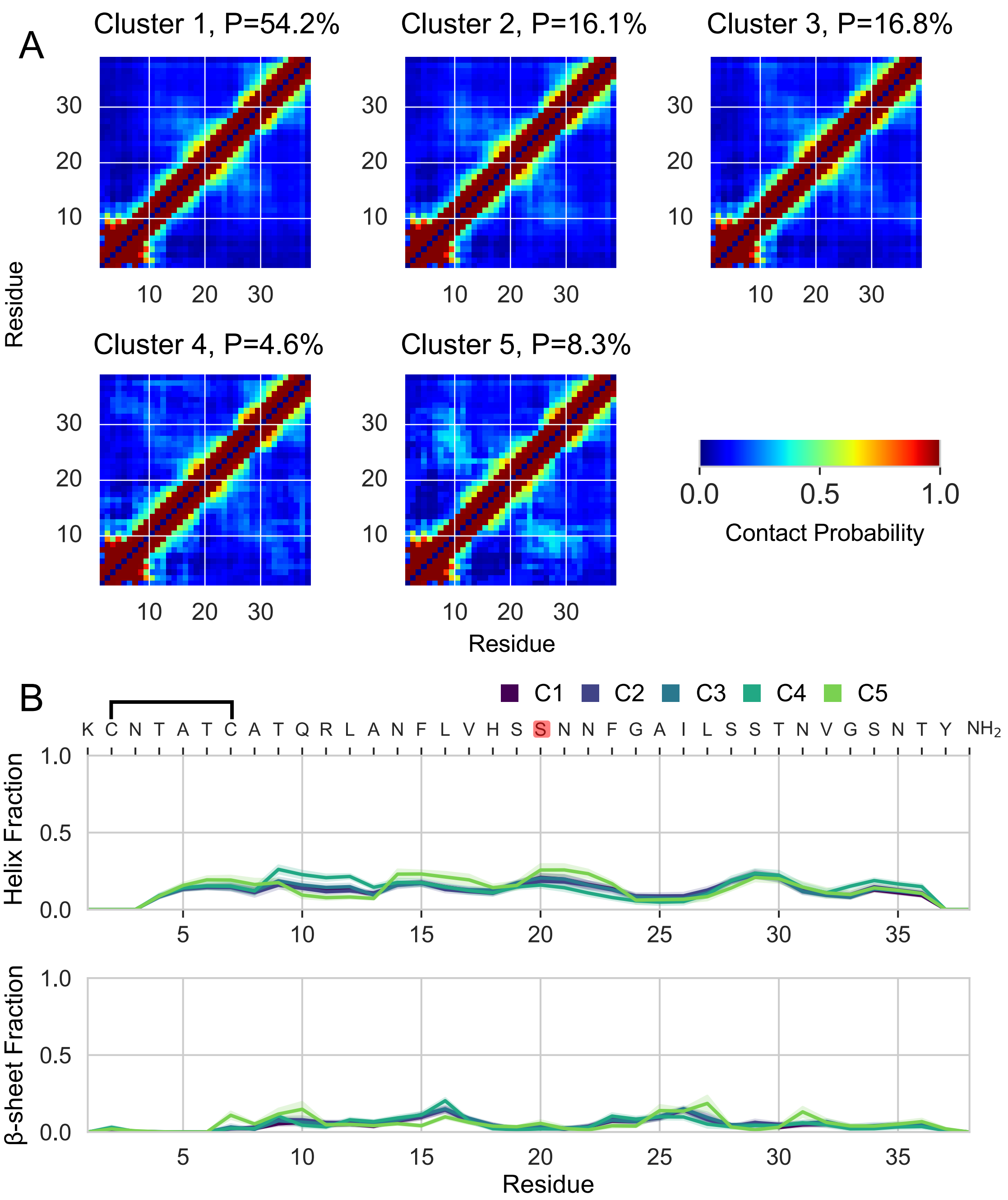


**Supplementary Figure 8. Conformational properties of apo WT hIAPP.** Populations of (A) intramolecular contacts and (B) secondary structure elements in hIAPP conformations identified by k-means clustering (k = 6) from the 300 K replica of a REST2 MD simulation of apo WT hIAPP. Residue–residue contacts are defined by an 8 Å heavy-atom distance cutoff. Secondary structure propensities were assigned by DSSP, with statistical error estimates from blocking shown as shaded regions. Cluster 6 was excluded due to insufficient population (6 frames).

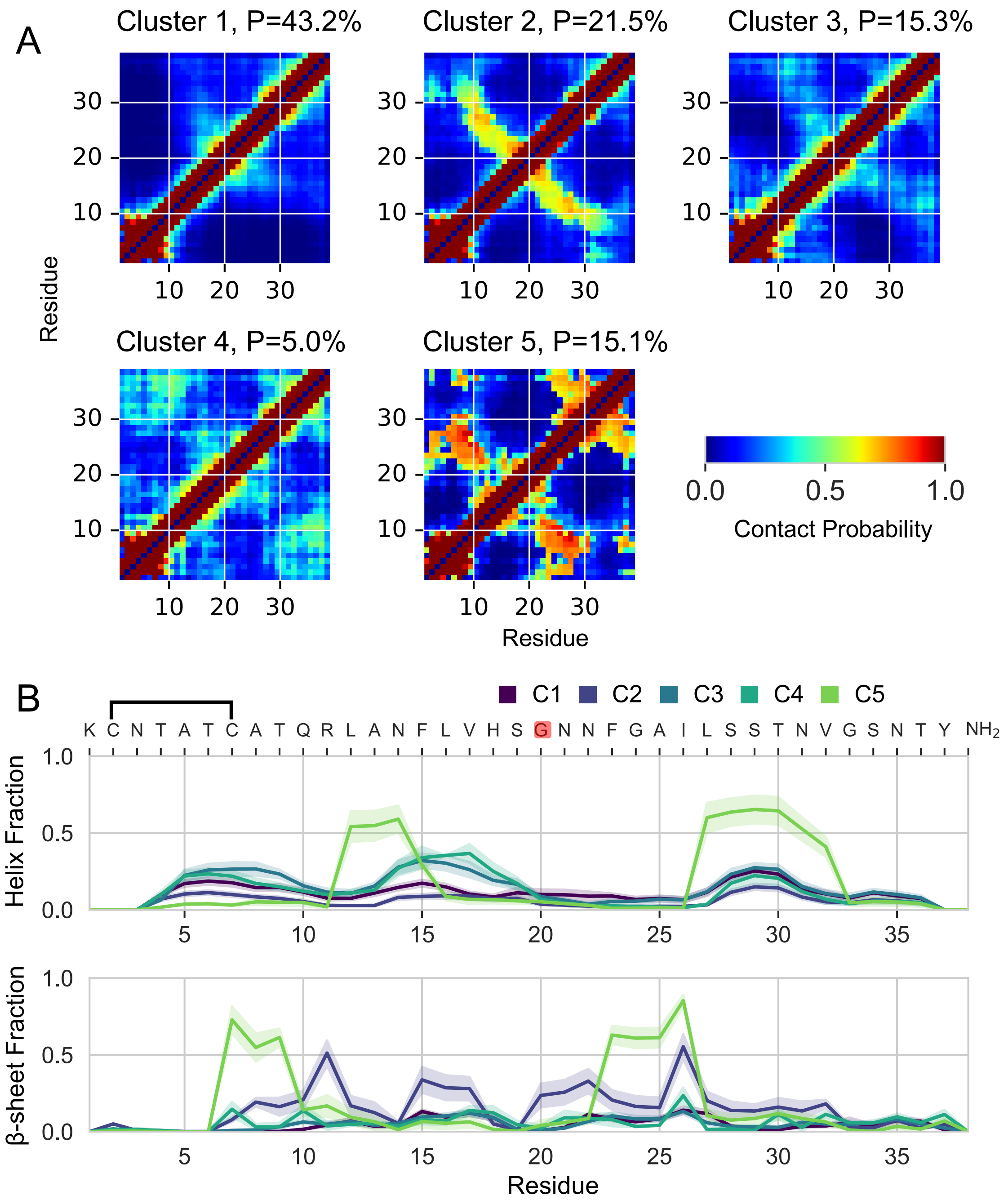


**Supplementary Figure 9. Conformational properties of apo S20G hIAPP.** Populations of (A) intramolecular contacts and (B) secondary structure elements in hIAPP conformations identified by k-means clustering (k = 6) from the 300 K replica of a REST2 MD simulation of apo S20G hIAPP. Residue–residue contacts are defined by an 8 Å heavy-atom distance cutoff. Secondary structure propensities were assigned by DSSP, with statistical error estimates from blocking shown as shaded regions. Cluster 6 was excluded due to insufficient population (1 frame).

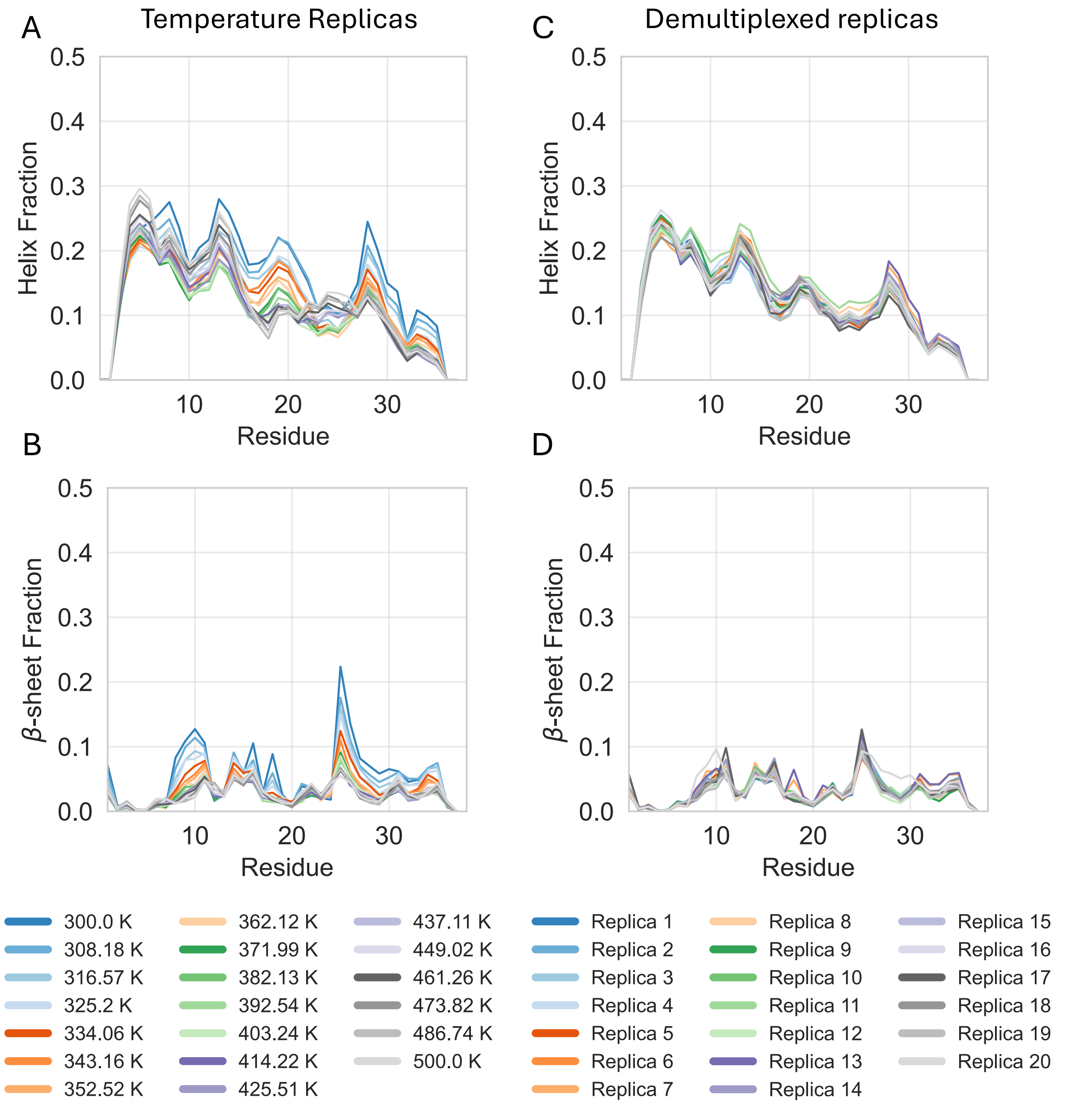


**Supplementary Figure 10. Secondary structure populations observed in a REST2 MD simulation of WT hIAPP in the presence of YX-I-1.** Comparison of α-helical and β-sheet propensities of WT hIAPP in the presence of YX-I-1 observed in the 20 solute temperature rungs spanning 300-500 K (A, B) and 20 demultiplexed replicas (C, D). The first microsecond of data was discarded as equilibration, yielding 2.4 μs per replica for analysis. Secondary structure content was calculated with the DSSP algorithm.

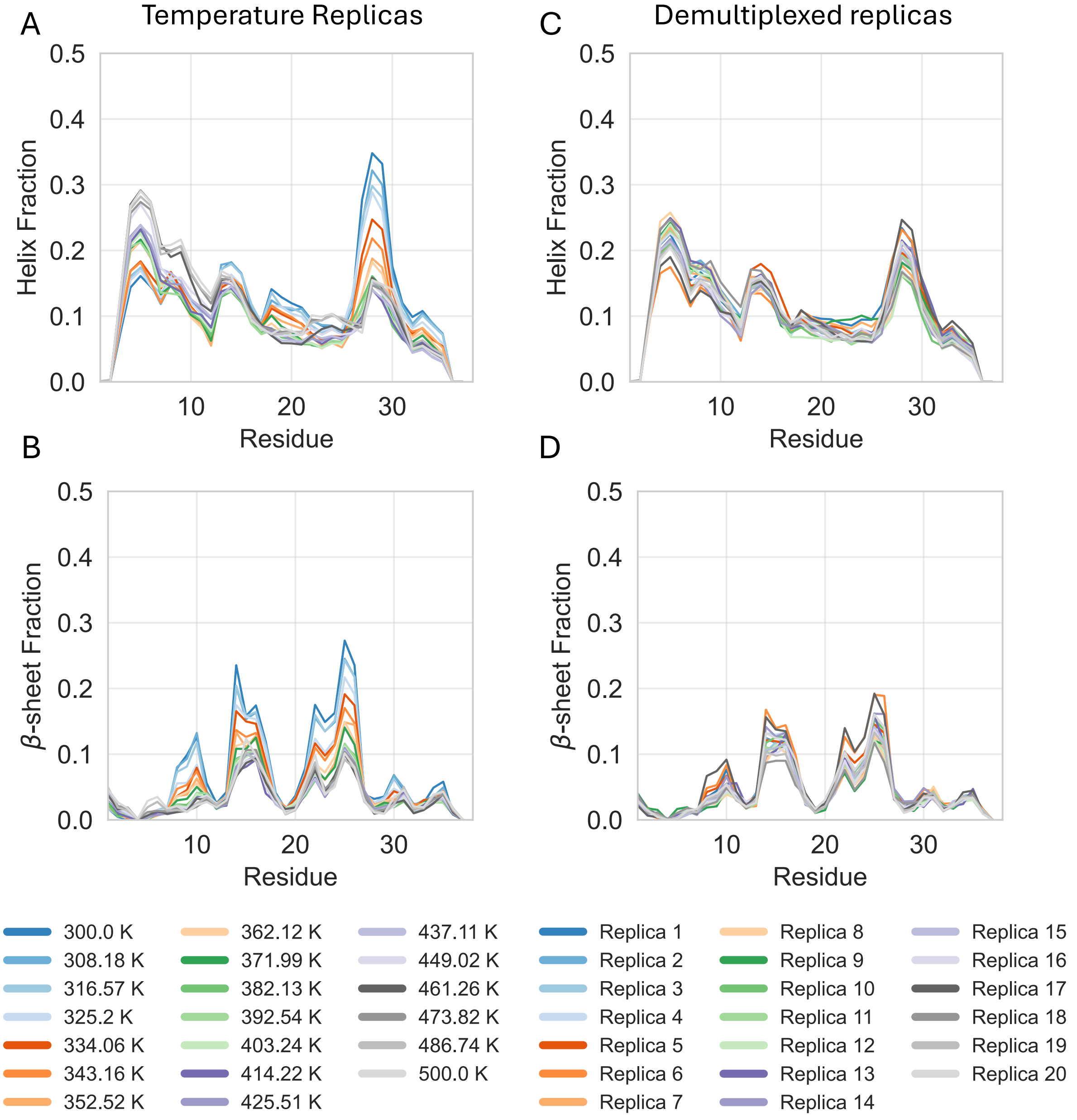


**Supplementary Figure 11. Secondary structure populations observed in a REST2 MD simulation of S20G hIAPP in the presence of YX-I-1.** Comparison of α-helical and β-sheet propensities of S20G hIAPP in the presence of YX-I-1 observed in the 20 solute temperature rungs spanning 300-500 K (A, B) and 20 demultiplexed replicas (C, D). The first microsecond of data was discarded as equilibration, yielding 1.8 μs per replica for analysis. Secondary structure content was calculated with the DSSP algorithm.

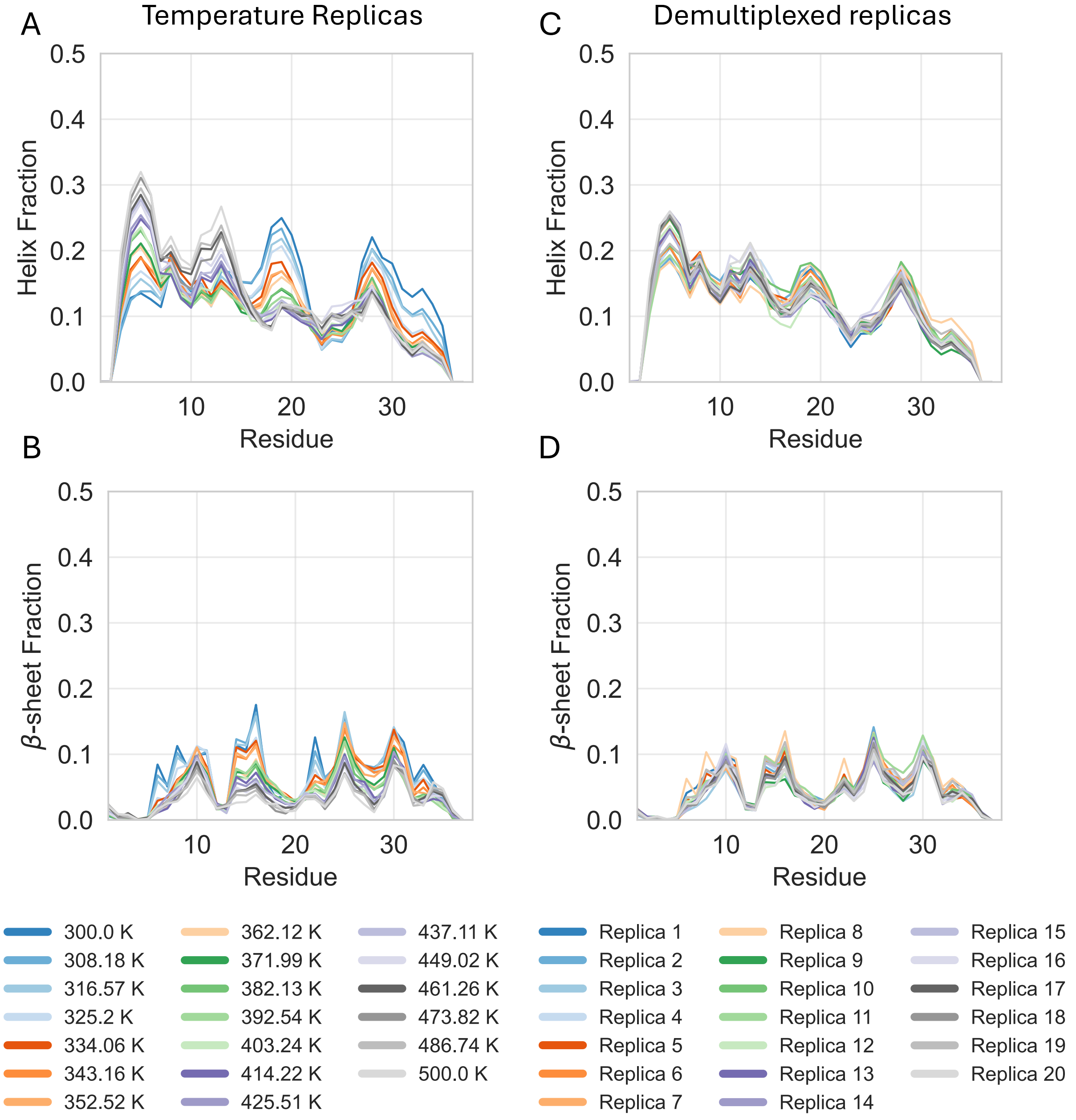


**Supplementary Figure 12. Secondary structure populations observed in a REST2 MD simulation of WT hIAPP in the presence of YX-A-1.** Comparison of α-helical and β-sheet propensities of WT hIAPP in the presence of YX-A-1 observed in the 20 solute temperature rungs spanning 300-500 K (A, B) and 20 demultiplexed replicas (C, D). The first microsecond of data was discarded as equilibration, yielding 1.7 μs per replica for analysis. Secondary structure content was calculated with the DSSP algorithm.

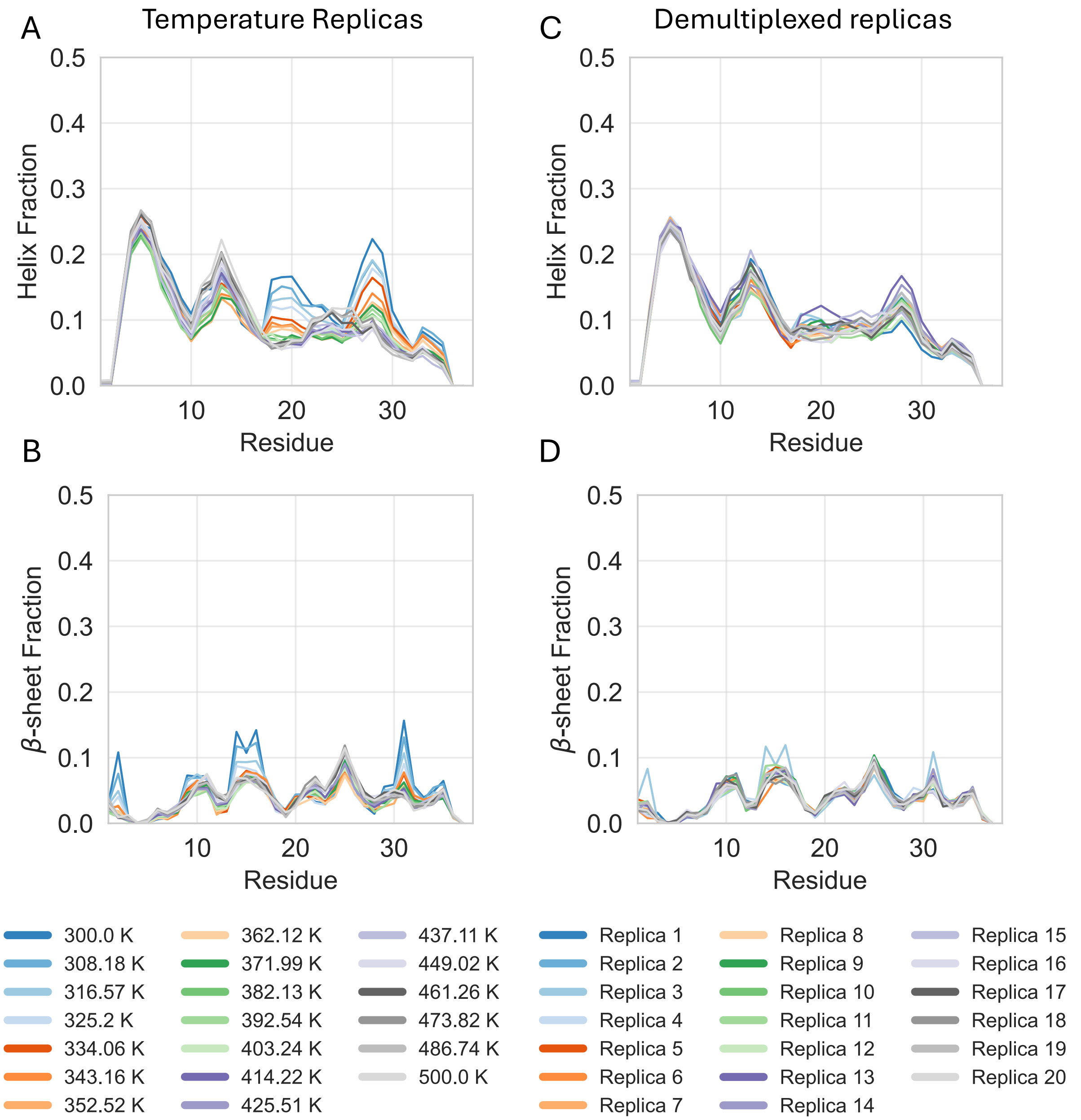


**Supplementary Figure 13. Secondary structure populations observed in a REST2 MD simulation of S20G hIAPP in the presence of YX-A-1.** Comparison of α-helical and β-sheet propensities of S20G hIAPP in the presence of YX-A-1 observed in the 20 solute temperature rungs spanning 300-500 K (A, B) and 20 demultiplexed replicas (C, D). The first microsecond of data was discarded as equilibration, yielding 1.8 μs per replica for analysis. Secondary structure content was calculated with the DSSP algorithm.

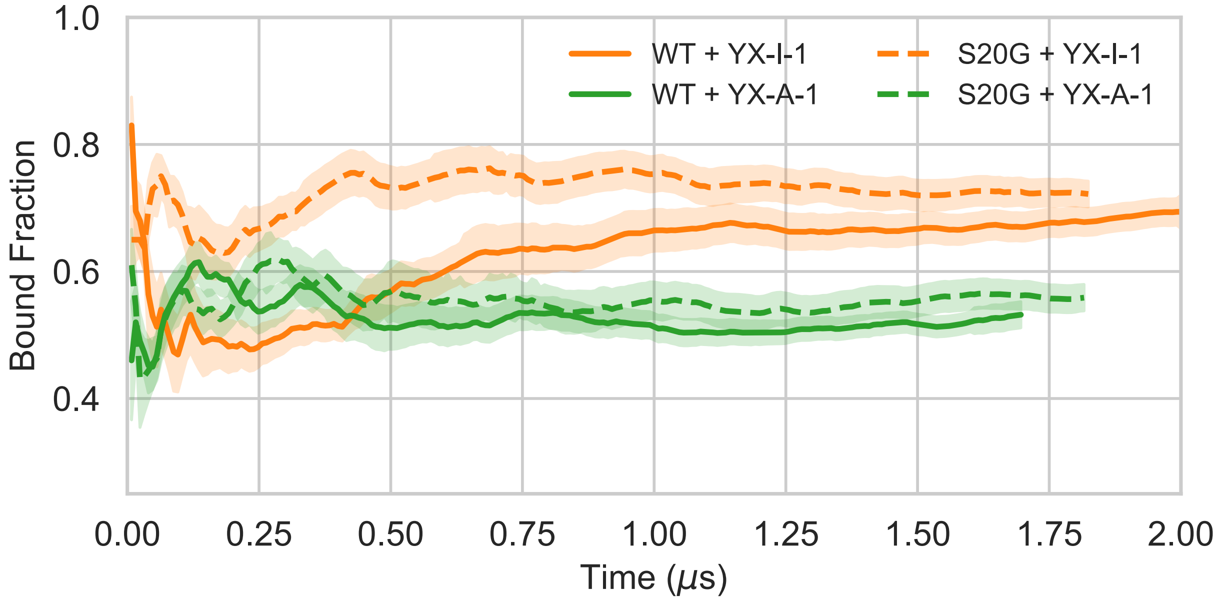


**Supplementary Figure 14. Time evolution of bound fraction in ligand-binding simulations of WT hIAPP and S20G hIAPP.** Bound fractions of ligands as a function of simulation time, with the first microsecond of each simulation discarded. Simulations of WT hIAPP are shown as solid lines and those of S20G hIAPP as dashed lines. Binding simulations of YX-I-1 are shown in orange and binding simulations of YX-A-1 are shown in green. Bound frames were defined using a 6 Å distance cutoff between the closest heavy atoms of the protein and ligand.

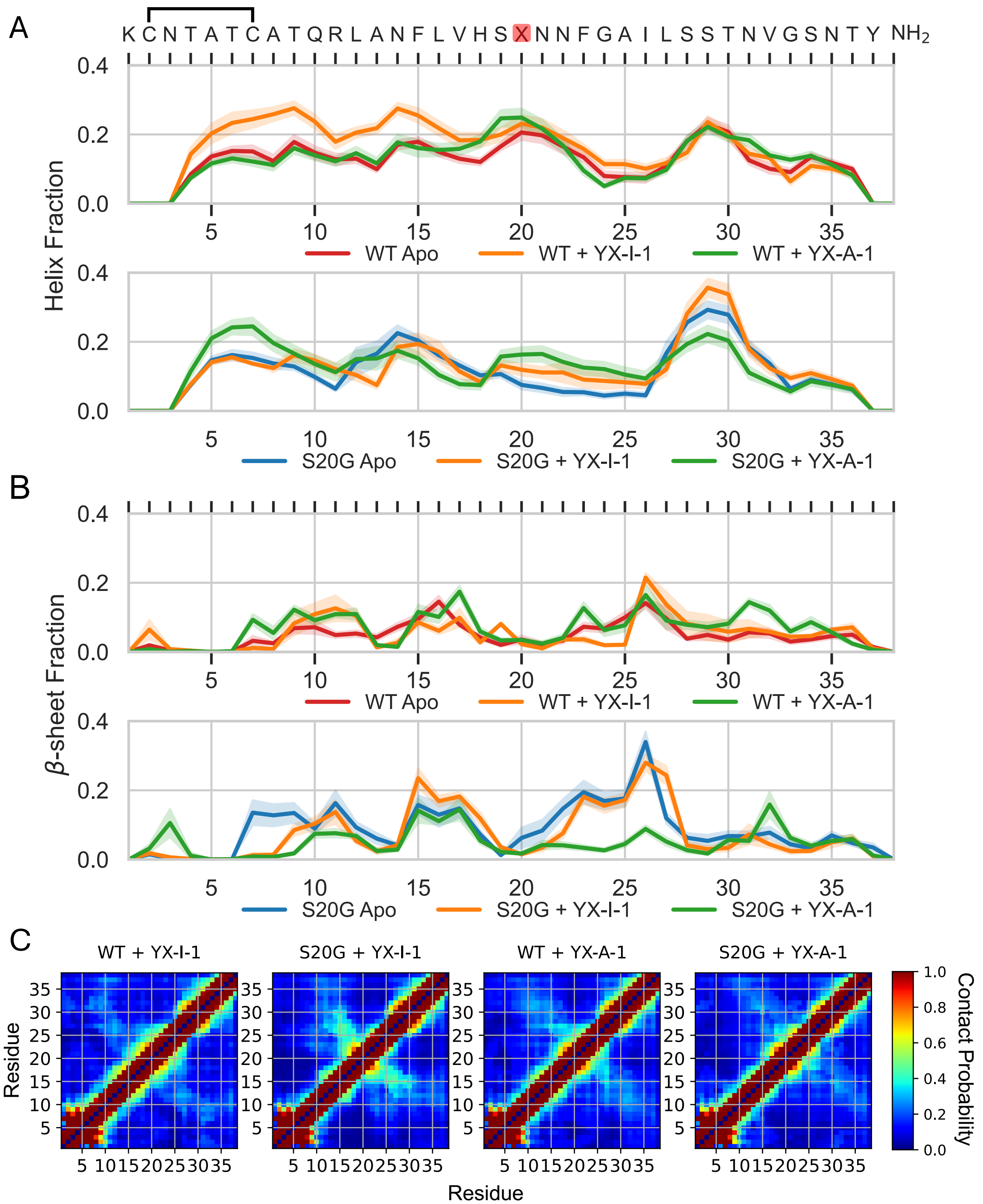


**Supplementary Figure 15. Conformational properties of WT and S20G hIAPP in ligand-binding simulations.** Populations of (A) helical and (B) β-sheet secondary structure elements, and (C) intramolecular contacts. Secondary structure propensities were assigned by DSSP, with statistical error estimates from blocking shown as shaded regions. Residue–residue contacts were defined by an 8 Å heavy-atom distance cutoff.

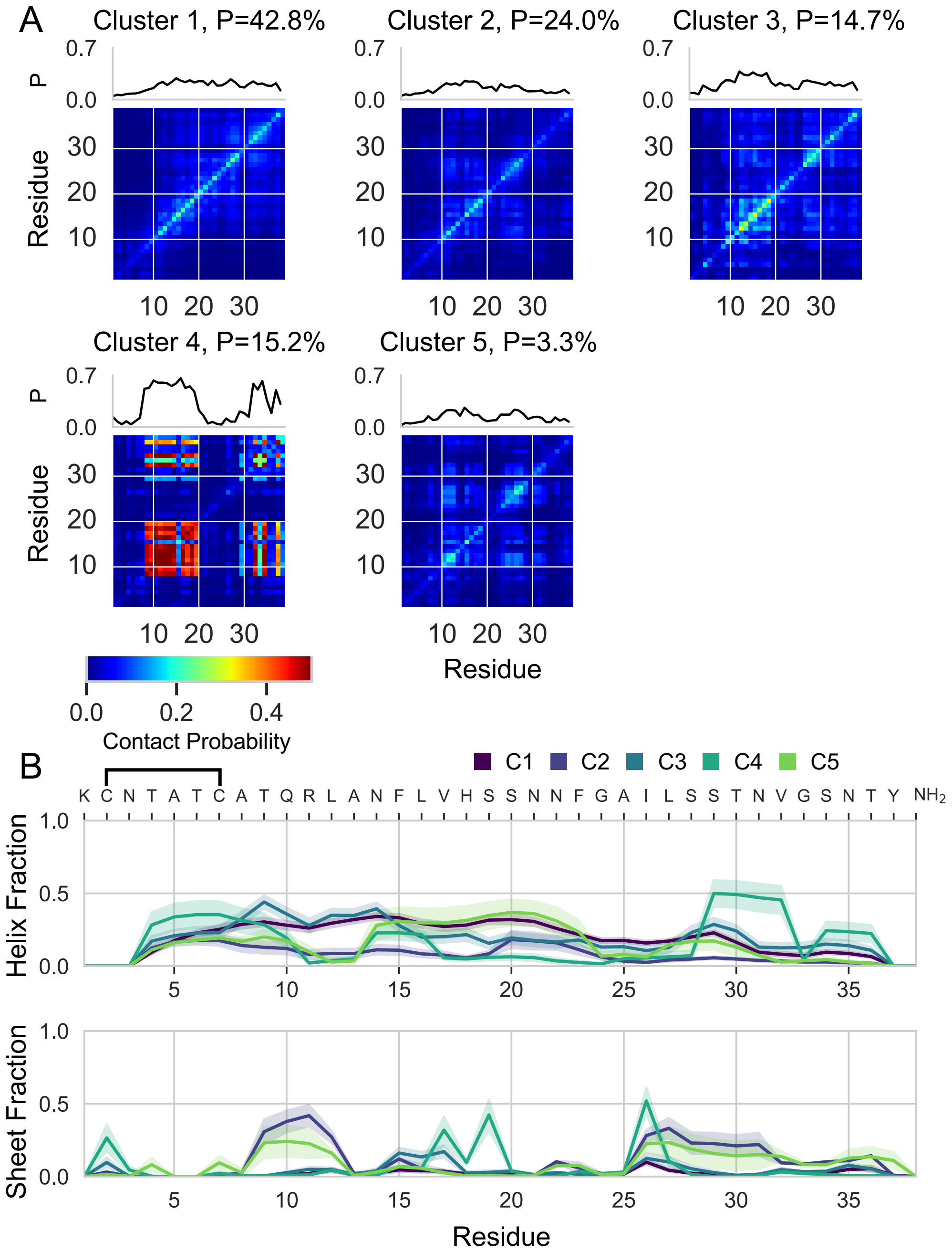


**Supplementary Figure 16. Ligand interactions and secondary structure populations of conformational states of WT hIAPP in the presence of YX-I-1.** (A) The probability that each pair of hIAPP residues simultaneously forms a contact with YX-I-1 and (B) the secondary structure populations in conformational states identified by k-means clustering with k = 6 clusters from the 300K replica of a REST2 MD simulation of WT hIAPP in the presence of YX-I-1. Secondary structure propensities were assigned by DSSP assignments and statistical error estimates from blocking are shown as shaded regions. No frames of this trajectory were assigned to cluster 6.

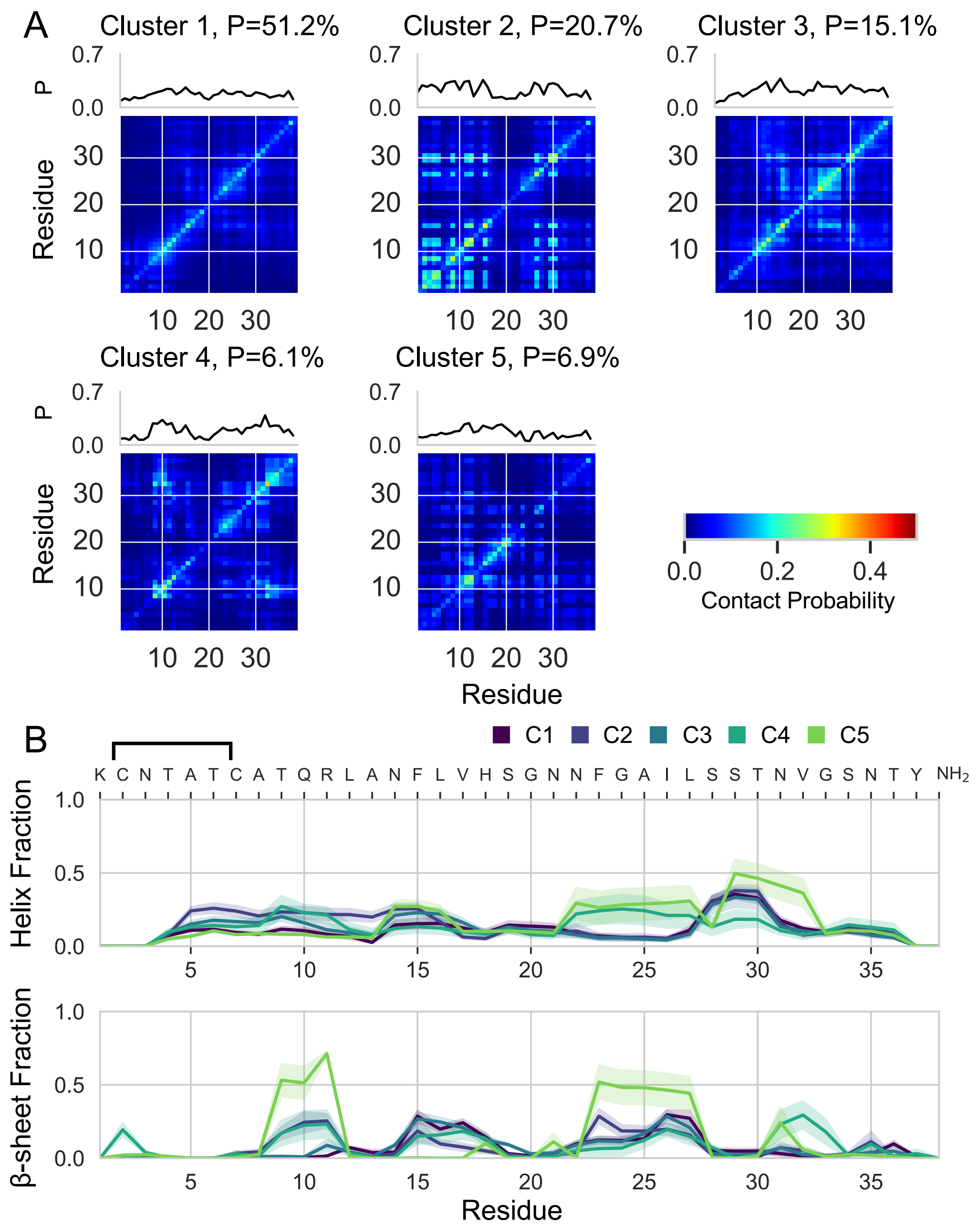


**Supplementary Figure 17. Ligand interactions and secondary structure populations of conformational states of S20G hIAPP in the presence of YX-I-1.** (A) The probability that each pair of hIAPP residues simultaneously forms a contact with YX-I-1 and (B) the secondary structure populations in conformational states identified by k-means clustering with k = 6 clusters from the 300K replica of a REST2 MD simulation of S20G hIAPP in the presence of YX-I-1. Secondary structure propensities were assigned by DSSP assignments and statistical error estimates from blocking are shown as shaded regions. No frames of this trajectory were assigned to cluster 6.

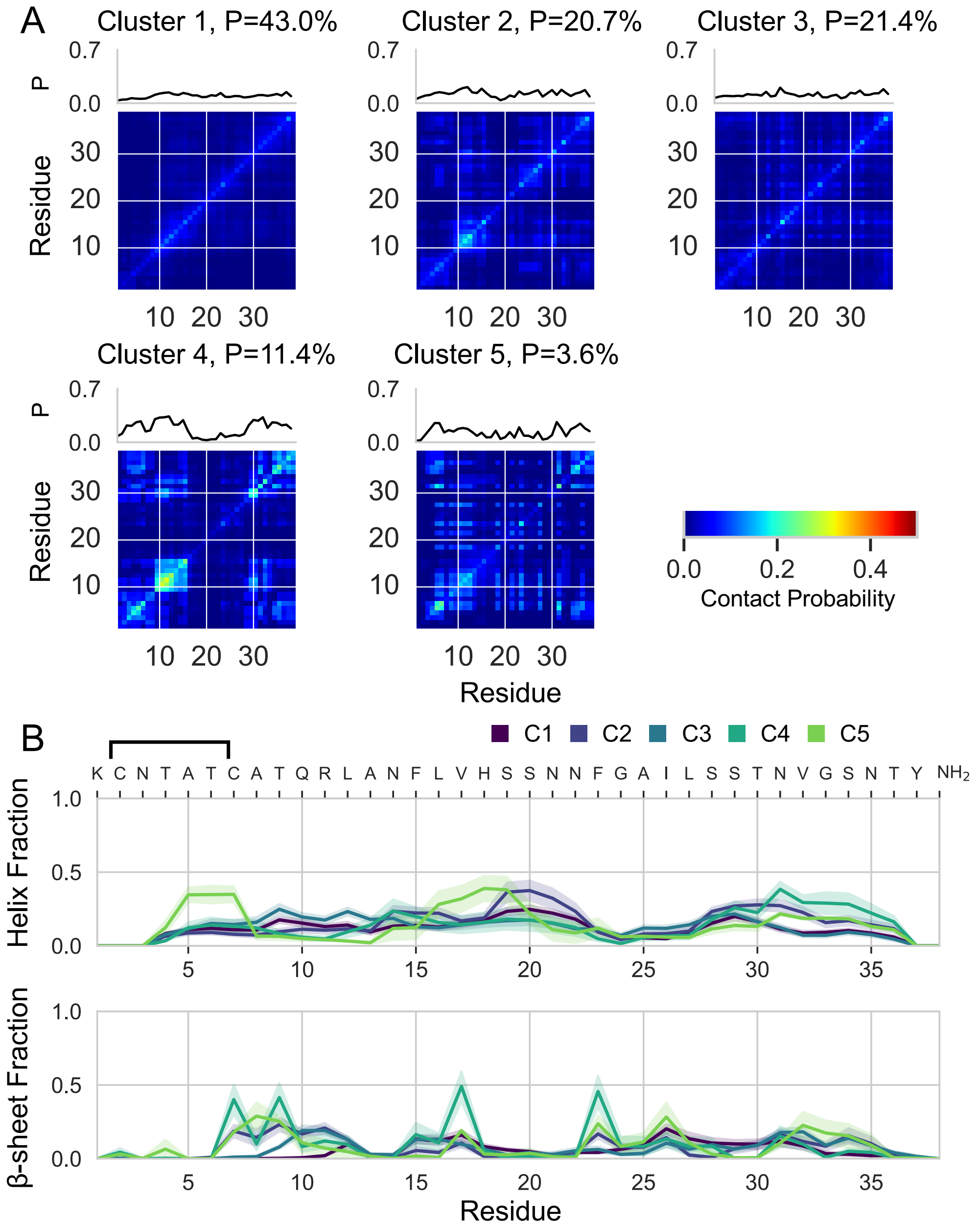


**Supplementary Figure 18. Ligand interactions and secondary structure populations of conformational states of WT hIAPP in the presence of YX-A-1.** (A) The probability that each pair of hIAPP residues simultaneously forms a contact with YX-A-1 and (B) the secondary structure populations in conformational states identified by k-means clustering with k = 6 clusters from the 300K replica of a REST2 MD simulation of WT hIAPP in the presence of YX-A-1. Secondary structure propensities were assigned by DSSP assignments and statistical error estimates from blocking are shown as shaded regions. Cluster 6 was excluded due to insufficient population (3 frames).

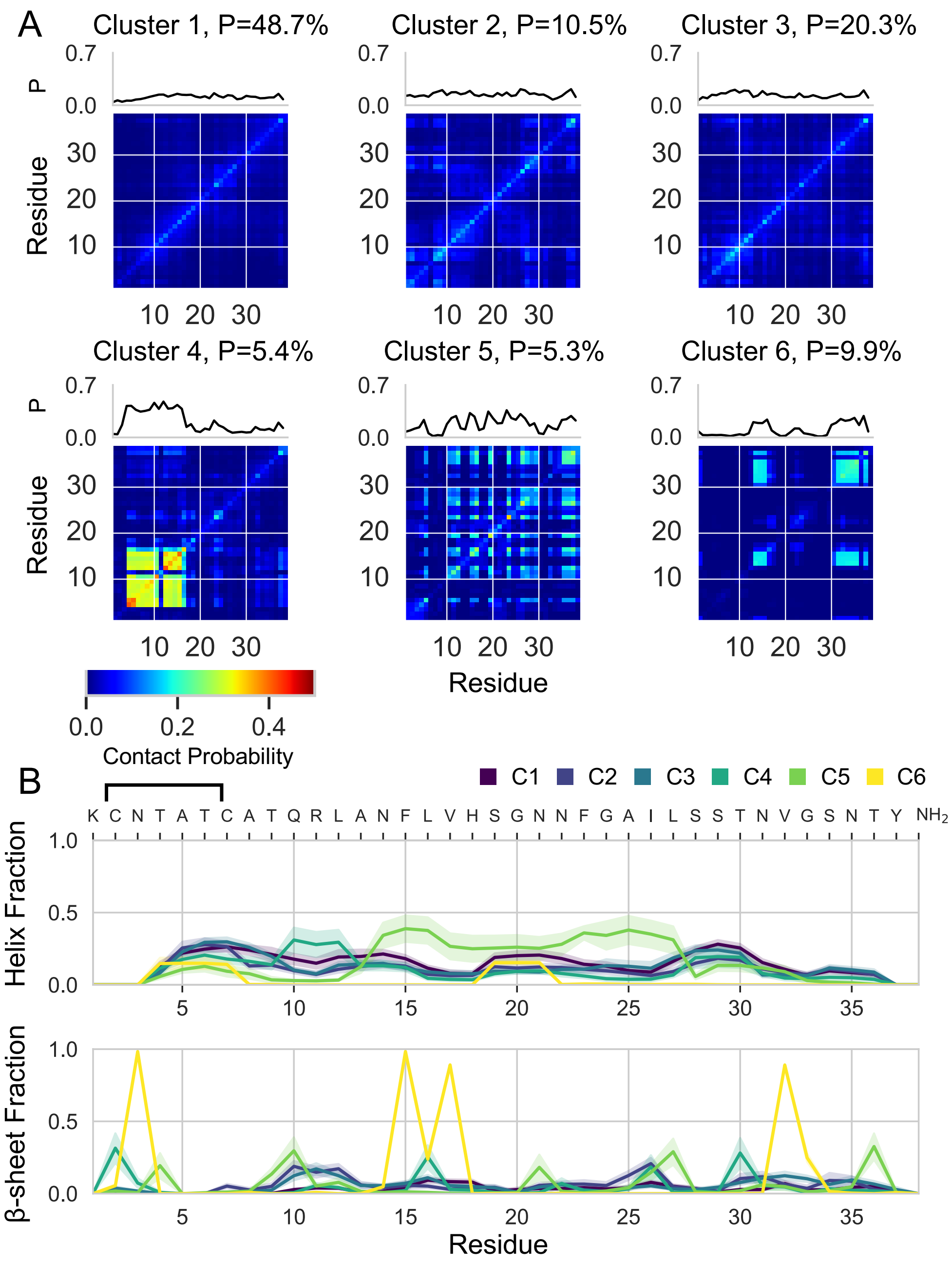


**Supplementary Figure 19. Ligand interactions and secondary structure populations of conformational states of S20G hIAPP in the presence of YX-A-1.** (A) The probability that each pair of hIAPP residues simultaneously forms a contact with YX-A-1 and (B) the secondary structure populations in conformational states identified by k-means clustering with k = 6 clusters from the 300K replica of a REST2 MD simulation of S20G hIAPP in the presence of YX-A-1. Secondary structure propensities were assigned by DSSP assignments and statistical error estimates from blocking are shown as shaded regions.

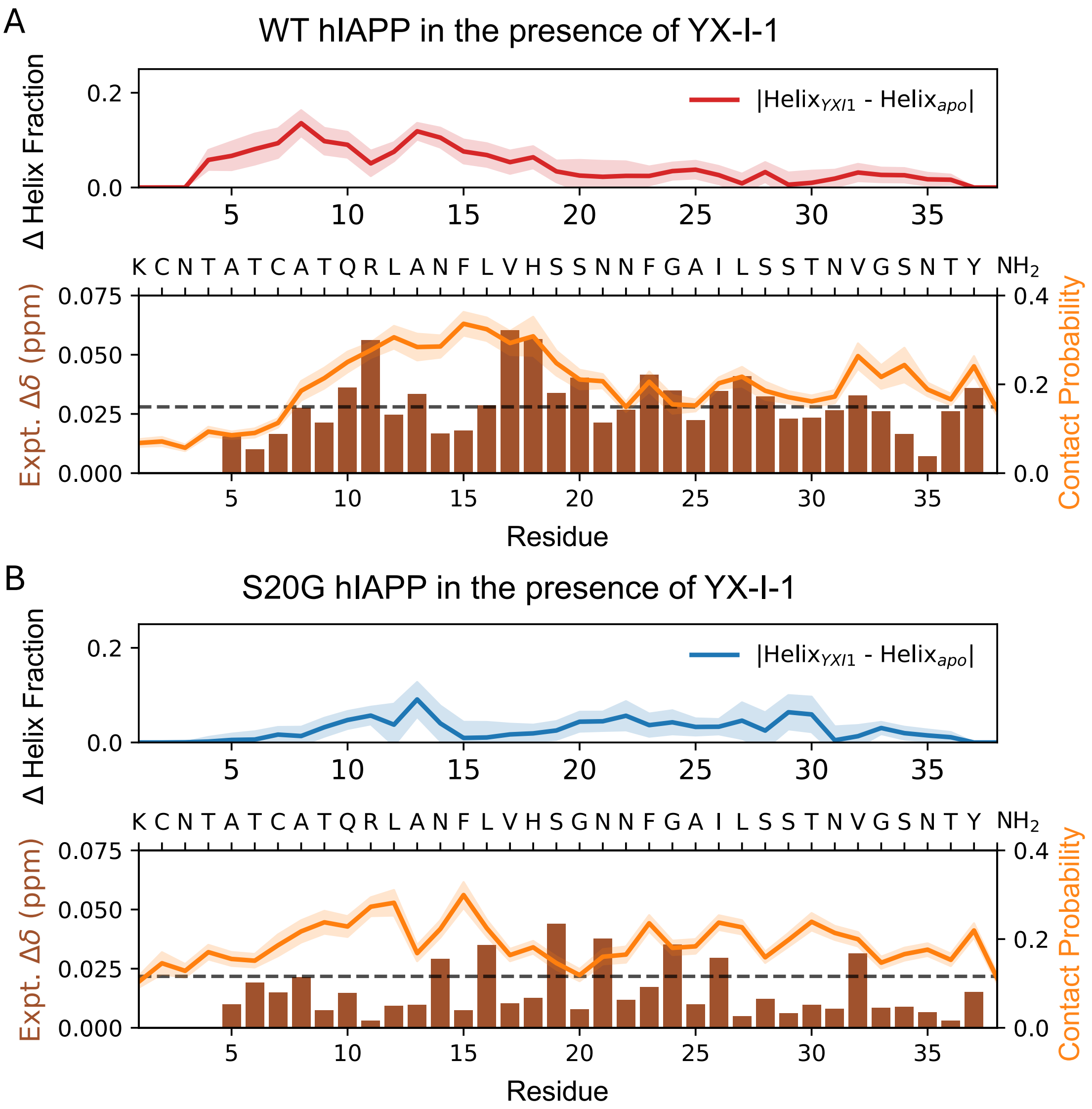


**Supplementary Figure 20. Comparison of changes in helix populations and populations of intermolecular protein-ligand contacts in ligand-binding simulations of YX-I-1 with experimental $^{1}$H-$^{15}$N NMR CSPs**. Top panels show absolute differences in simulated helix populations (Δ Helix Fraction) in apo and ligand-binding simulations. Bottom panels compare populations of intermolecular protein-ligand contacts (line plots, right axes) with previously reported $^{1}$H-$^{15}$N NMR CSPs (bar plot, left axes)[10]. Statistical error estimates from blocking shown as shaded regions. Experimental $^{1}$H-$^{15}$N NMR CSPs measured in a solution of 20 μM protein and 100μM YX-I-1 are reported as $\Delta\delta = \sqrt{[5 * (\Delta\delta H)^2 + (\Delta\delta N)^2\ ]}$. The dashed black line indicates the threshold of two standard deviations of reported $^{1}$H-$^{15}$N NMR CSPs.

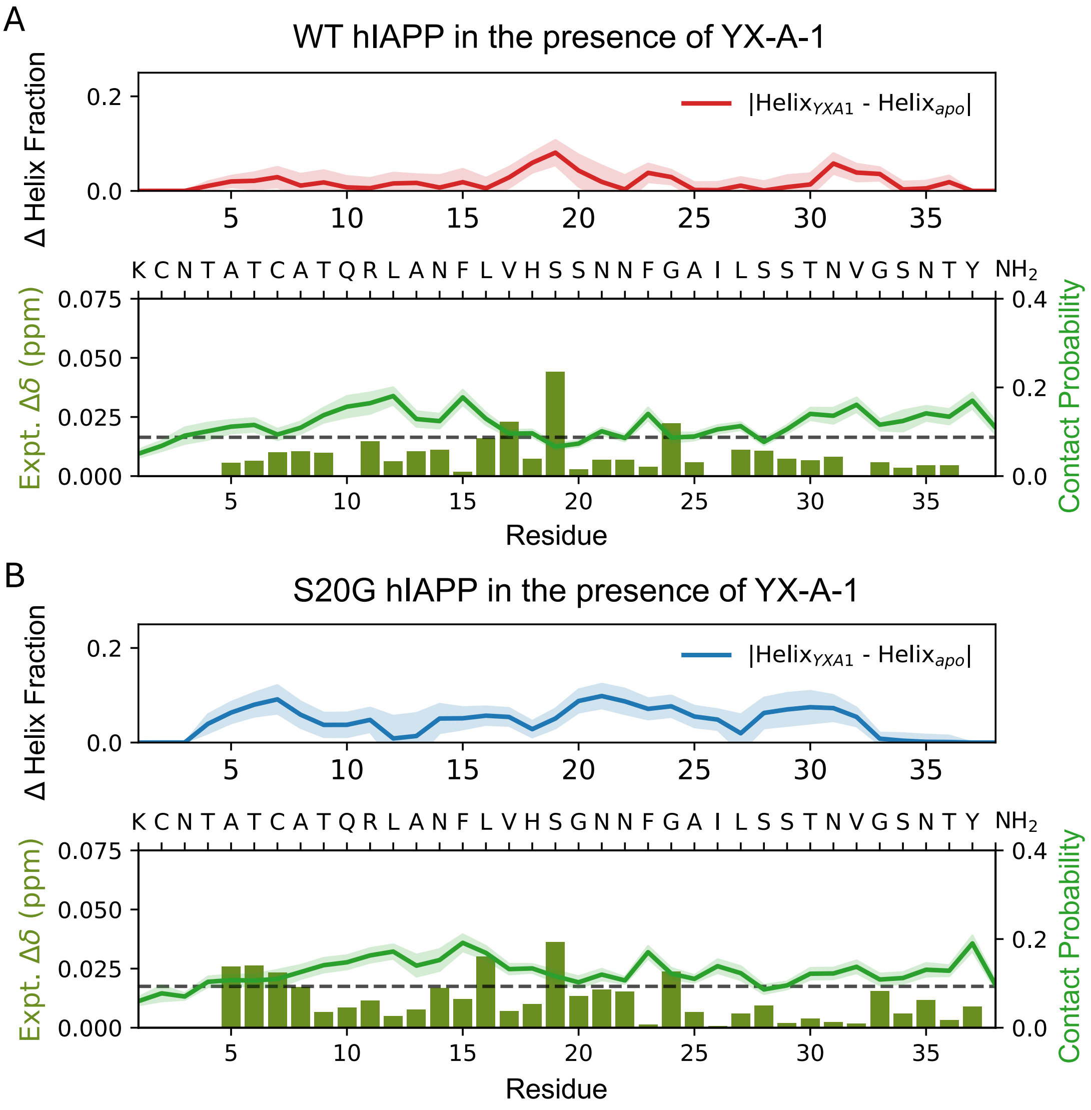


**Supplementary Figure 21. Comparison of changes in helix populations and populations of intermolecular protein-ligand contacts in ligand-binding simulations of YX-A-1 with experimental $^1$H-$^{15}$N NMR CSPs**. Top panels show absolute differences in simulated helix populations (Δ Helix Fraction) in apo and ligand-binding simulations. Bottom panels compare populations of intermolecular protein-ligand contacts (line plots, right axes) with previously reported $^1$H-$^{15}$N NMR CSPs (bar plot, left axes)[10]. Statistical error estimates from blocking shown as shaded regions. Experimental $^1$H-$^{15}$N NMR CSPs measured in a solution of 20 μM protein and 20 μM YX-A-1 are reported as $\Delta\delta = \sqrt{[5 * (\Delta\delta H)^2 + (\Delta\delta N)^2\ ]}$. The dashed black line indicates the threshold of two standard deviations of reported $^1$H-$^{15}$N NMR CSPs.

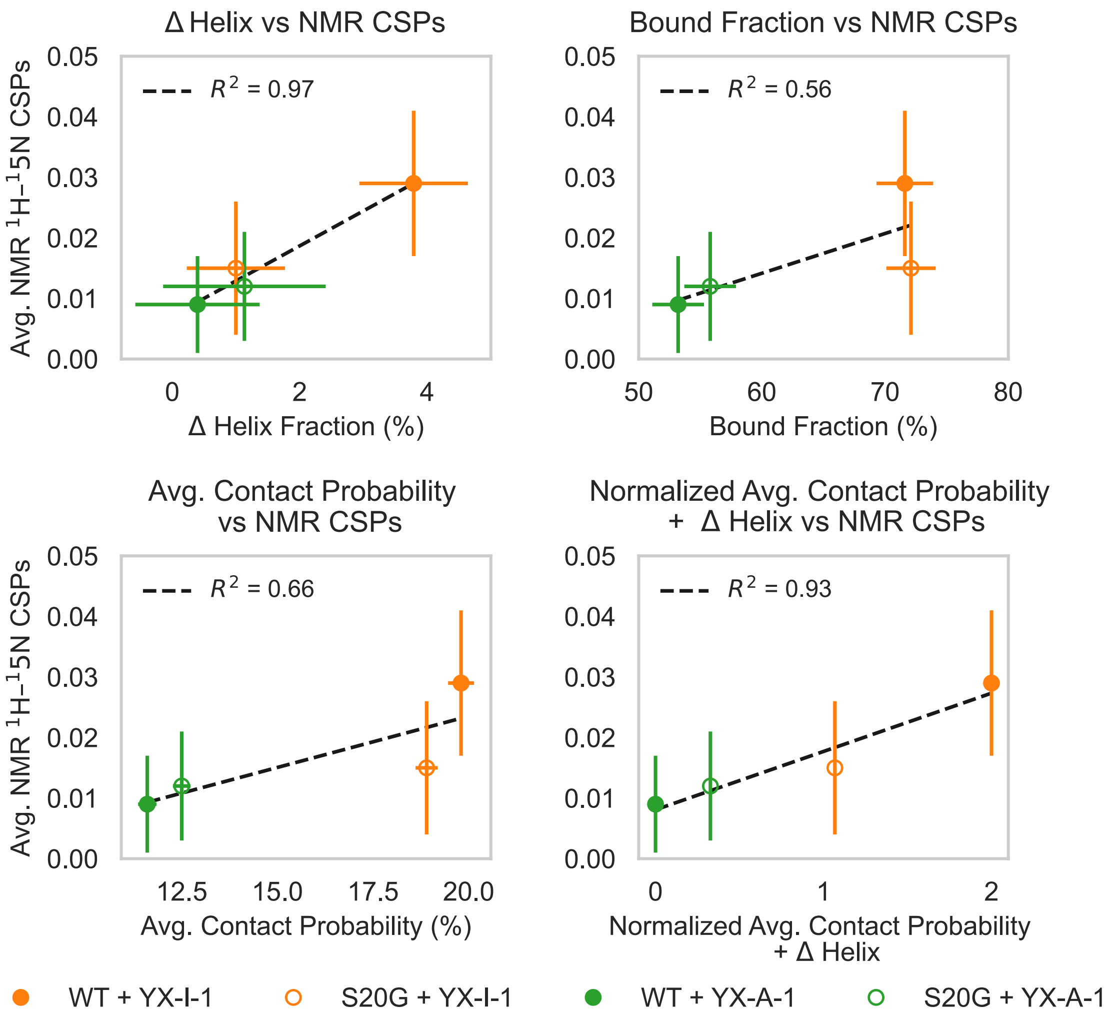


**Supplementary Figure 22. Correlation of average NMR CSP magnitudes with properties of ligand-binding simulations.** (A) Average absolute change of helix fraction (Δ Helix, relative to apo ensembles) vs average $^1$H-$^{15}$N NMR CSPs ($R^2$=0.97). Errors in Δ helix are average per-residue blocking errors. (B) Average bound fraction vs average $^1$H-$^{15}$N NMR CSPs ($R^2$=0.56). Errors in bound fractions were calculated from blocking analysis. (C) Average per-residue contact probability vs average $^1$H-$^{15}$N NMR CSPs ($R^2$=0.66). Errors in contact probabilities were obtained from blocking. (D) Combined normalized values of average per-residue contact probability and Δ Helix vs average $^1$H-$^{15}$N NMR CSPs ($R^2$=0.93). Contact probabilities and Δ Helix values were each normalized to 1 and summed (range 0–2). Errors of average $^1$H-$^{15}$N NMR CSPs correspond to one standard deviation. Linear regression fits are shown as dashed black lines. Symbols: filled orange = WT + YX-I-1; open orange = S20G + YX-I-1; filled green = WT + YX-A-1; open green = S20G + YX-A-1.

A

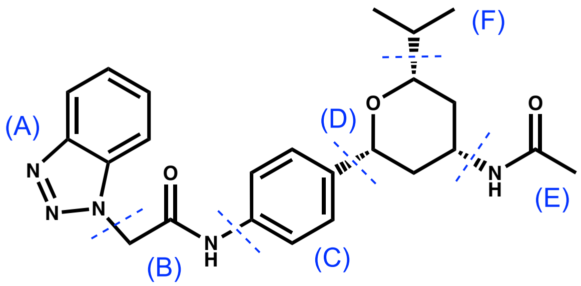


YX-I-1 Interaction Probability by Chemical Group

| | Benzotriazole (A) | Amide linker (B) | Phenyl (C) | Tetrahydro-pyran (D) | Terminal amide (E) | Isopropyl (F) |
|---|---|---|---|---|---|---|
| WT | 65.2 ± 2.5 | 65.6 ± 2.5 | 64.0 ± 2.5 | 60.5 ± 2.4 | 45.8 ± 1.7 | 48.5 ± 1.8 |
| S20G | 65.1 ± 2.2 | 65.2 ± 2.2 | 63.3 ± 2.3 | 57.0 ± 2.1 | 43.2 ± 1.4 | 43.2 ± 2.0 |

B

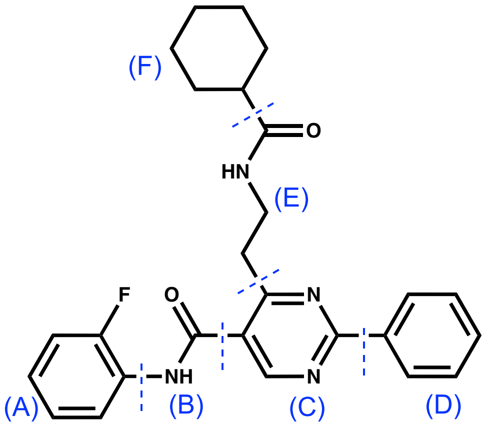


YX-A-1 Interaction Probability by Chemical Group

| | Fluorobenz-yl (A) | Amide linker (B) | Pyrimidine (C) | Phenyl (D) | Alkyl amide (E) | Cyclo-hexane (F) |
|---|---|---|---|---|---|---|
| WT | 41.6 ± 1.5 | 42.4 ± 2.1 | 45.1 ± 2.1 | 42.6 ± 2.0 | 43.6 ± 2.0 | 34.3 ± 1.3 |
| S20G | 42.7 ± 2.0 | 43.5 ± 2.0 | 47.7 ± 2.1 | 46.1 ± 2.2 | 46.1 ± 2.0 | 35.5 ± 1.8 |

**Supplementary Figure 23. Intermolecular contact populations of chemical groups of YX-I-1 and YX-A-1 in ligand-binding simulations.** The chemical structures of YX-I-1 (A) and YX-A-1 (B) are divided into six chemical groups by blue dashed lines. Tables list the fraction of frames in which at least one atom of a given group is within 5 Å of protein atom.

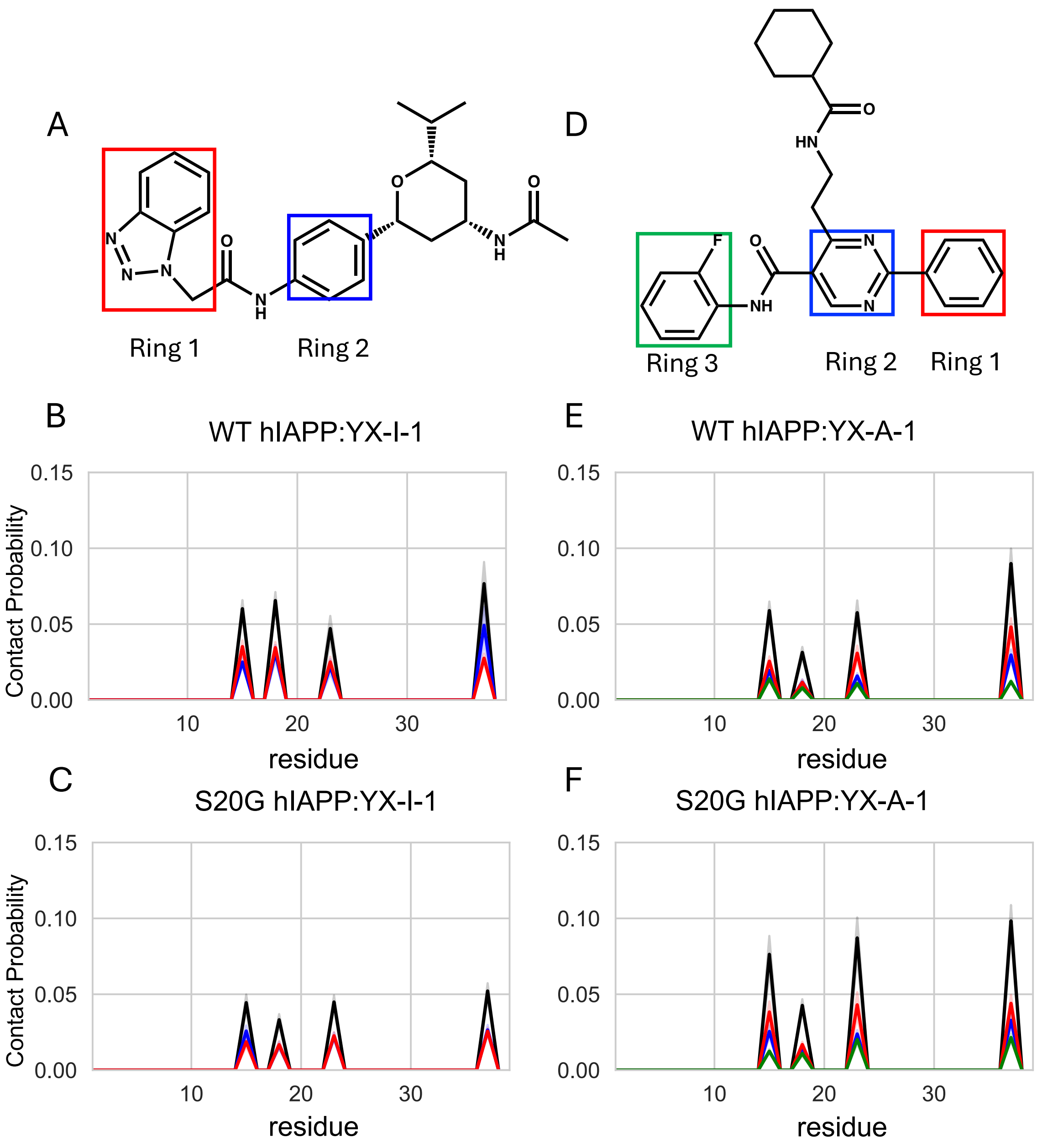


**Supplementary Figure 24. Per-residue aromatic stacking interactions in hIAPP binding simulations.** Chemical structures of YX-I-1 (A) and YX-A-1 (D) are shown with colored boxes highlighting aromatic rings. Per-residue total and ligand ring-specific stacking interactions are shown for binding simulations of WT hIAPP (C) and S20G hIAPP (D) with YX-I-1, and WT hIAPP (E) and S20G hIAPP (F) with YX-A-1. Black lines indicate total populations of stacking interactions, while colored lines correspond to populations of stacking interactions with individual rings.

### References


(1) Wang, L.; Friesner, R. A.; Berne, B. J. Replica Exchange with Solute Scaling: A More Efficient Version of Replica Exchange with Solute Tempering (REST2). *J. Phys. Chem. B* **2011**, *115* (30), 9431–9438. https://doi.org/10.1021/jp204407d.

(2) Bussi, G. Hamiltonian Replica Exchange in GROMACS: A Flexible Implementation. *Mol. Phys.* **2014**, *112* (3–4), 379–384. https://doi.org/10.1080/00268976.2013.824126.

(3) Sugita, Y.; Okamoto, Y. Replica-Exchange Molecular Dynamics Method for Protein Folding. *Chem. Phys. Lett.* **1999**, *314* (1), 141–151. https://doi.org/10.1016/S0009-2614(99)01123-9.

(4) Pietrucci, F.; Laio, A. A Collective Variable for the Efficient Exploration of Protein Beta-Sheet Structures: Application to SH3 and GB1. *J. Chem. Theory Comput.* **2009**, *5* (9), 2197–2201. https://doi.org/10.1021/ct900202f.

(5) Scalvini, B.; Sheikhhassani, V.; van de Brug, N.; Heling, L. W. H. J.; Schmit, J. D.; Mashaghi, A. Circuit Topology Approach for the Comparative Analysis of Intrinsically Disordered Proteins. *J. Chem. Inf. Model.* **2023**, *63* (8), 2586–2602. https://doi.org/10.1021/acs.jcim.3c00391.

(6) Ross, D. A.; Lim, J.; Lin, R.-S.; Yang, M.-H. Incremental Learning for Robust Visual Tracking. *Int. J. Comput. Vis.* **2008**, *77* (1), 125–141. https://doi.org/10.1007/s11263-007-0075-7.

(7) Pedregosa, F.; Varoquaux, G.; Gramfort, A.; Michel, V.; Thirion, B.; Grisel, O.; Blondel, M.; Prettenhofer, P.; Weiss, R.; Dubourg, V.; Vanderplas, J.; Passos, A.; Cournapeau, D.; Brucher, M.; Perrot, M.; Duchesnay, É. Scikit-Learn: Machine Learning in Python. *J. Mach. Learn. Res.* **2011**, *12* (85), 2825–2830.

(8) Alsabti, K.; Ranka, S.; Singh, V. An Efficient K-Means Clustering Algorithm. *Electr. Eng. Comput. Sci. - Scholarsh.* **1997**.

(9) Nielsen, J. T.; Mulder, F. A. A. CheSPI: Chemical Shift Secondary Structure Population Inference. *J. Biomol. NMR* **2021**, *75* (6), 273–291. https://doi.org/10.1007/s10858-021-00374-w.

(10) Shen, Y.; Bax, A. SPARTA+: A Modest Improvement in Empirical NMR Chemical Shift Prediction by Means of an Artificial Neural Network. *J. Biomol. NMR* **2010**, *48* (1), 13–22. https://doi.org/10.1007/s10858-010-9433-9.
